# Targeted high-resolution chromosome conformation capture at genome-wide scale

**DOI:** 10.1101/2020.03.02.953745

**Authors:** Damien J. Downes, Matthew E. Gosden, Jelena Telenius, Stephanie J. Carpenter, Lea Nussbaum, Sara De Ornellas, Martin Sergeant, Chris Q. Eijsbouts, Ron Schwessinger, Jon Kerry, Nigel Roberts, Arun Shivalingam, Afaf El-Sagheer, A. Marieke Oudelaar, Tom Brown, Veronica J. Buckle, James O.J. Davies, Jim R. Hughes

**Affiliations:** MRC Molecular Haematology Unit, MRC Weatherall Institute of Molecular Medicine, University of Oxford, Oxford, UK; MRC WIMM Centre for Computational Biology, MRC Weatherall Institute of Molecular Medicine, University of Oxford, Oxford, UK; Chemistry Research Laboratory, Department of Chemistry, University of Oxford, Oxford, UK; Big Data Institute, Li Ka Shing Centre for Health Information and Discovery, University of Oxford, Oxford, UK; Wellcome Centre for Human Genetics, Nuffield Department of Medicine, University of Oxford, Oxford, UK

**Keywords:** Gene regulation, chromosome conformation capture, 3C, genome structure

## Abstract

Chromosome conformation capture (3C) provides an adaptable tool for studying diverse biological questions. Current 3C methods provide either low-resolution interaction profiles across the entire genome, or high-resolution interaction profiles at up to several hundred loci. All 3C methods are affected to varying degrees by inefficiency, bias and noise. As such, generation of reproducible high-resolution interaction profiles has not been achieved at scale. To overcome this barrier, we systematically tested and improved upon current methods. We show that isolation of 3C libraries from intact nuclei, as well as shortening and titration of enrichment oligonucleotides used in high-resolution methods reduces noise and increases on-target sequencing. We combined these technical modifications into a new method Nuclear-Titrated (NuTi) Capture-C, which provides a >3-fold increase in informative sequencing content over current Capture-C protocols. Using NuTi Capture-C we target 8,061 promoters in triplicate, demonstrating that this method generates reproducible high-resolution genome-wide 3C interaction profiles at scale.

## Introduction

Chromosome conformation capture (3C) has emerged as the leading tool for studying the DNA folding associated with gene regulation and genome organization^1, 2^. 3C methods measure the proximity of DNA elements through restriction enzyme digestion and ligation; sequencing of the resultant chimeric fragments produces a population-based interaction frequency as the output. The resolution achieved by 3C comes from the choice of restriction enzyme, the depth of sequencing, and whether or not targeted enrichment is performed. Currently, 3C methods can be categorized into two broad classes depending on their resolution.

Low-resolution 3C methods, such as Hi-C^3^ and its derivatives, tend to use a 6-bp cutting enzyme to generate genome-wide interaction maps, with the standard experiment generating 10-50 kb resolution^2^. Higher-quality profiles can be achieved through combinations of hugely increased sequencing, use of a 4-bp cutter, targeted enrichment (e.g. Capture Hi-C^4^ [CHi-C], often called Promoter Capture Hi-C), and increased cell numbers. The prohibitive costs mean that such datasets rarely include sufficient number of replicates (triplicates) for statistical analysis and are not applicable to rare primary cell types due to the requirement for high cell numbers. Conversely, sub-kilobase resolution can be achieved by methods which enrich for target loci in 4-base cutter libraries; e.g. Capture-C^5^, 4C-seq^6, 7^, and their derivatives. The current best high-resolution 3C method for sensitivity is NG Capture-C, with 10,000-100,000+ unique interacting reporter reads per viewpoint^2, 8^. NG Capture-C achieves its high resolution and sensitivity using biotinylated oligonucleotide pull down of target loci from 3C material, generated with a 4-bp cutter. The use of sequential enrichment, or “double capture”, results in 30-50% on-target sequencing, an 160-fold increase over the initial Capture-C method^5, 8^.

High-resolution 3C comes at the expense of the number of viewpoints that can be practically included in a single experiment. This is due to the roughly 16-fold increase in complexity when generating a 3C library with a 4-bp cutter compared to a restriction enzyme with a 6-bp motif. The need to robustly sample these much more complex libraries has so far limited NG Capture-C to hundreds of viewpoints, generally performed in triplicate for statistical analysis. However, a large increase in the specificity of enrichment and the minimalization of off-target and technical noise would practically translate into the feasibility of much larger viewpoint designs using high-resolution methods.

To dramatically increase the capacity of NG Capture-C, we have systematically optimized multiple aspects of the protocol. 3C libraries in general are prone to technically induced noise, which results in an increased frequency of non-informative *trans* reporters^9^. These spurious reporters represent experimental background and so do not informatively add to the interaction profiles, but do increase the required amount of sequencing. Consistent with previous work^10^, we show that the 3C libraries can be separated into nuclear and non-nuclear fractions with differing levels of information content. By optimising the 3C method to enrich for and isolate intact nuclei after ligation we show a 30% increase in informative content.

We next tested the effect of probe length and concentration on enrichment. Reducing oligonucleotide probes from 120 to 50 bp resulted in a 5% increase in reads containing *Dpn*II sites. Additionally, titration of probe concentration resulted in a significant increase in capture specificity, and when combined with double capture resulted in up to 98% on-target capture; a 100-200% improvement over double capture alone. We have combined these optimisations, along with improvements to minimize losses during 3C DNA extraction and indexing^9^ to generate a modified protocol: Nuclear-Titrated (NuTi) Capture-C.

The two seminal descriptions of targeted genome-wide 3C landscapes were carried out in human CD34^+^ and GM12787 cells^4^, and in mouse embryonic stem cells (ESC) and fetal liver cultured erythroid cells^11^ using CHi-C with the low-resolution *Hind*III 6-base cutter and targeting every gene through its longest annotated promoter. To demonstrate that NuTi Capture-C can be used to improve upon these efforts we first used RNA-seq, DNaseI-seq and ChIP-seq from mouse ter119^+^ erythroid cells to identify 7,870 active promoters and 181 inactive promoters for targeting. We performed NuTi Capture-C in triplicate from primary mouse erythroid cells, detecting over 1,000 unique ligation events for 93.6% of targets (6,732 of 7,195 *Dpn*II fragments). Using a Bayesian modelling approach to interaction calling^12^ we were able to identify 472,270 promoter-interacting fragments across the genome, including 12,316 finely mapped promoter/enhancer interactions. When compared to erythroid interactions found by CHi-C, NuTi Capture-C had a higher enrichment for active chromatin marks and greater specificity at identifying promoter-enhancer interactions. Therefore, with the application of NuTi Capture-C researchers will be able to map the regulatory landscapes of thousands of loci, *en masse* and at high-resolution.

## RESULTS

### Nuclear isolation reduces the frequency of spurious ligation

The quality of 3C libraries, as measured by changes in *cis*-to-*trans* ligation frequencies, can be drastically affected by technical noise^9, 10^. Therefore, it is important to generate high-quality 3C libraries which minimize this noise. Previous work has shown that a portion of nuclei remain intact during 3C digestion/ligation and intact nuclei contain the informative 3C DNA^10^. Most 3C methods are performed using the *in situ*^13^ protocol which assumes a majority of ligation events occur within intact nuclei – rather than between DNA released from nuclei through either diffusion or nuclear rupture. Any DNA that does escape from nuclei can generate technical noise through inter-nuclear ligation, seen as higher “*trans*” interactions. To test the extent to which DNA is released from nuclei during digestion and ligation in *in situ* 3C libraries we used centrifugation to separate the post-ligation 3C library milieu into an insoluble nuclear fraction and a soluble DNA fraction (Fig. 1a). We found ∼25% of DNA could be found in the un-pelleted supernatant using the standard *in situ* 3C method (Fig. 1b). NG Capture-C of both the standard *in situ* 3C milieu and partitioned fractions showed a higher *cis*-ligation frequency in nuclear material, and a higher *trans*-ligation frequency in the soluble fraction (Fig. 1c). A comparison of interactions at the *Hba-1/2*, *Hbb-b1/2* and *Slc25a37* loci showed maintenance of the general interaction profile between different fractions, however there was an increase in the proximal signal in the soluble fraction (Fig. 1d, Supp. Fig. 1) at the expense of informative long–range interactions.

**Fig. 1.**
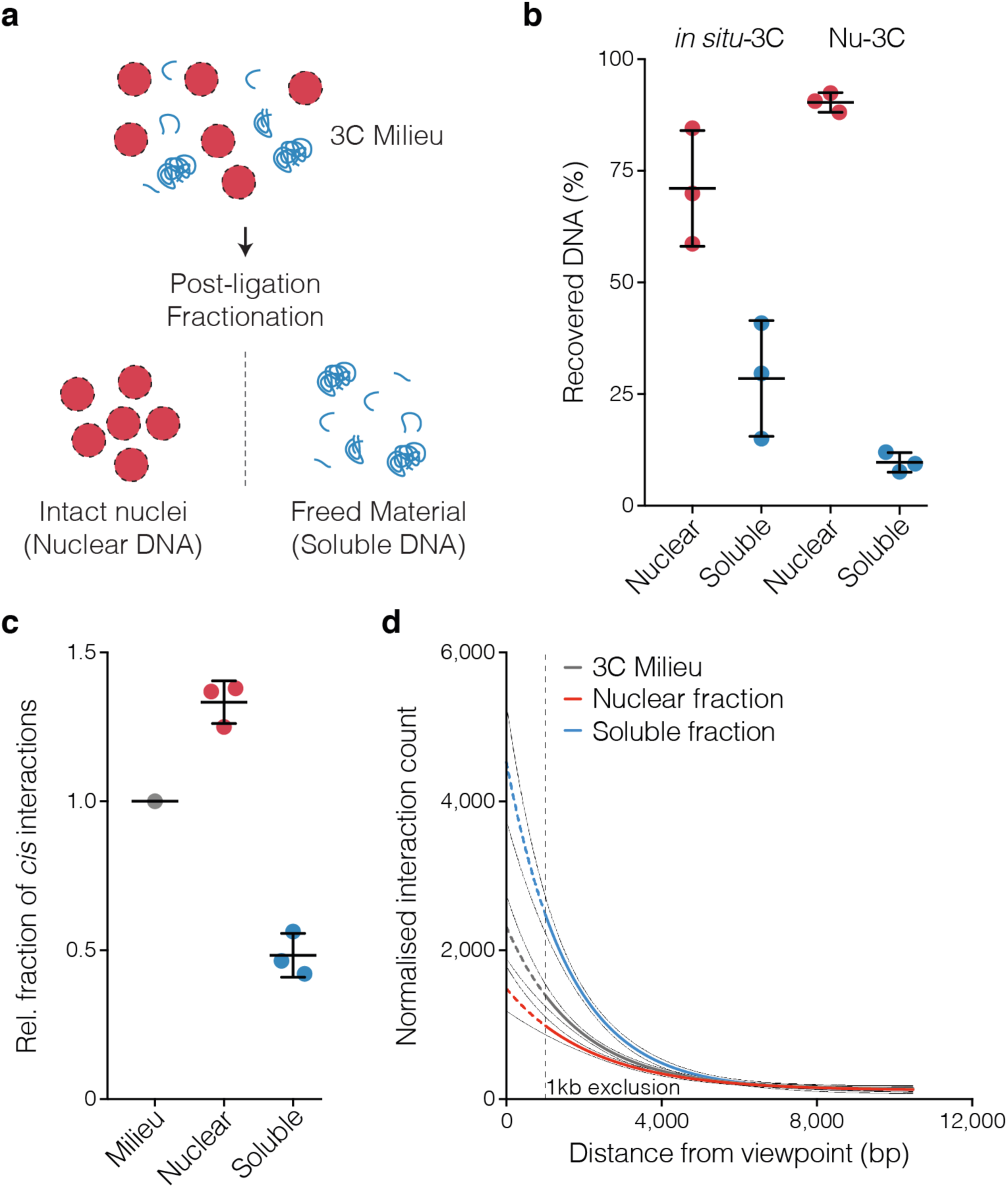
The 3C library milieu can be separated into high- and low-quality fractions. **a,** During digestion and ligation nuclei can shear leading to free soluble chromatin. Intact nuclei can be separated from freed material by centrifugation. **b,** Percent of total DNA recovered in the two fractions using standard *in situ*-3C and a modified Nuclear 3C (Nu-3C) approach. **c,** Relative fraction of *cis* interactions for libraries generated simultaneously using *in situ*-3C with and without fractionation. **d,** Average number of interactions within 10.5 kb of the *Hba-1/2*, *Hbb-b1/2* and *Slc25a37* capture viewpoints from *in situ*-3C fractions. Bars show mean and one standard deviation.

To measure the extent of spurious inter-nuclear ligation, we generated *in situ* 3C libraries from an admixture of human and mouse erythroid cells (Fig. 2a). By mixing samples during the 3C process half of inter-nuclear ligations are detectable as mouse-to-human ligations, or chimeric inter-species fragments. Detectable chimeric ligations represented 10-15% of reporter containing fragments (Fig. 2b). For each 3C library an equivalent number of fragments would contain inter-nuclear but undetectable mouse-mouse or human-human ligations; therefore 20-30% of all *in situ* 3C ligations were inter-nuclear artefacts which lack biological relevance. This is consistent with ∼25% of *in situ* 3C DNA being found in the un-pelleted supernatant. This high rate of spurious ligation suggests data quality could be improved by maintenance nuclear integrity and isolation of intact nuclei after ligation – as opposed to before restriction endonuclease digestion. Analysis of these Nuclear 3C (Nu-3C) libraries showed that soluble 3C material was reduced to 10% of all DNA (Fig. 1c), and only 4% of ligations were chimeric inter-nuclear events, which resulted in a significant increase in *cis* interactions (Fig. 2b). Therefore Nu-3C libraries represent a higher quality starting product for quantifying biologically relevant interactions.

**Fig. 2.**
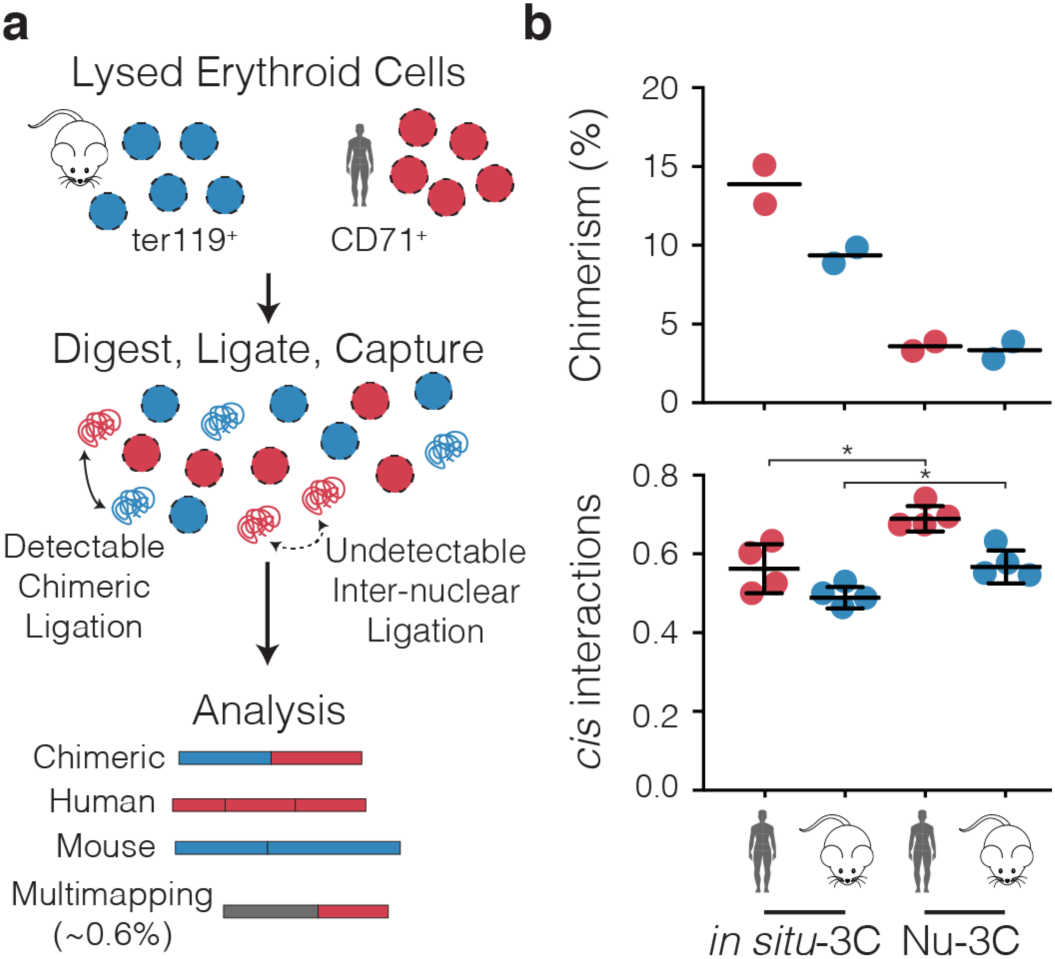
Detection of spurious ligation through inter-species ligation. **a,** Lysed erythroid cells from human and mouse were mixed in a 1:1 ratio prior to generation of 3C libraries. Ligation occurring between ruptured nuclei can be detected as inter-species chimeric DNA fragments after filtering for sequences that map to both genomes. **b,** Analysis of the level of inter-species chimeras and number of reported *cis* interactions when using standard *in situ*- 3C or modified Nuclear 3C (Nu-3C) at the *Hba-1/2* and *Slc25a37* promoters (n=2). Nu-3C reduced the amount of noise in libraries from spurious internuclear ligations. Bars show mean and one standard deviation. *p<0.05 from a Mann Witney test.

### Probe titration increases targeting efficiency

NG Capture-C was designed to capture target viewpoints with tens or hundreds of 120-bp biotinylated DNA oligonucleotides^8^; high enrichment is achieved through double capture. This method uses a commercial exome sequencing kit optimized to include several thousand oligonucleotides. To determine whether targeting efficiency is affected by oligonucleotide concentration we tested serial dilutions while targeting 11 loci in mouse erythroid cells and ESC. Lower probe concentrations resulted in reduced yields of DNA following single capture (Fig. 3a). Sequencing of captured material with each individual probe at a working concentration of 2.9 nM, produced 31.61% on-target sequencing (Stdev=2.00, n=4), similar to that of double capture without dilution^8^. When lower concentration probes were used in combination with double capture (Titrated Capture-C), 85-98% on-target sequencing was achieved; indicating the two optimizations are additive. When this combined method was applied to *Slc25a37* alone, a 97.70% on target sequencing was seen, equating to a 6.26-million-fold enrichment (see Methods). Increased on-target sequencing reduces the depth of sequencing required to generate the same number of informative reads. We *in silico* tested the number of raw reads required to generate high-quality profiles by down-sampling fastq files. When using probes targeting both ends of a viewpoint only 250,000 reads are required to exceed 30,000 unique interactions (Supp. Fig. 2). This depth of signal is 2.1 times better than the original NG Capture-C method^8^, and 11.6 times better than for an equivalent depth of sequencing for UMI-4C^14^.

**Fig. 3.**
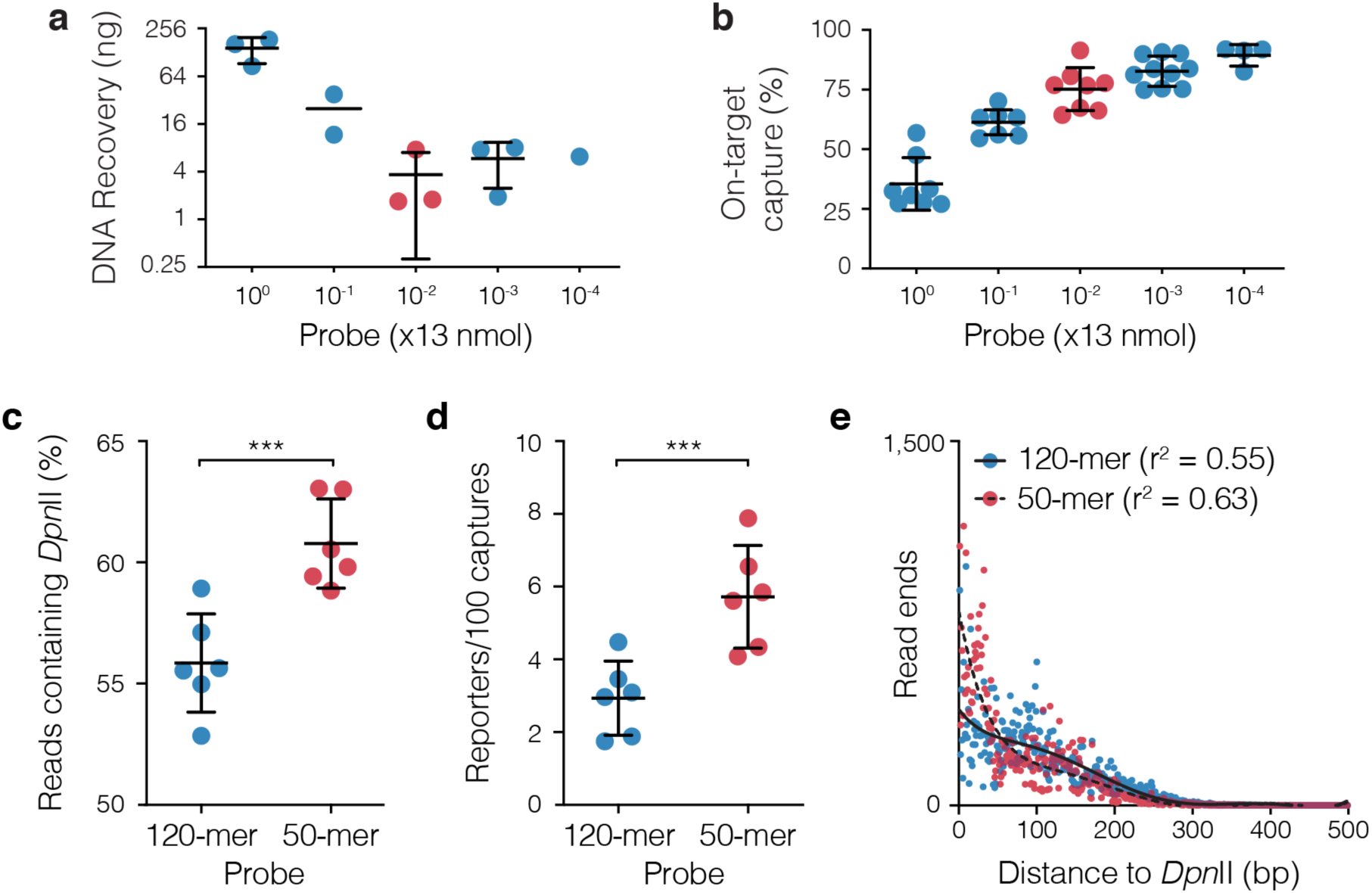
Probe concentration and length alter capture efficiency. Total yield of DNA recovered following single capture **(a)** and total number of mapped reads containing on-target capture sequence following double capture **(b)** when 11 probe pairs were serial diluted from 2.9 µM to 0.29 nM. For DNA recovery each dot is a multiplex capture with between 3 and 6 libraries. For On-target capture each dot is a separate 3C library. Bars show mean and one standard deviation. Percent of reads with a *Dpn*II site **(b)**, number of PCR duplicate filtered reporters per 100 mapped reads containing a reporter **(c)** following capture of six 3C libraries with 120-mer and 50-mer oligonucleotides. ***p=0.0001 using Mann-Witney rank sum test. Bars show mean and one standard deviation. **d,** Counts of read-ends generated by sonication breakpoints as the distance to the nearest end of the *Slc25a37* viewpoint. Each dot is average depth normalized count at each position for 100,000 mapped reads (n=12). Lines of best fit were generated as a sixth order polynomial with r^2^ shown in the legend.

A reduced read requirement represents a significant saving in the overall cost of Capture-C based experiments, which previously was a criticism of the method^13^. Another significant cost for NG Capture-C has been the 120-bp biotinylated oligonucleotides – though current pricing is significantly reduced. We performed capture with 50-bp oligonucleotides targeted to the well-characterized mouse globin and mitoferrin encoding genes. Shorter oligonucleotides generated reads with proportionally more *Dpn*II restriction sites and significantly more informative reads per captured fragment (Fig. 3c,d). Consistent with these findings, analysis of the ends of captured *Slc25a37* fragments showed sonication breakpoints tended to be closer to the captured *Dpn*II sites when using shorter oligonucleotides (Fig. 3e). This increase in informative capture events had no major changes to the local profiles of *Hba-1/2* and *Slc25a37*; with generally the same level of *cis* interactions and high reporter correlation between oligonucleotide lengths (Supp. Figs. 3,4). However, at *Hbb-b1/2* additional peaks of interaction were seen in both erythroid and ESC cells leading to reduced correlation between oligonucleotide lengths (Supp. Fig. 5). Analysis of the sequences underlying these peaks showed a higher proportion of sequence identity for the 50-bp oligonucleotides. Given the increased similarity and that these peaks were fragment specific, they are likely artefacts arising from off-target capture. Therefore, while short probes provide more informative capture, they can also generate interaction artefacts through reduced specificity in highly duplicated loci.

### Enrichment generates significant bias at co-targeted fragments

Ligation frequency is the core readout of 3C techniques; many approaches use targeted enrichment through either oligonucleotide pull down (NG Capture-C^8^, Capture Hi-C^4^), immunoprecipitation (HiChIP^15^, ChIA-PET^16^, ChIA-Drop^17^) or RNA enrichment (HiChIRP^18^) to generate this readout. The introduction of bias in 3C experiments by enriching at multiple sites (i.e. “co-targeting”) is widely acknowledged^5, 13^, but its magnitude has not been specifically reported. We first generated a mathematical model for enrichment-based bias (Supp. Note). Our model shows that bias will be variable from 1-to-20 fold, and affected by both the true interaction frequency of co-targeted fragments, and their relative enrichment efficiencies. To experimentally validate this model, we performed two captures at the well-characterized mouse *Hba-1/2* and *Hbb-b1/2* loci^19, 20^. In the first capture four promoters and three enhancers were targeted; in the second capture an additional 54 evenly spaced targets were included^21^. The addition of the nearby oligonucleotides led to a significant difference in interaction counts at the co-targeted fragments (Fig. 5a). The bias was confined specifically to the co-targeted fragments, and its magnitude was consistent with modelling. Moreover the level of bias depended on both the underlying signal and the viewpoint (Fig. 5b,c, Supp. Fig. 6a) – indicating our model is a good first order approximation of co-targeting bias. This specific bias is also seen in CHi-C^11^, such as at *Hba-1* in mouse erythroid cells (Supp. Fig. 6b). As bias from oligonucleotide pull-down is limited to targeted fragments, large high-resolution designs with thousands of viewpoints are possible, provided the correct data analysis is used to exclude bias.

**Fig. 4.**
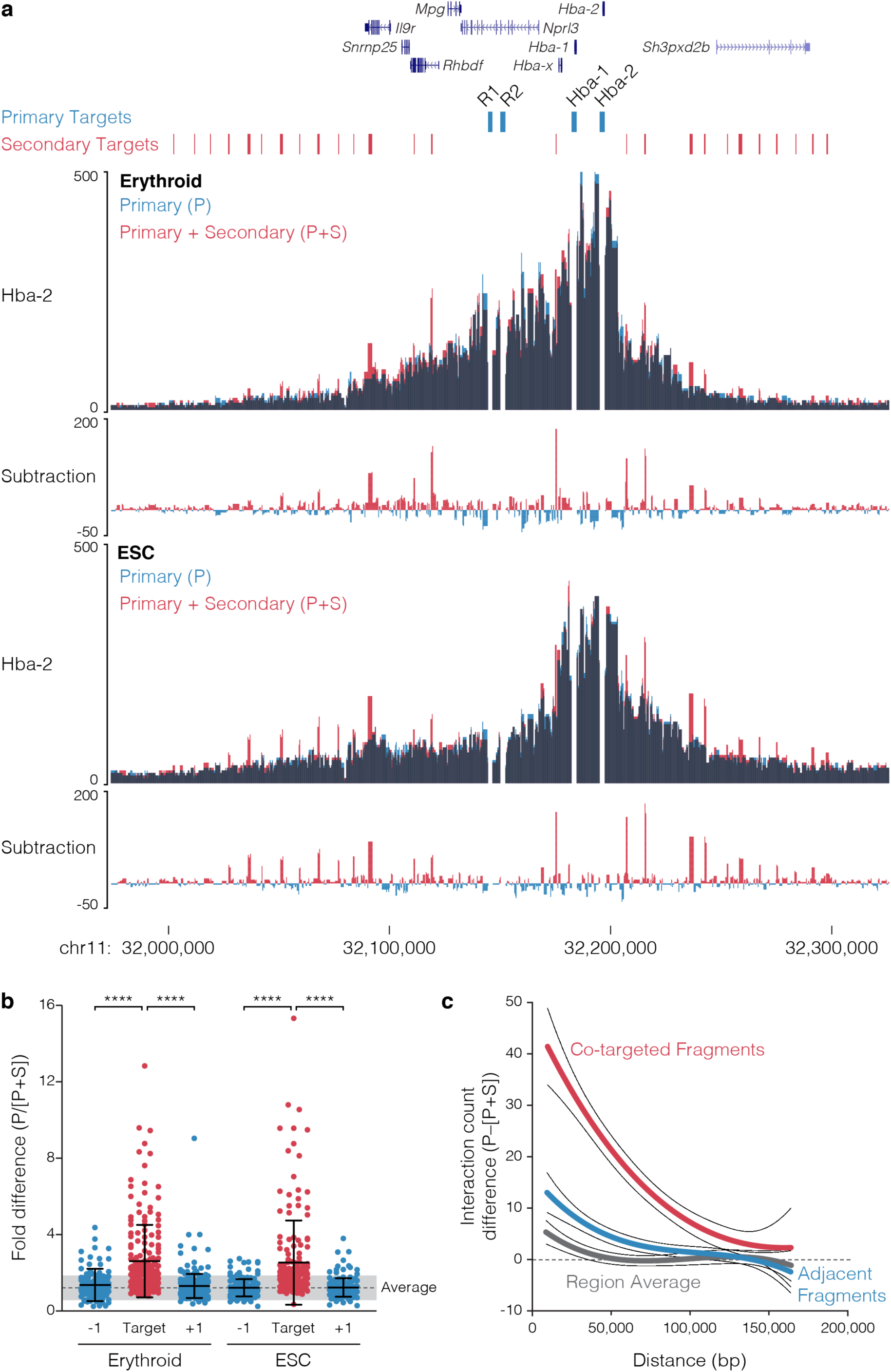
Co-targeting results in target specific bias. **a,** 3C libraries from mouse erythroid (n=3) and embryonic stem cells (ESC; n=3) were captured with either a pool of probes containing eight primary (P) viewpoints, or a pool of probes containing both the primary viewpoints and 54 additional, or secondary (S), viewpoints. Captured fragments were analyzed only for the primary viewpoints. Data is shown as an overlay for the Hba-2 capture viewpoint, with dark areas showing where signal overlaps. **b**, Comparison of the relative difference in interaction counts at co-targeted fragment and the adjacent fragments (±1). Average is shown for all fragments within 160 kb of the primary targets. **c,** Distance dependent difference in signal caused by co-targeting compared with adjacent fragments and the region average. ****p<0.0001 using Mann-Witney rank sum test. Bars show mean and one standard deviation.

**Fig. 5.**
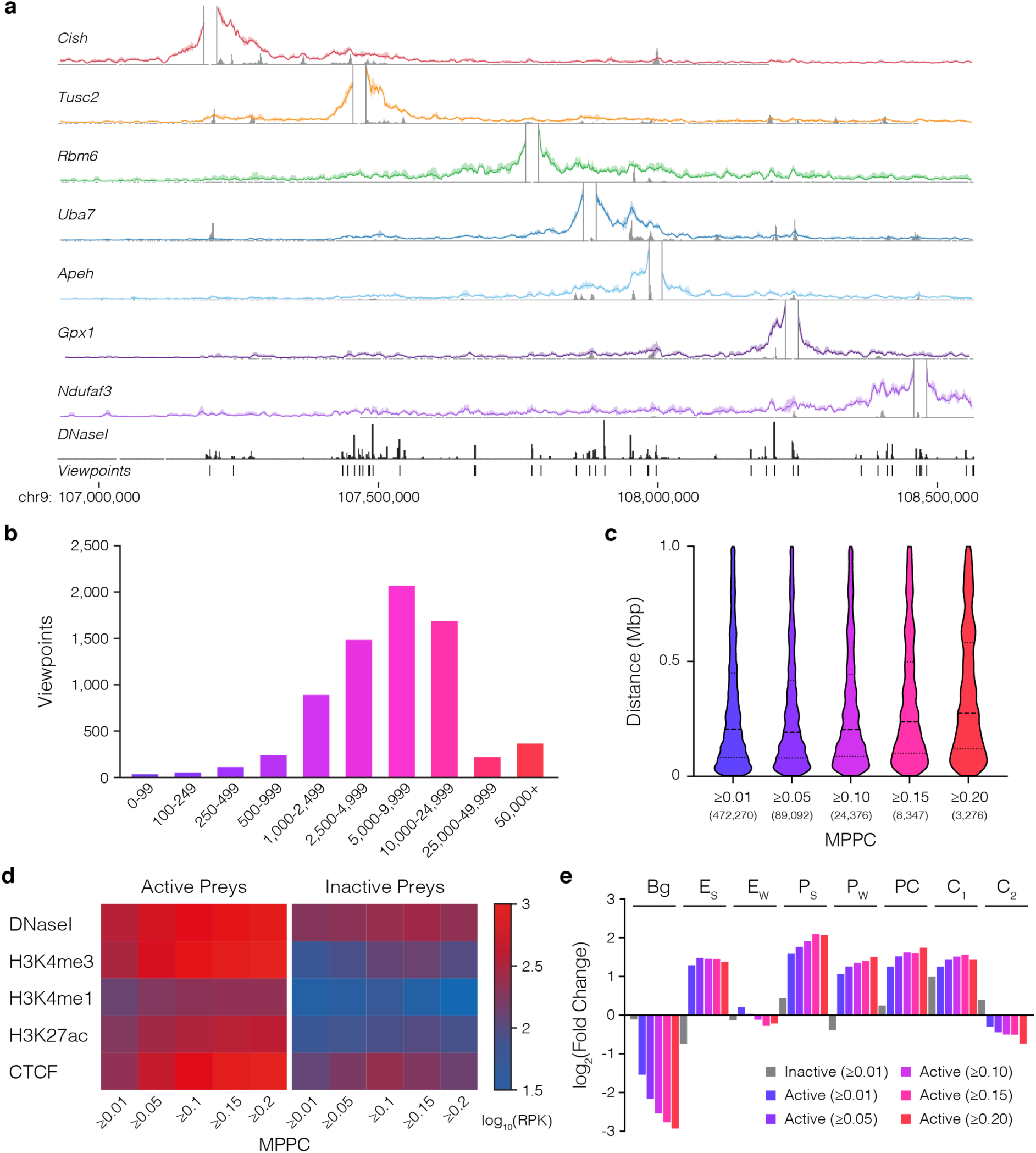
High resolution capture of 8,026 promoters. **a)** Windowed 3C interactions over 1.5Mb for seven NuTi Capture-C viewpoints with peaky Marginal Posterior Probability of Contact (MPPC) scores in grey (mm9: chr9:106926158-108566246) **b)** Histogram of total number of unique reporters identified per viewpoint from triplicate 3C libraries. **c)** Violin plots of the distance between the midpoints of captured promoters and peaky identified interacting fragments with increasing MPPC thresholds. **d)** Chromatin signal for interacting fragments (preys) of increasing MPPC identified by capturing either active or inactive promoters **e)** Enrichment of GenoSTAN annotations for interacting fragments with increasing MPPC. Bg: background, E_S_: Enhancer (Strong H3K27ac), E_W_: Enhancer (Weak H3K27ac), P_S_: Promoter (Strong H3K27ac), P_W_: Promoter (Weak H3K27ac), PC: Promoter/CTCF, C_1_: CTCF near Promoter/Enhancer, C_2_: CTCF.

### Erythroid interaction maps for 8,061 promoters

Although NG Capture-C has been employed widely for reproducible high-resolution characterization of local chromatin interactions^2, 8^, and CHi-C is used to generate low-resolution profiles at thousands of loci^4, 11^, no method has yet been implemented to generate high-resolution 3C maps for thousands of loci in triplicate. By combining higher quality Nu-3C libraries, low-cell optimizations^9, 22^, increased efficiency targeting through Titrated Capture-C, and a reduction in PCR cycles, our new method, Nuclear-Titrated (NuTi) Capture-C (Supp. Fig. 7), could feasibly be applied to generate reproducible high-resolution data in both small and genome-scale experiments. To this end we used DNaseI-seq and ChIP-seq for H3K27ac, H3Kme1, H3Kme3 signals from mouse ter119^+^ erythroid cells^19, 23^ to annotate tissue-specific transcription start sites of protein coding genes, identifying 7,870 active promoters for targeting (Supp. Fig. 8). We also included in the design a further 191 inactive control promoters, in total covering 7,195 *Dpn*II fragments. Using this design, NuTi Capture-C was performed in triplicate for ter119^+^ erythroid cells and sequenced to an average of 150-300k read-pairs per viewpoint (Fig. 5a). We identified 140.8M unique ligation events with over 1,000 unique *cis*-ligation events for 93.5% of targets (n=6,730; Fig. 5b). We first compared the profiles of the well-characterized *Hba*-1/2, *Hbb-b1/2*, *Slc25a37* loci between small- and genome-scale capture designs (Supp. Fig. 9), finding good correlation between experiments (Pearson r^2^: 0.75-0.87). Interestingly, viewpoints shorter than 300 bp tended to have higher levels of *trans* interactions despite nuclear isolation (Supp. Fig. 10a,b). Analysis of non-nuclear DNA from *Hind*III and *Dpn*II 3C digestion found higher amounts of DNA from the 4-bp cutter (Supp. Fig. 10c,d). This suggests short fragments may either evade crosslinking, or be freed as small, diffusible fragments by digestion – resulting in the observed differences in *cis*-to-*trans* frequencies. Therefore, a minimum fragment length could be considered during viewpoint selection.

To identify significant distal interactions for each promoter we employed Bayesian modelling with peaky^12^ (Fig. 5a). Peaky identified 473,270 interacting fragment pairs (Marginal Posterior Probability of Contact [MPPC] ≥0.01) covering 75.8% of targeted viewpoints (n=5,451) and distributed between 2,500 bp and 1 Mb from the midpoint of the target. Identified fragments had strong enrichment for chromatin marks associated with active promoters and enhancers, with stronger enrichment seen for fragments with higher MPPC scores (Fig. 5c,d). To determine the identity of interacting regions we annotated 68,723 erythroid open-chromatin sites into eight classes using the GenoSTAN Hidden Markov Model^24^ (Supp. Fig. 11a,b). By intersecting significantly interacting fragments with these annotations we found 22,767 pairwise element interactions, accounted for by 56.7% (n=4,082) of targeted genes (Supp. Fig. 10c,d). When comparing the types of elements active promoters interact with, we found specific enrichment for both promoters and enhancers (Fig. 5d), with each gene interacting with an average of 2.8 promoters and 2.3 enhancers.

As NuTi Capture-C represents a technical advance in resolution over CHi-C, we directly compared our results with published CHi-C results in murine erythroid cells^11^. In general, the high-resolution method produced more fine-grained interaction profiles for promoters, including for genes in adjacent regulatory domains (Fig. 6), and shared regulatory domains (Supp. Figs. 12-15), even when resolution is reduced with a 5 kb window. The smaller fragment size also meant fewer fragments were affected by co-targeting bias, which provided more informative profiles in gene dense regions and allowed targeting of alternate promoters (Supp. Figs. 12,15-17). The interaction calls identified using NuTi Capture-C also appear more specific to functional elements than the broad regulatory domain calls of CHi-C (Supp. Figs. 12-28). We systematically compared our interaction calls with reported interaction calls. While we also found promoter-promoter interactions, consistent with the idea of promoter-hubs as reported by CHi-C^11^, we find many fewer constituent promoters (3.8 versus >20), likely due to the removal of co-targeting bias from NuTi Capture-C analysis. Next, where overlapping viewpoints were captured by the two methods, we found a higher level of active chromatin marks at interacting fragments identified with NuTi Capture-C (Supp. Fig. 29a). We also compared the types of annotated elements identified within interacting fragments. Given the high degree of co-capture bias observed with CHi-C, we focused on Promoter-Enhancer and Promoter-CTCF interactions. While both methods enriched for active enhancers, the extent of enrichment was greater in NuTi Capture-C (Supp. Fig. 29b). Therefore, NuTi Capture-C can be applied to produce unprecedented high-resolution 3C interaction maps at genome-wide scale.

**Fig. 6.**
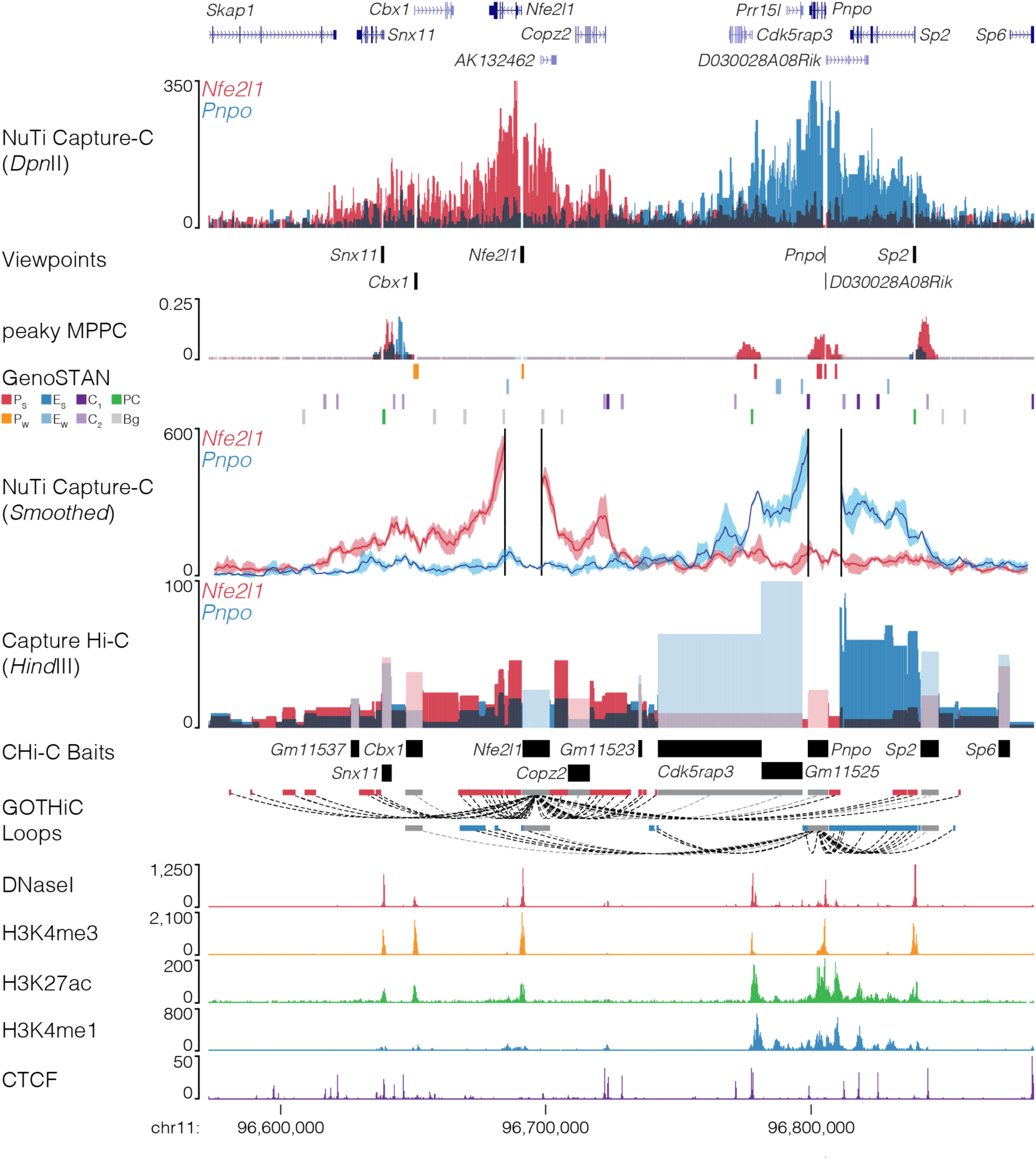
Comparison of capture methods at the *Nfe2l1* and *Pnpo* promoters. Sequence tracks showing the difference between high-resolution 3C (*Dpn*II, NuTi Capture-C) and low-resolution 3C (*Hind*III, Capture Hi-C) from nearby gene promoters (mm9, chr11:96,572,876-96,883,917) in erythroid cells. Tracks in order: UCSC gene annotation, *cis*-normalized mean interactions per *Dpn*II fragment using NuTi Capture-C (n=3), NuTi Capture-C viewpoints, peaky Marginal Posterior Probability of Contact (MPPC) scores with fragments with MPPC ≥0.01 darker, GenoSTAN open chromatin classification, windowed mean interactions using NuTi Capture-C, total supporting reads per *Hind*III fragment with CHi-C (n=2; co-targeted fragments are lighter in colour), CHi-C bait fragments, loops between reported significantly interacting fragments (co-targeting loops are coloured grey), erythroid tracks for open chromatin (DNaseI), promoters (H3K4me3), active transcription (H3K27ac), enhancers (H3K4me1), and boundaries (CTCF). Note overlapping blue and red signals appear darker in colour (NuTi Capture-C, peaky MPPC, CHi-C).

## DISCUSSION

Chromosome conformation capture is a powerful tool for the study of DNA folding within the nucleus. NG Capture-C has been applied to numerous biological questions, including enhancer characterization and super-enhancer dissection^19, 25–28^, understanding the dynamics of Polycomb Bodies^29, 30^ and X-chromosome inactivation^31, 32^, characterizing CTCF boundaries^23, 33^, and mapping the effector genes for polygenic human traits^34, 35^. Despite their widespread applicability, the sequencing needs and cost of high-resolution methods have limited their use in large-scale experiments. To this end we have substantially improved the scale upon which the NG Capture-C method can be employed. Our results show that efficiency gains can be made in both 3C library generation and in targeted enrichment. We have combined these technical improvements into a new method, NuTi Capture-C. Using NuTi Capture-C we generated high-resolution 3C interaction maps for over 8,000 genes in triplicate from erythroid cells. Demonstrating that with thoughtful optimisation of every stage of the process, high-resolution 3C methods can be taken to a genome-wide scale.

In optimising the production of 3C libraries, we found that the soluble and nuclear fractions of *in situ* 3C libraries have vastly different proximity signals and information content. Many statistical methods, including CHiCAGO^36^, peakC^37^, r3C-seq^38^, FourCSeq^39^ and peaky^12^, model this proximity decay curve to identify significant interactions. Our finding that the decay curve can be altered by technical fluctuation will be of particular concern when using these methods, especially when comparing different cell-types, which may respond differently to fixation, lysis, digestion and ligation. Our solution to this was to isolate intact nuclei after ligation. This optimization also reduced the amount of noise from inter-nuclear ligation, the majority of which would be reported as *trans* interactions. This easy to implement protocol adaptation would therefore likely improve any 3C method, leading to more reliable interaction calling, particularly as *trans* gene regulation through interaction has recently emerged as important for control of olfactory receptor genes^40^.

We have also robustly tested the effect of probe length, concentration, and pool composition for 3C enrichment. Shortening the length of probes delivered a predictable yield in higher informative sequencing content and titrating the amount of probe increased the specificity of sequencing. In combination with nuclear isolation these modifications are easily implemented and will lead to immediate benefits, making possible very-large scale 3C capture designs. One consideration when targeting multiple viewpoints is: would the same result be returned by targeting each viewpoint independently, or does co-enrichment skew the underlying interaction frequencies? Through modelling and experimental approaches, we show that co-enrichment in 3C methodologies does introduce bias. Disconcertingly, we find that the bias introduced by co-targeting is affected by both the relative efficiency of viewpoint enrichment and their true interaction frequency. Controlling for this bias is essential to avoid misleading findings, such as the likely overinflated finding of >20 significant promoter-promoter interactions per targeted promoter^11^. For biotinylated oligonucleotide capture, used in Capture-C and Capture Hi-C, specific co-targeting sites are known, therefore bias can be avoided in these methods by simply masking interaction counts between co-targeted fragments. Bias introduced from methods where the target sites are not precisely defined, e.g. immunoprecipitation for ChIA-PET/HiChIP/ChIA-Drop^15–17^ and RNA purification enrichment for HiChIRP^18^, is considerably more complex and at present no such correction for co-enrichment skew is used in these methods. Our findings indicate that to accurately adjust for bias in these methods, researchers must determine the underlying interaction frequency, and the efficiency of targeting at each site. Realistically this could only be done by performing independent 3C (e.g. Hi-C) and enrichment (e.g. ChIP-seq) experiments prior to performing a now moot fusion experiment.

In this paper we have presented NuTi Capture-C, which provides an improved method for targeted high-resolution 3C experiments. The optimizations described here allow NuTi Capture-C to be applied at a genome-wide scale, but could also be implemented to improve the quality and reproducibility of other 3C techniques. Using this method, we expect researchers will be able to provide more reliable insights into biology while studying genome organization throughout growth and development.

## MATERIALS AND METHODS

### Cell culture and fixation

Protocols were approved through the Oxford University Local Ethical Review process. Experimental procedures were performed in accordance with European Union Directive 2010/63/EU and/or the UK Animals (Scientific Procedures) Act, 1986. Murine erythroid cells were obtained from spleens of C57BL/6 or C57BL/6-cross-CBA/J F1 hybrid mice treated with phenylhydrazine (40 mg g^-1^ body weight per dose, with three doses given 12 h apart; mice were killed on day 5). Spleens, consisting of >80 % CD71^+^ ter119^+^ erythroid cells due to hemolytic anemia, were dissociated in Phosphate buffered solution (PBS) and strained through a 30 µM filter (Miltenyi Biotec) to remove clumps. For ter119^+^ selection, 3×10^8^ cells were resuspended in 3 ml of FACS buffer (PBS with 10% FBS) and stained with 0.9 µg anti-ter119-PE (130-102-338; Miltenyi Biotec). Stained cells were conjugated to anti-PE microbeads (130-048-801; Miltenyi Biotec) and passed through 3 LS Columns (Miltenyi Biotec). Mouse embryonic stem cells (ESC) from the feeder free line ES-E14TGA2a.IV (Strain 129/Ola) were grown on 0.1% gelatin (BHK-21 Glasgow Minimal Essential Medium (MEM) [21710025; Invitrogen], 10% Fetal bovine serum (FBS) [10270106; Invitrogen], 2 mM glutamine [25030024; Invitrogen], 100 U ml^-1^ Penicillin-Streptomycin [15140122; Invitrogen], 1 mM sodium pyruvate [11360039; Invitrogen], 1 × MEM non-essential amino acids [11140035; Invitrogen], 0.1 mM 2-mercaptoethanol [31350010; Invitrogen], 1000 U ml^-1^ Leukemia Inhibition Factor) and re-suspended with 0.05% trypsin for 5 minutes 37°C before washing with PBS. Human erythroid cells were generated from CD34+ cells as described^34, 41^ with ethics approval (MREC 03/08/097) and stored according to HTA guidelines (License 12433). Mouse erythroid and ESC were resuspended in RPMI (11875093; Invitrogen) with 15% FBS for fixation. Human erythroid cells were fixed in growth media. For all cell types, cells were resuspended at 1-2×10^6^ cells per ml and fixed at room temperature with 2% v/v formaldehyde for 10 minutes. Fixation was quenched with 120 mM glycine. Cells were washed with ice cold PBS before 3C library preparation.

### 3C library preparation

*In situ* 3C libraries were prepared as previously described^8^. For Nu-3C, cells were lysed on ice in 5 ml lysis buffer (10 mM Tris-HCl, pH 8, 10 mM NaCl, 0.2% Igepal NP-40 (Sigma), 1× cOmplete protease inhibitor (Roche) then pelleted by centrifugation (15 min, 4°C, 500 rcf). Lysis buffer was discarded and nuclei were resuspended in 1 ml PBS before snap freezing and storage at −20°C for up to 12 months. For digestion, up to 5×10^6^ nuclei were defrosted, pelleted (15 min, 4°C, 500 rcf) then resuspended in 215 µl 1× *Dpn*II buffer. Nuclei were then permeabilized with 0.28% SDS in a single reaction (200 µl nuclei, 60 µl 10× *Dpn*II buffer, 434 ml PCR grade water, 10 µl 20% vol/vol SDS) and one undigested control (15 µl nuclei, 28.5 µl 10× *Dpn*II buffer, 227.5 ml PCR grade water, 4 µl 20% vol/vol SDS) for 1 hour at 37°C on a thermomixer (500 rpm). SDS was quenched into micelles for one hour by addition of 20% Triton-X (1.67% final concentration, 66 µl for digest and 25 µl for the undigested control). *Dpn*II was added to digests in three aliquots of 10µl (500 U) spaced several hours apart for a total digest time of 16-24 hours at 37°C. *Dpn*II was neutralized by incubation at 65°C for 15 minutes and then immediate transfer to ice to reduce potential for de-crosslinking. 100 µl was removed from the digestion reaction and combined with 200 µl PCR grade water as an un-ligated control. Controls were de-crosslinked, Proteinase-K treated, RNAse A treated, and phenol chloroform extracted as described below. Crosslinked digested DNA was re-ligated by addition of 240 U T7 ligase (500 ml PCR grade water, 134 ml 10× ligation buffer, 8 µl ligase) and incubated overnight at 16°C on a thermomixer (500 rpm). Following ligation, nuclei were isolated by centrifugation (15 min, 4°C, 500 rcf) and the supernatant, containing both freed DNA and the high levels of DTT from the ligation buffer, discarded. Nuclei were resuspended in 300 µl of TRIS-EDTA and de-crosslinked overnight at 65°C with 5 µl Proteinase-K (3 U). RNA was removed by treatment with 5 µl RNAse A (7.5 mU) for 30 minutes at 37°C. DNA was extracted by addition of 310 µl phenol-chloroform-isoamylalcohol with thorough vortexing before transfer to a phase lock tube and centrifugation (10 min, 12,600 rcf, room temp). The upper layer was transferred to a new tube and DNA precipitated overnight at −20°C (30 µl 3M sodium acetate, 1 µl glycoblue, 900 µl 100% ethanol). DNA was pelleted by centrifugation (30 min, 21,000 rcf, 4°C) and washed twice with 70% ice cold ethanol before resuspension in 150 µl water (30 µl for controls). Samples and controls were quantified using Qubit (Invitrogen), run on a 1% agarose gel and tested by qPCR to determine library quality. Only libraries with a digestion efficiency >70% were used for Capture-C.

### Library indexing

Libraries were either indexed with NEBNext DNA Library Prep Master Mix for Illumina (New England Biolabs) using 6 µg input 3C DNA as previously described^8^ or using NEBNext Ultra II DNA Library Prep Kit for Illumina (New England Biolabs). When using the Ultra II kit 3 µg 3C material was sonicated to 200 bp as previously described^8^, and purified using Ampure XP SPRI beads (Beckman Coulter). DNA was eluted into 53 µl with 1 µl used for D1000 TapeStation analysis (Agilent) and 2 µl used for Qubit quantification (Invitrogen). 50 µl of DNA (≤2 µg) was then indexed with the following modifications; for the End Prep reaction, the 20°C incubation was lengthened to 45 min, 5 µl of NEBNext Adaptor was added and incubated for 30 min at 20°C, the USER Enzyme incubation was extended to 30 min (37°C), and indexing was performed in two reactions with Herculase II Fusion Polymerase (Agilent) using 6 cycles of amplification as previously described^8^.

### Oligonucleotide synthesis and capture

Pools of biotinylated oligonucleotides (Supp. Table 1) were sourced from IDT, Sigma or synthesized in house or had been previously reported^8, 21, 34^. We synthesized biotinylated oligonucleotides on a Combimatrix CustomArray B3 DNA synthesiser (B3Synth_v25.1 software) using CustomArray 12K Blank Slides (CustomArray Inc., PN: 2000100-Oligo pool Application) as described^42^. Oligonucleotide pull down for single and double capture of multiplexed 3C libraries was performed using the Nimblegen SeqCap EZ kit as previously described^8^ with various masses of oligonucleotides with 10 cycles of DNA amplification.

### Sequencing and Data analysis

Fastq reads for small design captures were generated using paired-end sequencing (75/75, and 150/150 cycles) on either a MiSeq or NextSeq Illumina platform. The active gene design was sequenced by Novogene (Hong Kong) using 75/75bp paired-end reads on the Illumina NovaSeq platform to generate at least 10^5^ read pairs per viewpoint for each of the three libraries. Sequenced reads were processed using either CCseqBasic^43^ or a modified script (CCseqBasicM) which improves throughput for thousands of oligonucleotides by parallelising analyses for groups of targets (available on Github: https://github.com/Hughes-Genome-Group/CCseqBasicM). Target enrichment was calculated as the percent of mapped read pairs containing the target fragment divided by the total number of restriction endonuclease fragments in the genome. For sequencing depth analysis, deeply sequenced human data was used (GSE129378). Reporter counts were normalized to reporters per 100,000 *cis* reporter fragments and replicates combined using CaptureCompare^43^. Alignment of *Hbb-b1/2* oligonucleotides to off target peaks was performed with Clustalω in MacVector. Statistical comparisons were carried out using Prism. Genes were characterized as active or inactive using published H3K4me3, H3K27ac, DNaseI-seq and RNA-seq data^19, 23^. Peaky analysis was performed on the average reporter count per fragments as described^34^ with the following modification: to adjust for overcalling in bins with sparse data residuals were normalized to have a mean of 0 and a standard deviation of 1 in each distance bin. We performed chromatin segmentation of ter119^+^ erythroid cells using GenoSTAN^24^. Segmentation used a peak centric approach, rather than signal across the whole genome, with H3K4me1, H3K4me3, H3K27ac, and CTCF (GSE97871, GSE78835)^19, 23^ read coverage calculated (deepTools^44^ v2.4.2) for 1 kb windows over open chromatin peaks (bedtools^45^ merge -d 10) to capture histone modifications. The HMM model was trained using Poisson log-normal distributions with 10 initial states. These were manually curated to eight final states based on similarity of chromatin signature.

## ACKNOWLEDGEMENTS

We thank Gerton Lunter, Ed Sanders and Ed Morrissey for their insights into the skew model. This work was carried out as part of the WIGWAM Consortium (Wellcome Investigation of Genome Wide Association Mechanisms) funded by a Wellcome Trust Strategic Award (106130/Z/14/Z) and Medical Research Council (MRC) Core Funding (MC_UU_00016). S.d.O was supported by an MRC Project Award (MR/N00969X/1) to J.R.H., T.B., and V.J.B. Wellcome Trust Doctoral Programmes supported C.Q.E. (203141/Z/16/Z), R.S. (203728/Z/16/Z), and A.M.O. (105281/Z/14/Z), who was also supported by the Stevenson Junior Research Fellowship (University College, Oxford). J.O.J.D. is funded by an MRC Clinician Scientist Award (MR/R008108) and received Wellcome Trust Support (098931/Z/12/Z).

## AUTHOR CONTRIBUTIONS

D.J.D., J.R.H., A.M.O., J.K., and J.O.J.D. designed experiments. D.J.D., M.E.G., S.J.H., and L.N. performed experiments. D.J.D., M.E.G., J.T., N.R., C.Q.E., and R.S. analysed data. S.D.O, A.S., and A.E-S, generated essential reagents. Funding was acquired by T.B. V.J.B. and J.R.H., who also supervised works carried out. D.J.D wrote the manuscript and made the figures.

## AVAILABILITY OF DATA AND MATERIALS

Sequence reads and processed data for the active gene capture have been archived with GEO (GSEREF). Profiles for interactions of active genes in mouse erythroid cells are available at https://capturesee.molbiol.ox.ac.uk/projects/capture_compare/1086.

## SUPPLEMENTARY MATERIAL

**Supplementary Table 1. Capture oligonucleotide sequence and co-ordinates (mm9)**

**Supplementary Note. Mathematical modelling of co-capture bias**

**Supp. Fig. 1.**
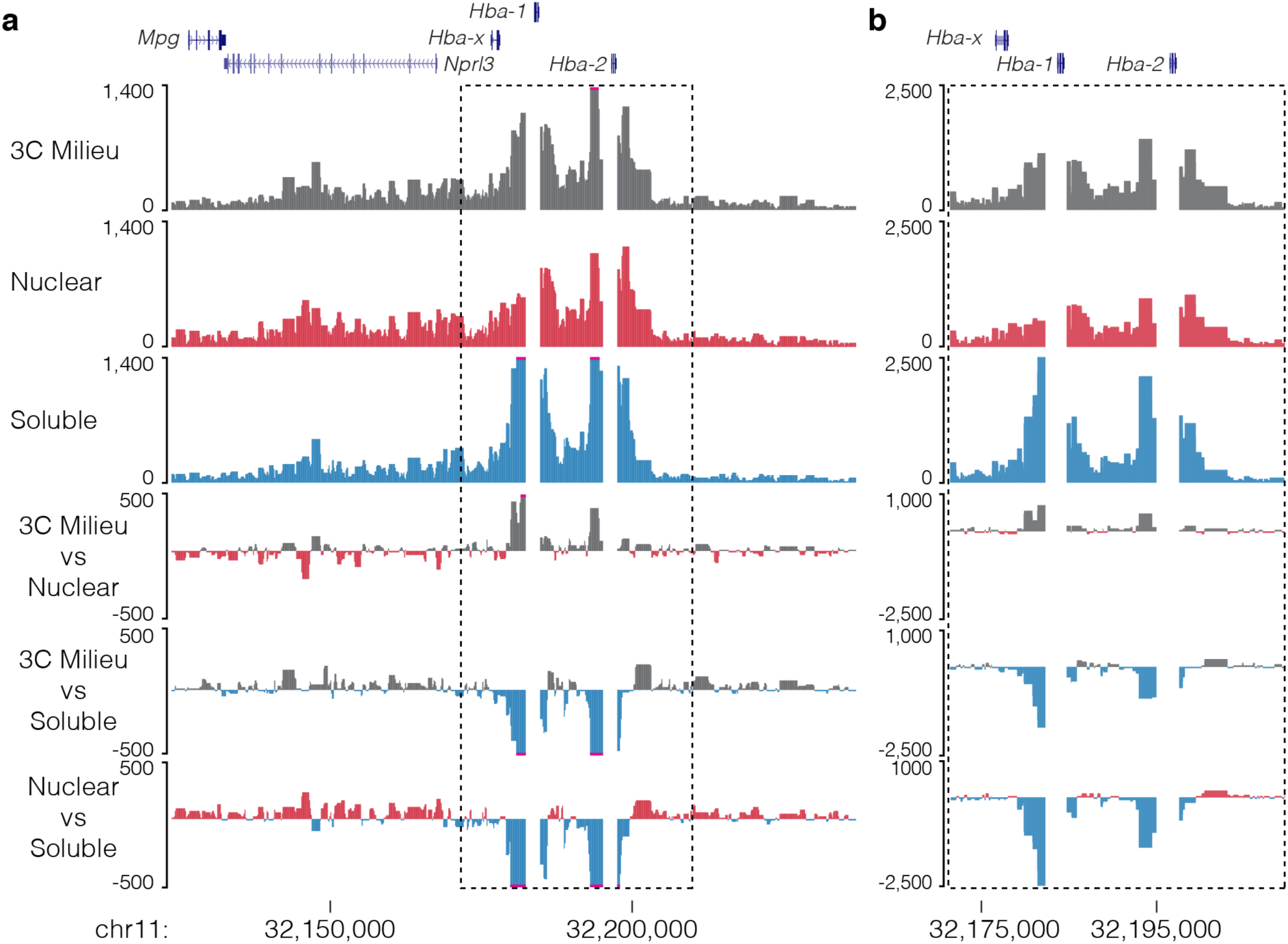
Soluble 3C material has a higher proximity signal. Capture profiles and comparison tracks for *Hba-1/2* capture in mouse erythroid cells **(a)** from total 3C library Milieu or its fractionated nuclear and soluble fractions shows soluble material has a higher proximity signal **(b)**, likely from small diffusing chunks of digested crosslinked chromatin.

**Supp. Fig. 2.**
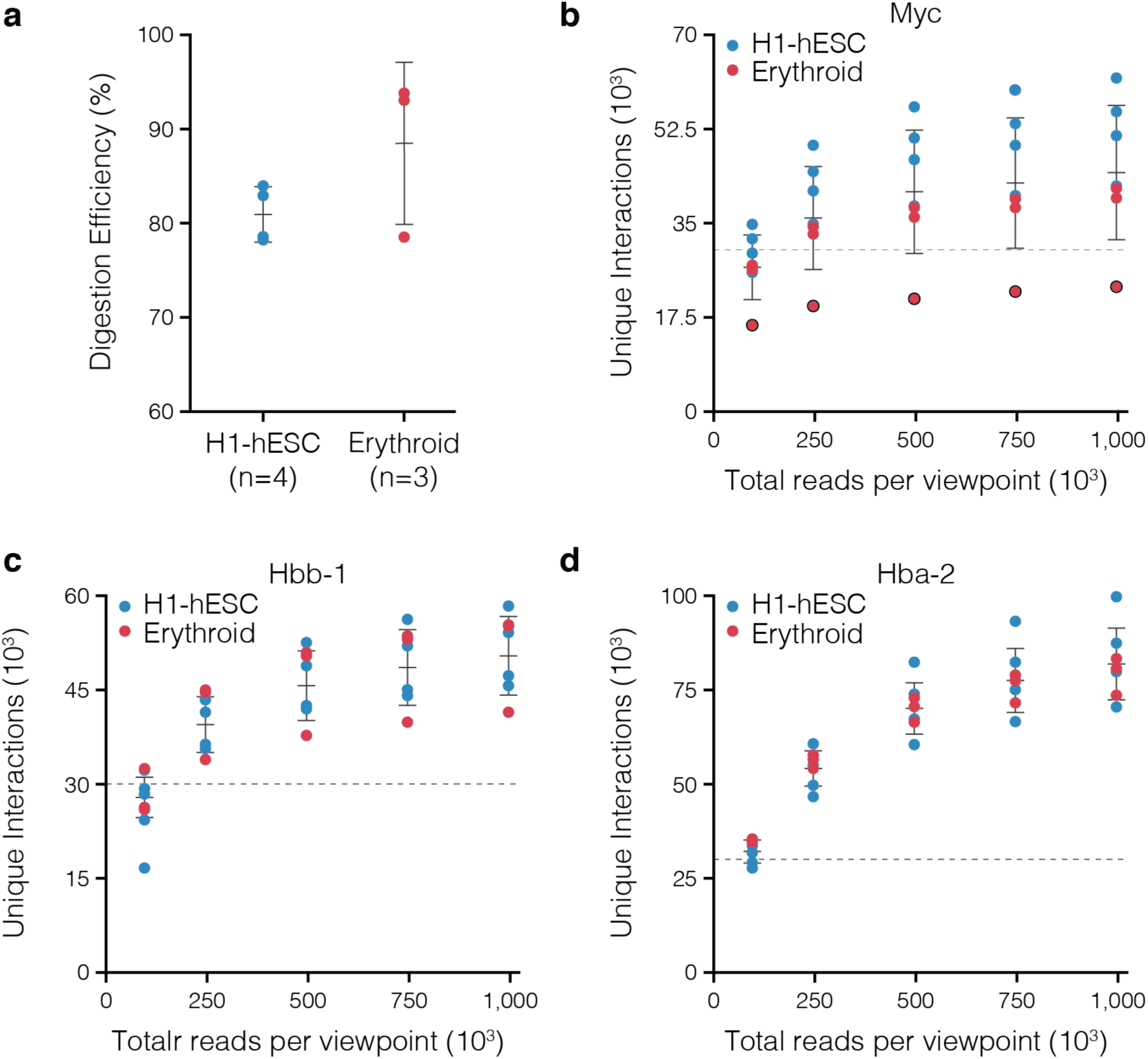
Reporter sensitivity through sequencing depth. **a,** Digestion efficiency for Nu-3C libraries from human embryonic stem cells (H1-hESC) and erythroid cells. NuTi Capture-C was performed for the seven multiplexed libraries targeting *Myc* **(b)**, *Hbb-b1/2* **(c)**, and *Hba-1/2* **(d)** and sequenced to over 10^6^ reads per viewpoint per library. Sequence files were subsampled and analyzed to determine number of unique reporters. Dashed lines represent 30,000 unique reporters, or extremely high-sensitivity capture. For *Myc*, one donor has a polymorphism that removes one of the two *Dpn*II sites on the targeted fragment (black outline) – illustrating the effect of using a single probe. Bars show mean and one standard deviation.

**Supp. Fig. 3.**
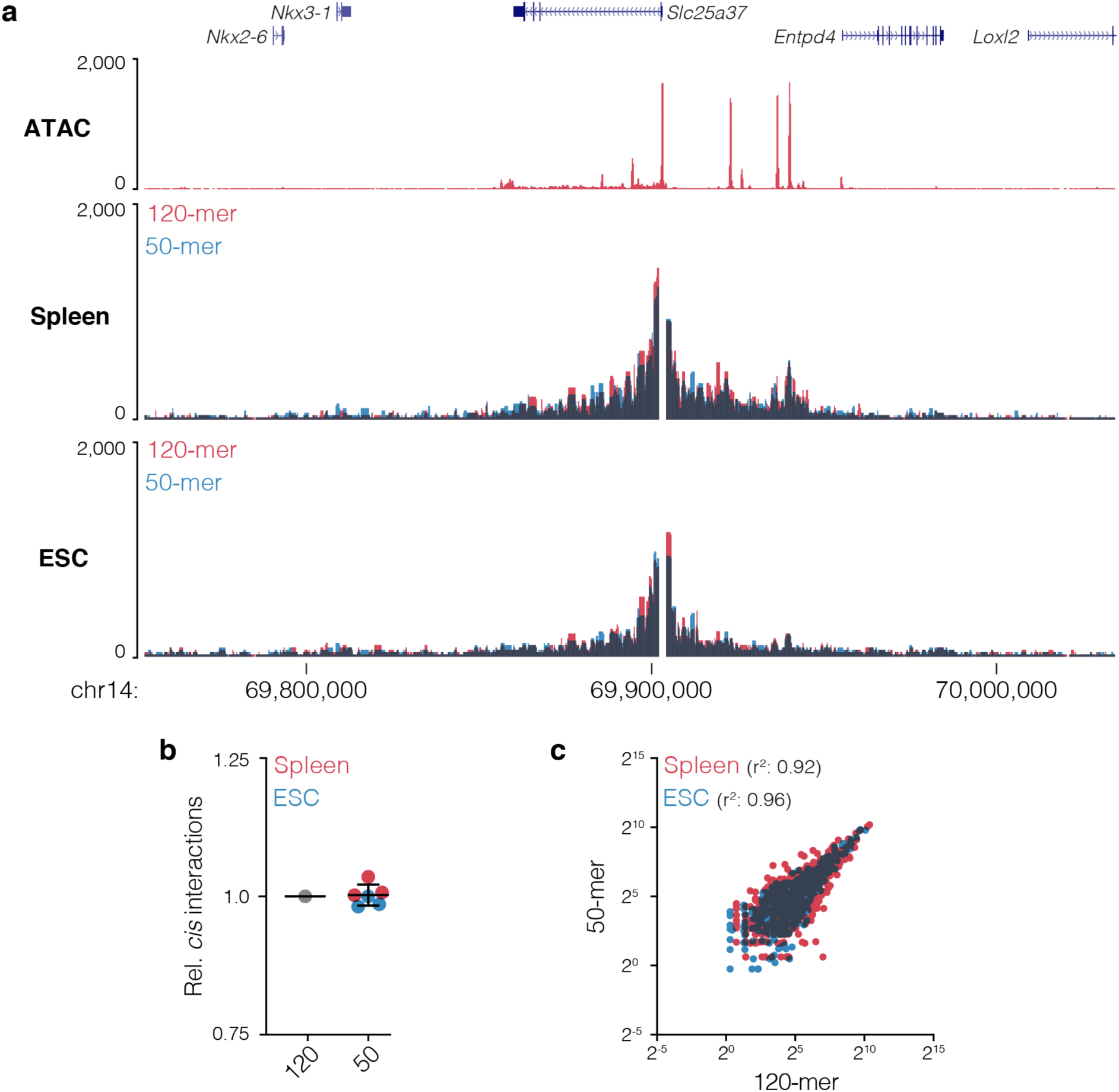
Capture of *Slc25a37* with short probes. **a,** Overlaid 3C interaction profile for *Slc25a37*, which encodes mitoferrin, from mouse erythroid (n=3) and embryonic stem cells (ESC, n=3) captured with either 120-mer or 50-mer probes. Darkened areas show overlapping signals. **b**, Number of cis reporters relative to 120-mer capture. **c**, Comparison of interactions counts from using long or short probes for fragments displayed in panel **a** with Pearson’s correlation.

**Supp. Fig. 4.**
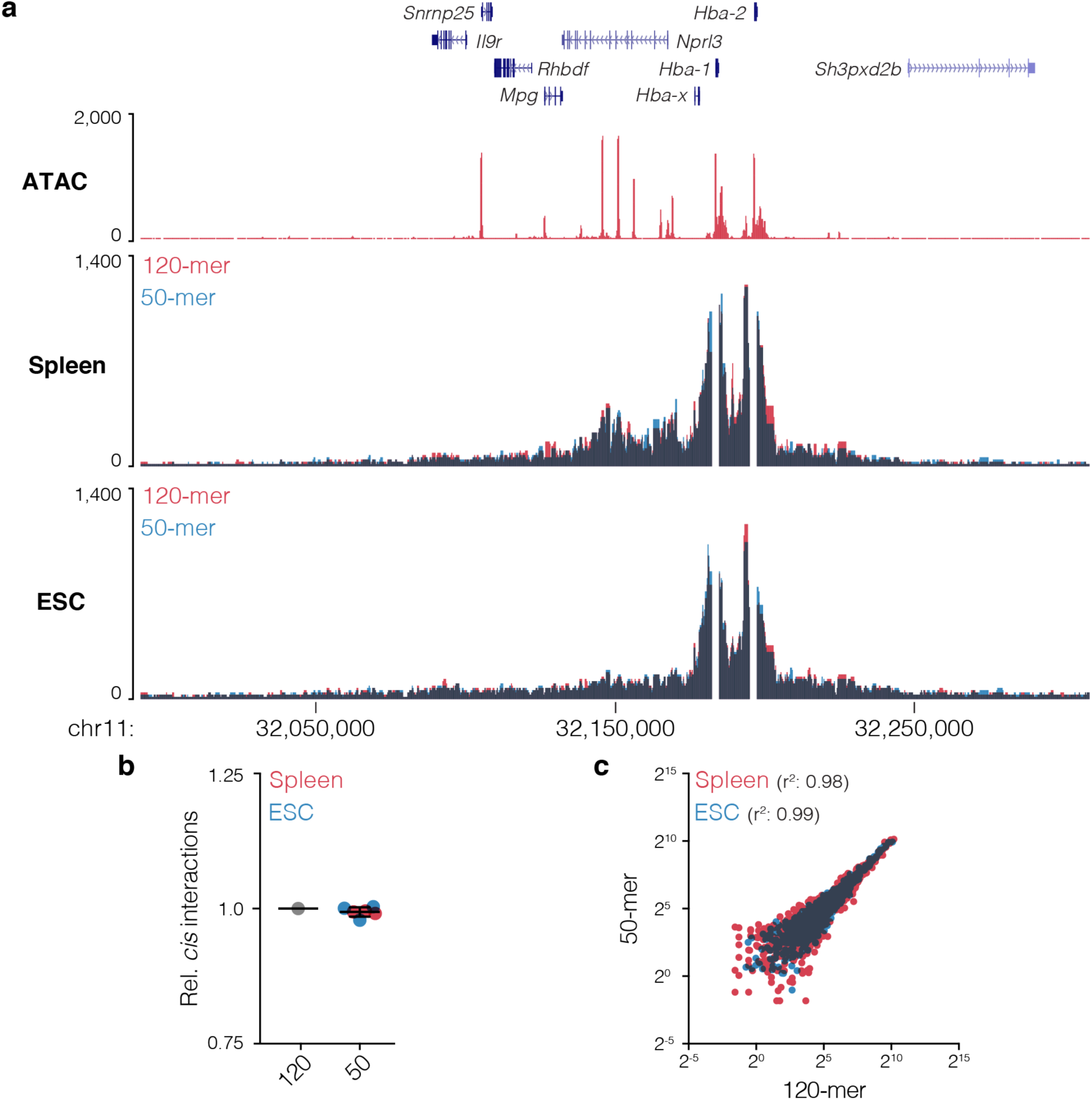
Capture of *α*-globin locus with short probes. **a,** Overlaid 3C interaction profile for *Hba-1 and Hba-2*, which encode *α*-globin, from mouse erythroid (n=3) and embryonic stem cells (ESC, n=3) captured with either 120-mer or 50-mer probes. Darkened areas show overlapping signals. **b**, Number of cis reporters relative to 120-mer capture. **c**, Comparison of interactions counts from using long or short probes for fragments displayed in panel **a** with Pearson’s correlation.

**Supp. Fig. 5.**
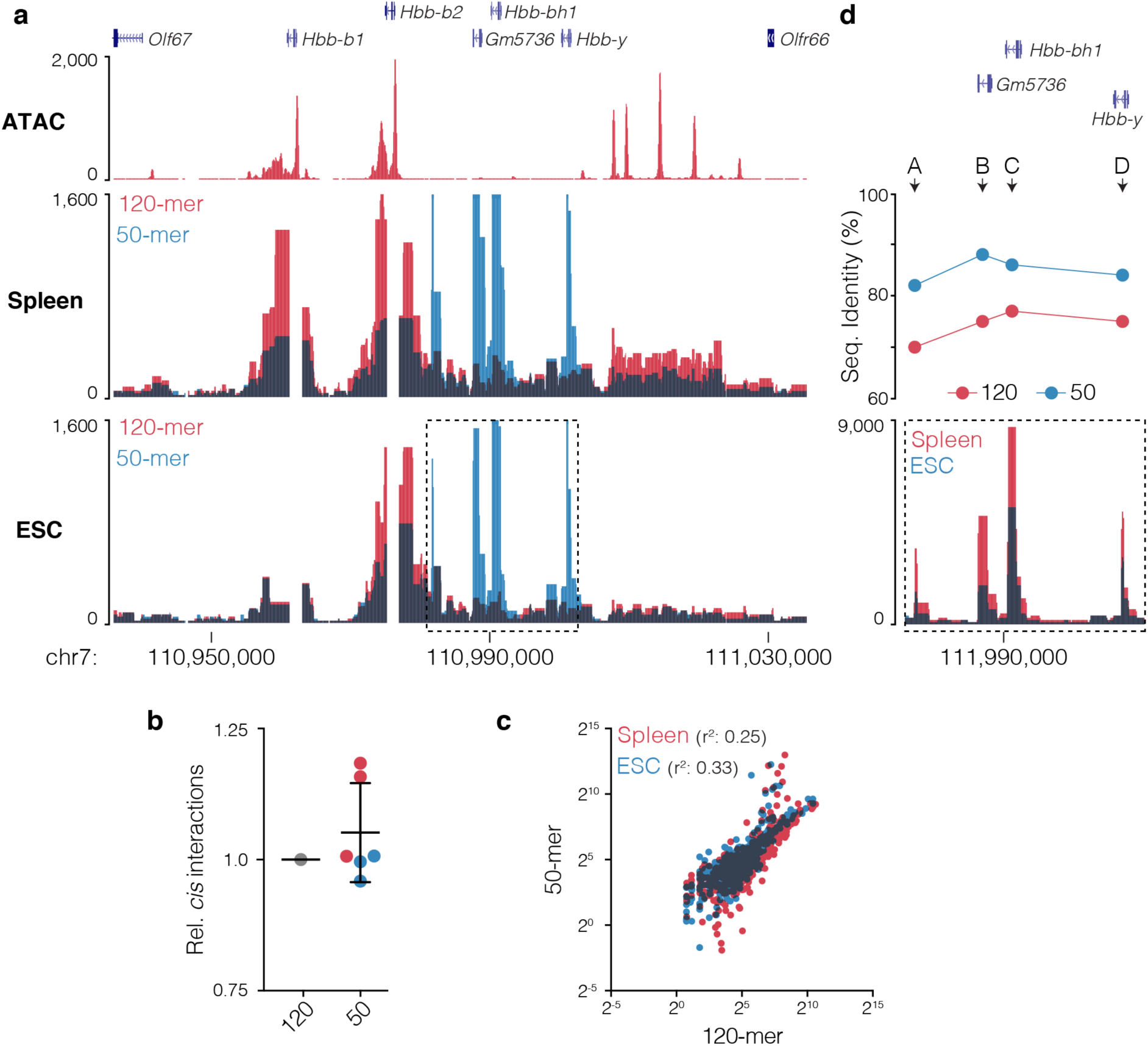
Capture of *β*-globin locus with short probes. **a,** Overlaid *cis*-normalized 3C interaction profile for *Hbb-b1 and Hbb-b2*, which encode *β*-globin, from mouse erythroid (n=3) and embryonic stem cells (ESC, n=3) captured with either 120-mer or 50-mer probes. Darkened areas show overlapping signals. **b**, Number of cis reporters relative to 120-mer capture. **c**, Comparison of interactions counts from using long or short probes for fragments displayed in panel **a** with Pearson’s correlation. **d,** Clustalω determined sequence identity for 120-mer and 50-mer probes with the four novel peaks of interactions seen with the shorter probes.

**Supp. Fig. 6.**
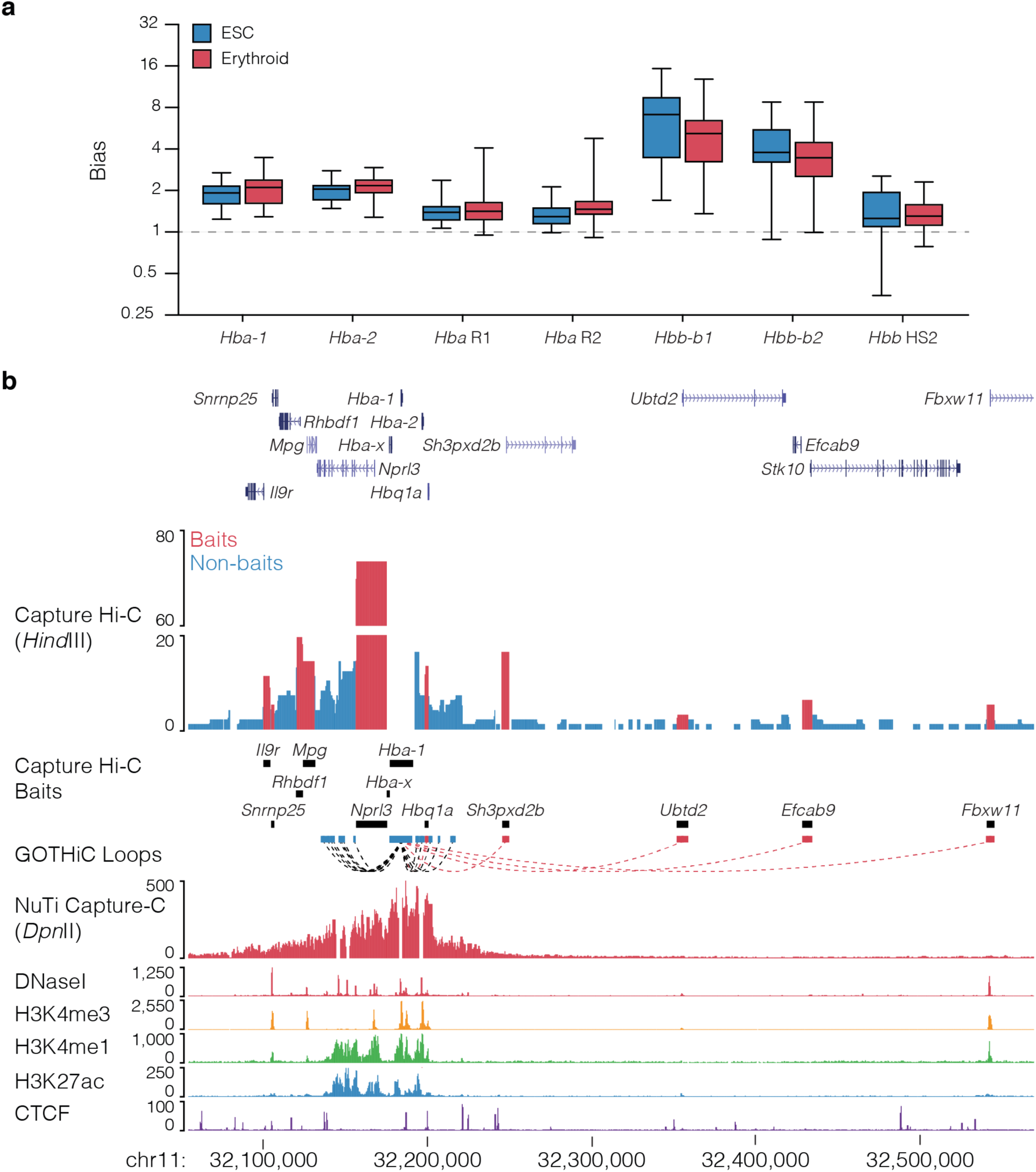
Co-targeting bias observed in Capture-C and Capture Hi-C. **a,** Per viewpoint levels of bias at co-targeted viewpoints around the *α*-globin locus (*Hba-1* and *Hba-2* promoters, R1 and R2 enhancers) and the *β*-globin locus (*Hbb-b1* and *Hbb-b2* promoters, HS2 enhancer). Levels of bias varies across viewpoints and co-targeted fragments but not between erythroid and embryonic stem cells (ESC) indicating bias is primarily caused through the identity of the targeted fragment rather than by cell type signal. **b,** Comparison of the 3C interaction profiles for *Hba-1* generated with Capture Hi-C (targeting all promoters) and NuTi Capture-C (targeting specifically *Hba-1/2* and their two main enhancers – excluded from analysis and seen as gaps in the signal). Total interaction counts for CHi-C in erythroid cells are shown (n=2), fragments and reported significant interactions involving co-targeting coloured red. Note that the peaks over reported long-range significant interactions are not present in NuTi Capture-C and occur specifically at co-targeted fragments (and not adjacent fragments). Erythroid tracks show open chromatin (DNaseI), promoters (H3K4me3), active transcription (H3K27ac), enhancers (H3K4me1), and boundaries (CTCF).

**Supp. Fig. 7.**
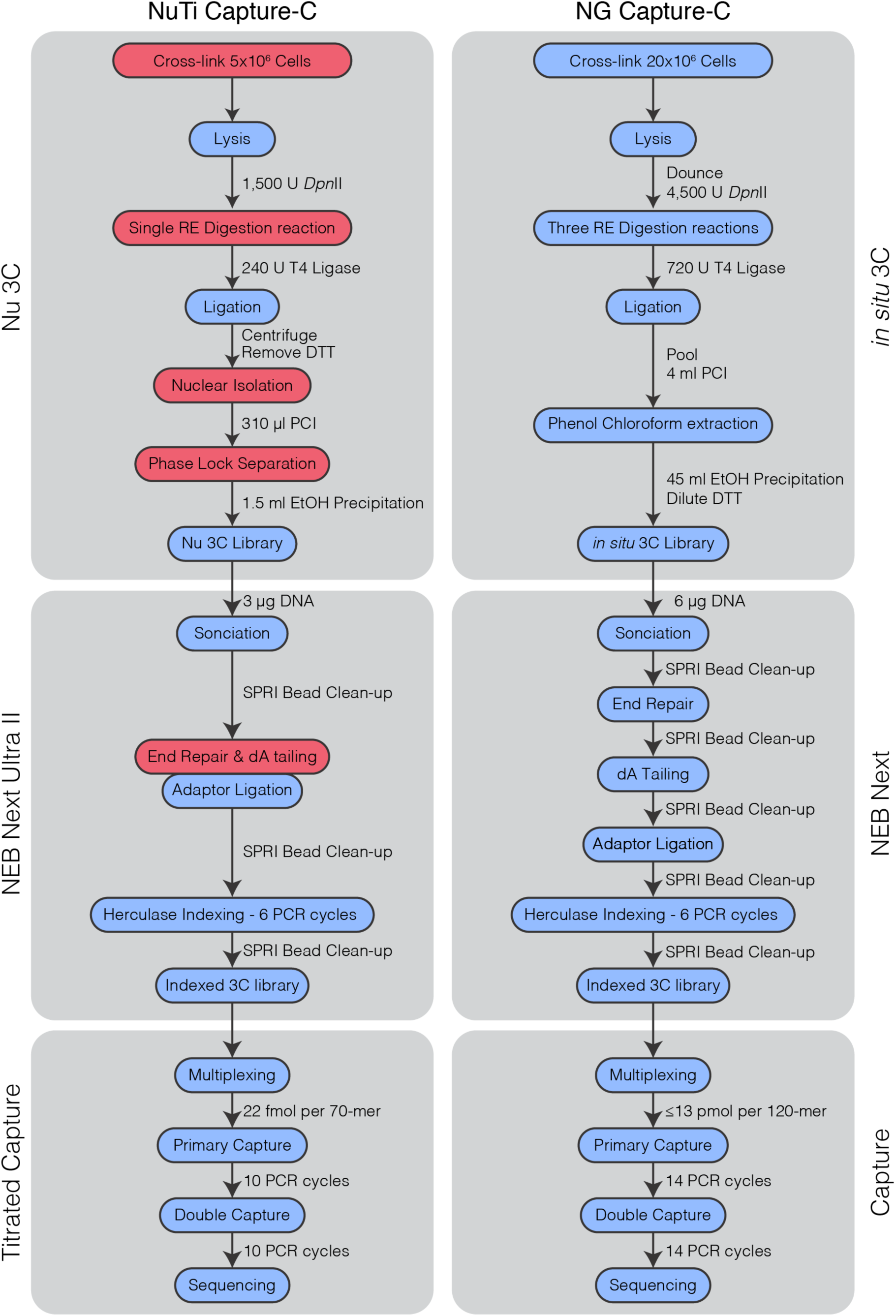
Capture-C workflows. Comparison of experimental workflows for Nuclear-Titratred (NuTi) and Next Generation (NG) Capture-C. Main steps are in blue bubbles, with key innovations for NuTi Capture-C highlighted by red bubbles. Differences in reagents and PCR cycles are shown at individual steps. DTT: Dithiothreitol, PCI: Phenol-Chloroform Isoamyl-alcohol, EtOH: ethanol, SPRI: solid phase reversible immobilisation.

**Supp. Fig. 8.**
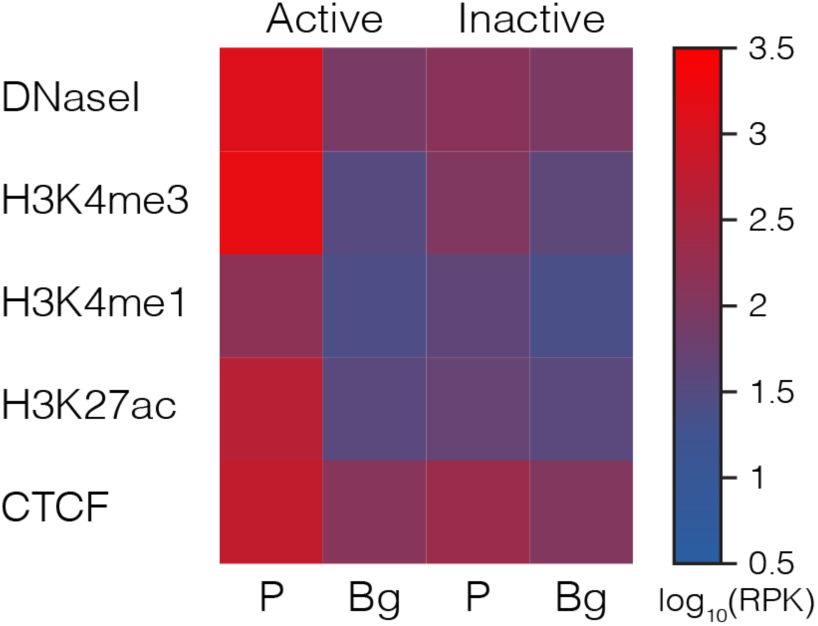
Chromatin signature of captured promoters. Average sequence coverage signature of promoter (P) containing fragments (±1kb) classified as active (n=7,014) or inactive (n=181). Chromatin marks from mouse erythroid cells show open chromatin (DNaseI), promoters (H3K4me3), active transcription (H3K27ac), enhancers (H3K4me1), and boundaries (CTCF). Background (Bg) signal was calculated by generating random peaks of the same number and size using BEDtools shuffle. RPK: reads per kilobase.

**Supp. Fig. 9.**
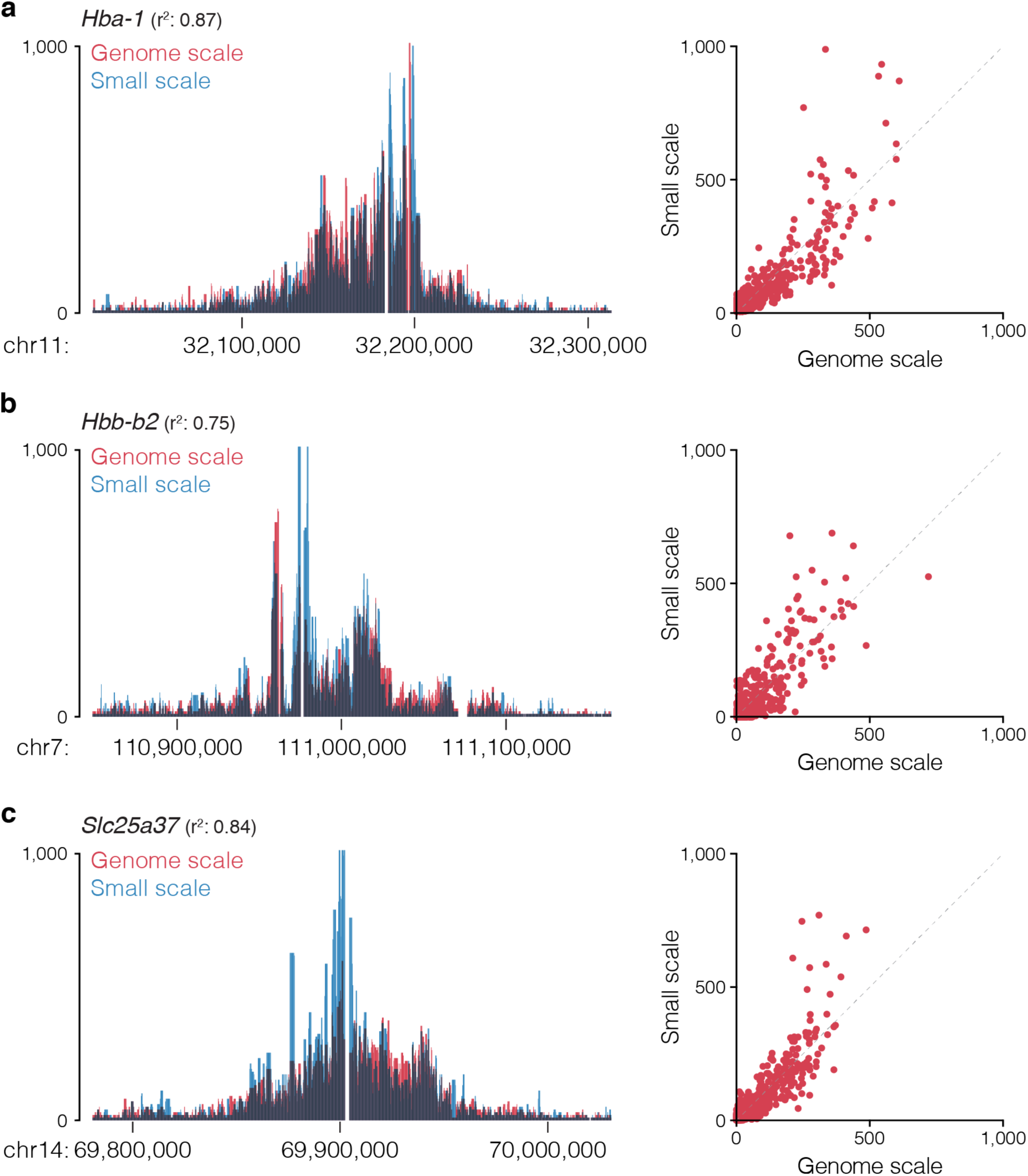
Genome scale capture closely matches designs with fewer probes. Overlaid 3C profiles, Pearson correlation values, and per fragment count correlation plots for the *Hba-1* (**a**), *Hbb-b2* (**b**) and *Slc25a37* (**c**) promoters in mouse erythroid cells when targeting <10 (small scale or >7000 (genome scale) viewpoints with NuTi Capture-C. Note overlaid track go dark where signals overlap, seven values >1,000 not shown.

**Supp. Fig. 10.**
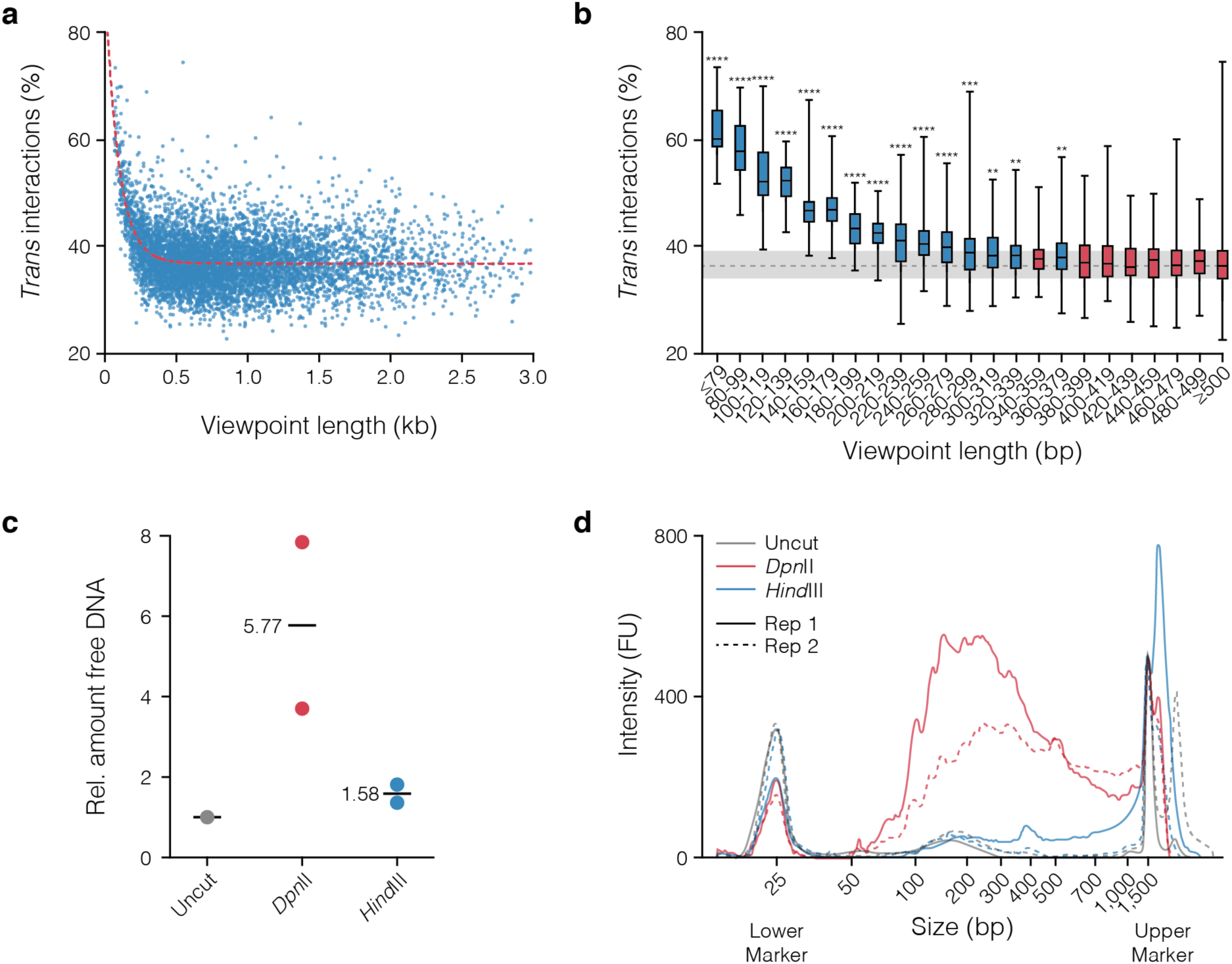
Short fragments have higher levels of *trans* interactions. **a,** Plot of mean percent of *trans* interactions (n=3) for all viewpoints shorter than 3 kb (n=6,659). Red line shows a non-linear fit to the data (r^2^=0.2150, d.f. 19,972). **b,** Box and whisker plot of viewpoints shorter than 500 bp in 20bp bins (n≥12). A one-way ANOVA was carried out with multiple comparisons for each bin against all viewpoints over 500 bp (n=5,017). Significantly different bins identified by a Dunnett’s multiple comparisons test are coloured blue (**p<0.005, ***p<0.0005, ****p<0.0001). Relative amount **(c)** and D1000 tapestation profile **(d)** of DNA recovered from the soluble (non-nuclear) fractions of two 3C samples divided across three tubes each and digested overnight with no enzyme (Uncut), a 4-bp cutter (*Dpn*II), and a 6-bp cutter (*Hind*III). FU: Fluorescent units.

**Supp. Fig. 11.**
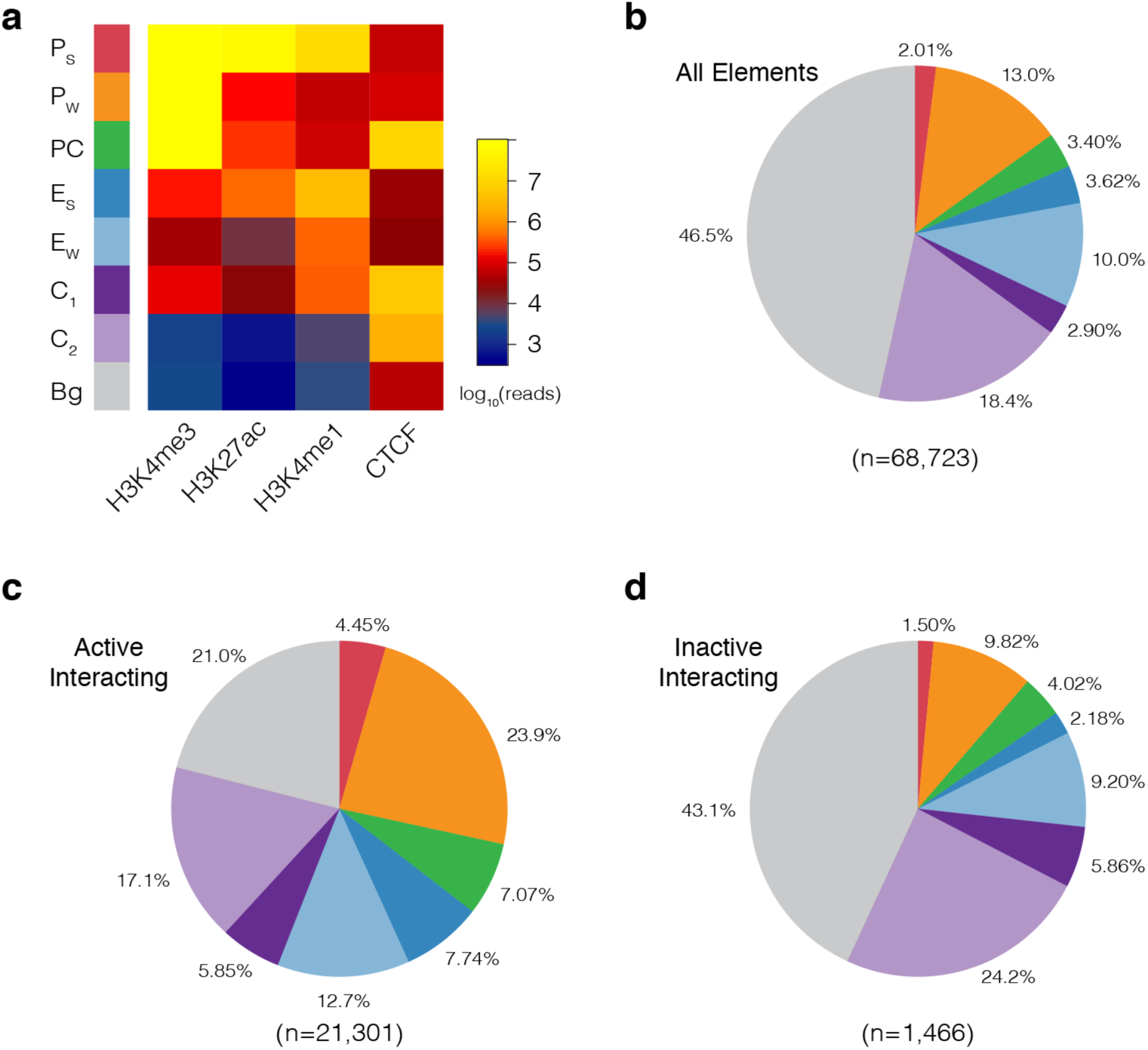
GenoSTAN annotation of the mouse genome in erythroid cells. **a,** Following curation for similar signal profiles, the GenoSTAN Hidden Markov Model identified eight states for 1 kb erythroid open chromatin regions using ChIP-seq for marks associated with promoters (H3K4me3), active transcription (H3K27ac), enhancers (H3K4me1), and boundaries (CTCF). Identified states were named based on average chromatin profile (shown) as: P_S_: Promoter (Strong H3K27ac), P_W_: Promoter (Weak H3K27ac), PC: Promoter/CTCF, E_S_: Enhancer (Strong H3K27ac), E_W_: Enhancer (Weak H3K27ac), C_1_: CTCF near promoter/enhancer, C_2_: CTCF, Bg: Background. Pie charts showing the proportion of unique annotations for all open chromatin regions **(b)**, fragments significantly interacting with active promoters **(c)**, and fragments significantly interacting with inactive promoters **(d)**. Pie chart colours and order match the key in panel a.

**Supp. Fig. 12.**
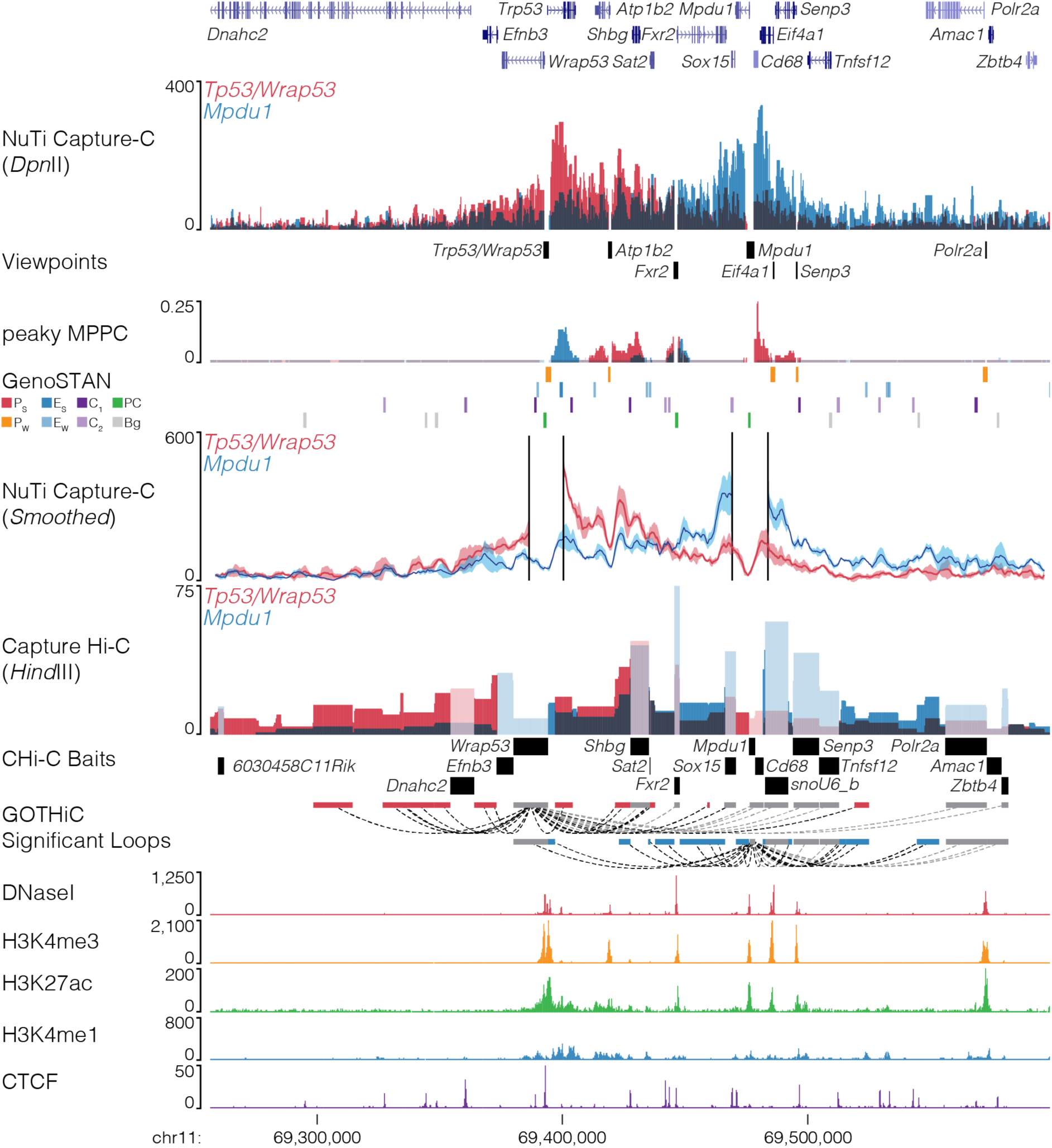
NuTi Capture-C from the *Tp53, Wrap53* and *Mpdu1* promoters. Sequence tracks showing the difference between high-resolution 3C (*Dpn*II, NuTi Capture-C) and low-resolution 3C (*Hind*III, Capture Hi-C) at gene promoters in the same regulatory domain in erythroid cells (mm9, chr11:69,256,536-69,598,480). Tracks in order: UCSC gene annotation, *cis*-normalized mean interactions per *Dpn*II fragment using NuTi Capture-C (n=3), NuTi Capture-C viewpoints, peaky Marginal Posterior Probability of Contact (MPPC) scores with fragments with MPPC ≥0.01 darker, GenoSTAN open chromatin classification, windowed mean interactions using NuTi Capture-C, total supporting reads per *Hind*III fragment with CHi-C (n=2; co-targeted fragments are lighter in colour), CHi-C bait fragments, loops between reported significantly interacting fragments (co-targeting loops are coloured grey), erythroid tracks for open chromatin (DNaseI), promoters (H3K4me3), active transcription (H3K27ac), enhancers (H3K4me1), and boundaries (CTCF). Note overlapping blue and red signals appear darker in colour (NuTi Capture-C, peaky MPPC, CHi-C).

**Fig. 13.**
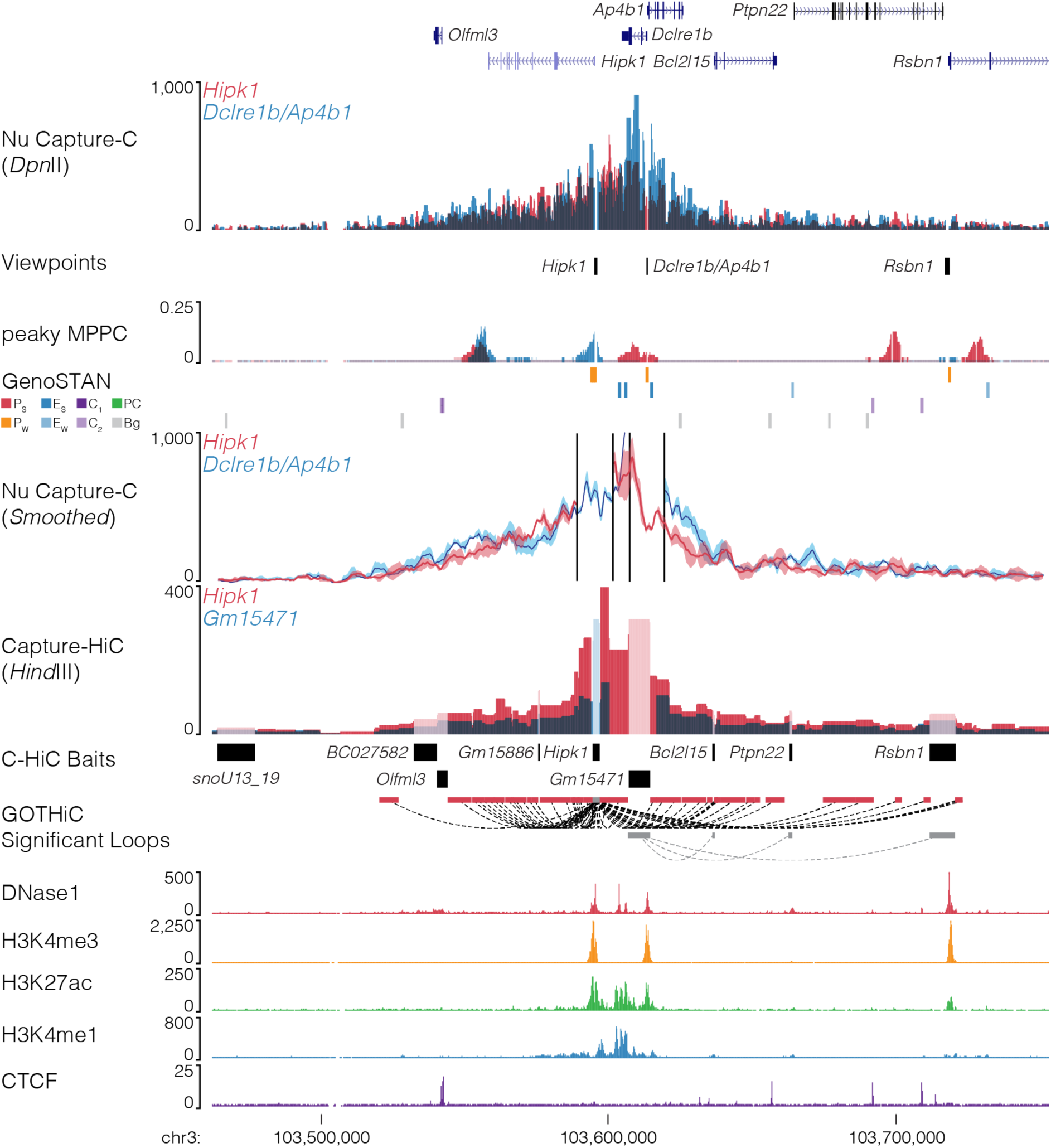
NuTi Capture-C from the *Hipk1, Dclre1b* and *Ap4b1* promoters. Sequence tracks showing the difference between high-resolution 3C (*Dpn*II, NuTi Capture-C) and low-resolution 3C (*Hind*III, Capture Hi-C) at gene promoters in the same regulatory domain in erythroid cells (mm9, chr3:103,462,115-103,753,122). Tracks in order: UCSC gene annotation, *cis*-normalized mean interactions per *Dpn*II fragment using NuTi Capture-C (n=3), NuTi Capture-C viewpoints, peaky Marginal Posterior Probability of Contact (MPPC) scores with fragments with MPPC ≥0.01 darker, GenoSTAN open chromatin classification, windowed mean interactions using NuTi Capture-C, total supporting reads per *Hind*III fragment with CHi-C (n=2; co-targeted fragments are lighter in colour), CHi-C bait fragments, loops between reported significantly interacting fragments (co-targeting loops are coloured grey), erythroid tracks for open chromatin (DNaseI), promoters (H3K4me3), active transcription (H3K27ac), enhancers (H3K4me1), and boundaries (CTCF). Note overlapping blue and red signals appear darker in colour (NuTi Capture-C, peaky MPPC, CHi-C).

**Fig. 14.**
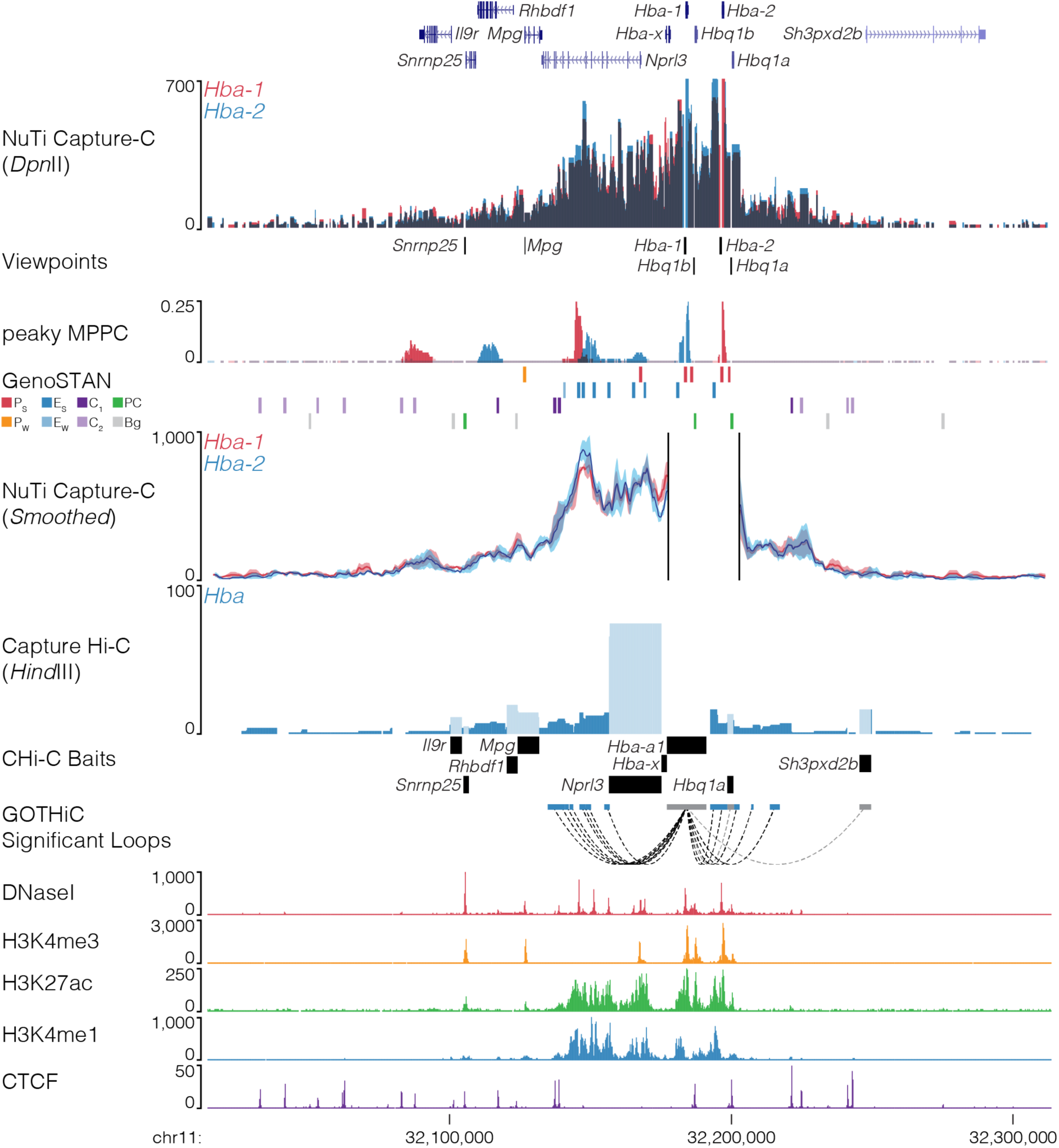
NuTi Capture-C from the *Hba-1* and *Hba-2* promoters. Sequence tracks showing the difference between high-resolution 3C (*Dpn*II, NuTi Capture-C) and low-resolution 3C (*Hind*III, Capture Hi-C) at gene promoters in the same regulatory domain in erythroid cells (mm9, chr3:103,462,115-103,753,122). Tracks in order: UCSC gene annotation, *cis*-normalized mean interactions per *Dpn*II fragment using NuTi Capture-C (n=3), NuTi Capture-C viewpoints, peaky Marginal Posterior Probability of Contact (MPPC) scores with fragments with MPPC ≥0.01 darker, GenoSTAN open chromatin classification, windowed mean interactions using NuTi Capture-C, total supporting reads per *Hind*III fragment with CHi-C (n=2; co-targeted fragments are lighter in colour), CHi-C bait fragments, loops between reported significantly interacting fragments (co-targeting loops are coloured grey), erythroid tracks for open chromatin (DNaseI), promoters (H3K4me3), active transcription (H3K27ac), enhancers (H3K4me1), and boundaries (CTCF). Note overlapping blue and red signals appear darker in colour (NuTi Capture-C, peaky MPPC, CHi-C).

**Fig. 15.**
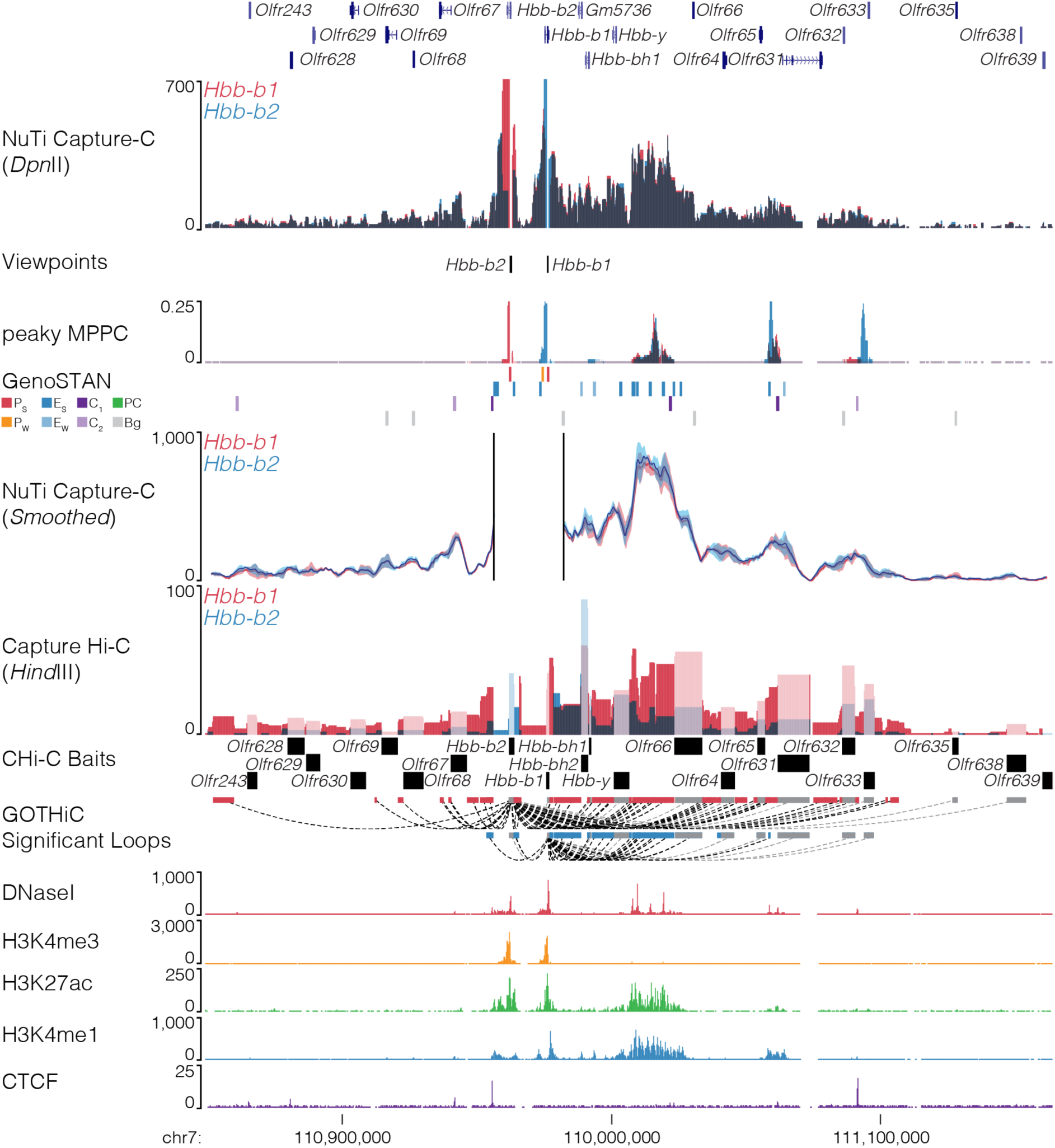
NuTi Capture-C from the *Hbb-b1* and *Hbb-b2* promoters. Sequence tracks showing the difference between high-resolution 3C (*Dpn*II, NuTi Capture-C) and low-resolution 3C (*Hind*III, Capture Hi-C) at gene promoters in the same regulatory domain in erythroid cells (mm9, chr7:110,848,909-111,163,908). Tracks in order: UCSC gene annotation, *cis*-normalized mean interactions per *Dpn*II fragment using NuTi Capture-C (n=3), NuTi Capture-C viewpoints, peaky Marginal Posterior Probability of Contact (MPPC) scores with fragments with MPPC ≥0.01 darker, GenoSTAN open chromatin classification, windowed mean interactions using NuTi Capture-C, total supporting reads per *Hind*III fragment with CHi-C (n=2; co-targeted fragments are lighter in colour), CHi-C bait fragments, loops between reported significantly interacting fragments (co-targeting loops are coloured grey), erythroid tracks for open chromatin (DNaseI), promoters (H3K4me3), active transcription (H3K27ac), enhancers (H3K4me1), and boundaries (CTCF). Note overlapping blue and red signals appear darker in colour (NuTi Capture-C, peaky MPPC, CHi-C).

**Fig. 16.**
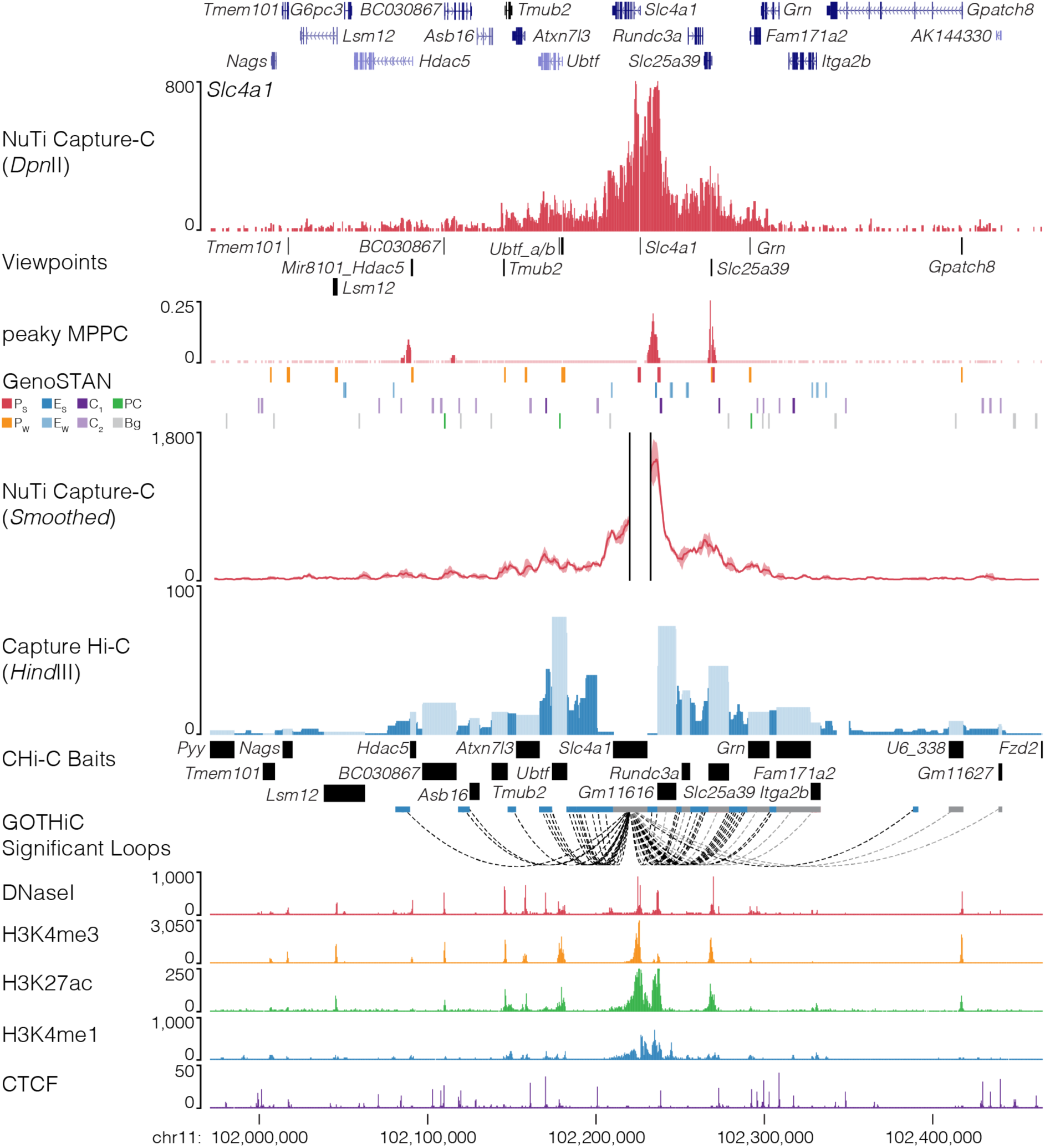
NuTi Capture-C from the *Slc4a1* promoter. Sequence tracks showing the difference between high-resolution 3C (*Dpn*II, NuTi Capture-C) and low-resolution 3C (*Hind*III, Capture Hi-C) at calling interacting fragments (mm9, chr11:101,971,435-102,465,294) in erythroid cells. Tracks in order: UCSC gene annotation, *cis*-normalized mean interactions per *Dpn*II fragment using NuTi Capture-C (n=3), NuTi Capture-C viewpoints, peaky Marginal Posterior Probability of Contact (MPPC) scores with fragments with MPPC ≥0.01 darker, GenoSTAN open chromatin classification, windowed mean interactions using NuTi Capture-C, total supporting reads per *Hind*III fragment with CHi-C (n=2; co-targeted fragments are lighter in colour), CHi-C bait fragments, loops between reported significantly interacting fragments (co-targeting loops are coloured grey), erythroid tracks for open chromatin (DNaseI), promoters (H3K4me3), active transcription (H3K27ac), enhancers (H3K4me1), and boundaries (CTCF). Note overlapping MPPC signals appear darker in colour.

**Fig. 17.**
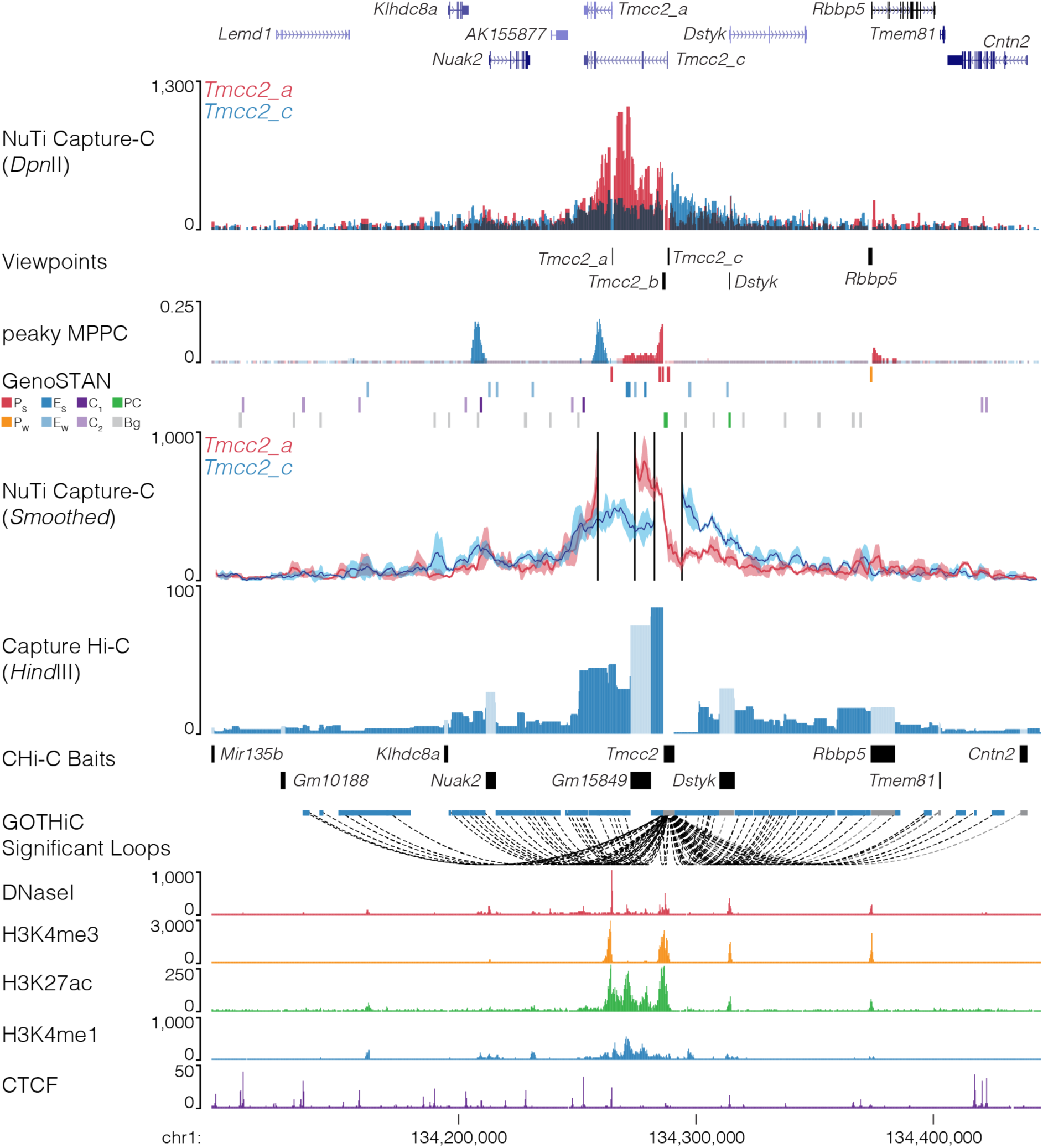
NuTi Capture-C from alternative *Tmcc2* promoters. Sequence tracks showing the difference between high-resolution 3C (*Dpn*II, NuTi Capture-C) and low-resolution 3C (*Hind*III, Capture Hi-C) from alternative *Tmmc2* promoters (mm9, chr1:134,095,540-134,445,539) in erythroid cells. Tracks in order: UCSC gene annotation, *cis*-normalized mean interactions per *Dpn*II fragment using NuTi Capture-C (n=3), NuTi Capture-C viewpoints, peaky Marginal Posterior Probability of Contact (MPPC) scores with fragments with MPPC ≥0.01 darker, GenoSTAN open chromatin classification, windowed mean interactions using NuTi Capture-C, total supporting reads per *Hind*III fragment with CHi-C (n=2; co-targeted fragments are lighter in colour), CHi-C bait fragments, loops between reported significantly interacting fragments (co-targeting loops are coloured grey), erythroid tracks for open chromatin (DNaseI), promoters (H3K4me3), active transcription (H3K27ac), enhancers (H3K4me1), and boundaries (CTCF). Note overlapping blue and red signals appear darker in colour (NuTi Capture-C, peaky MPPC, CHi-C).

**Fig. 18.**
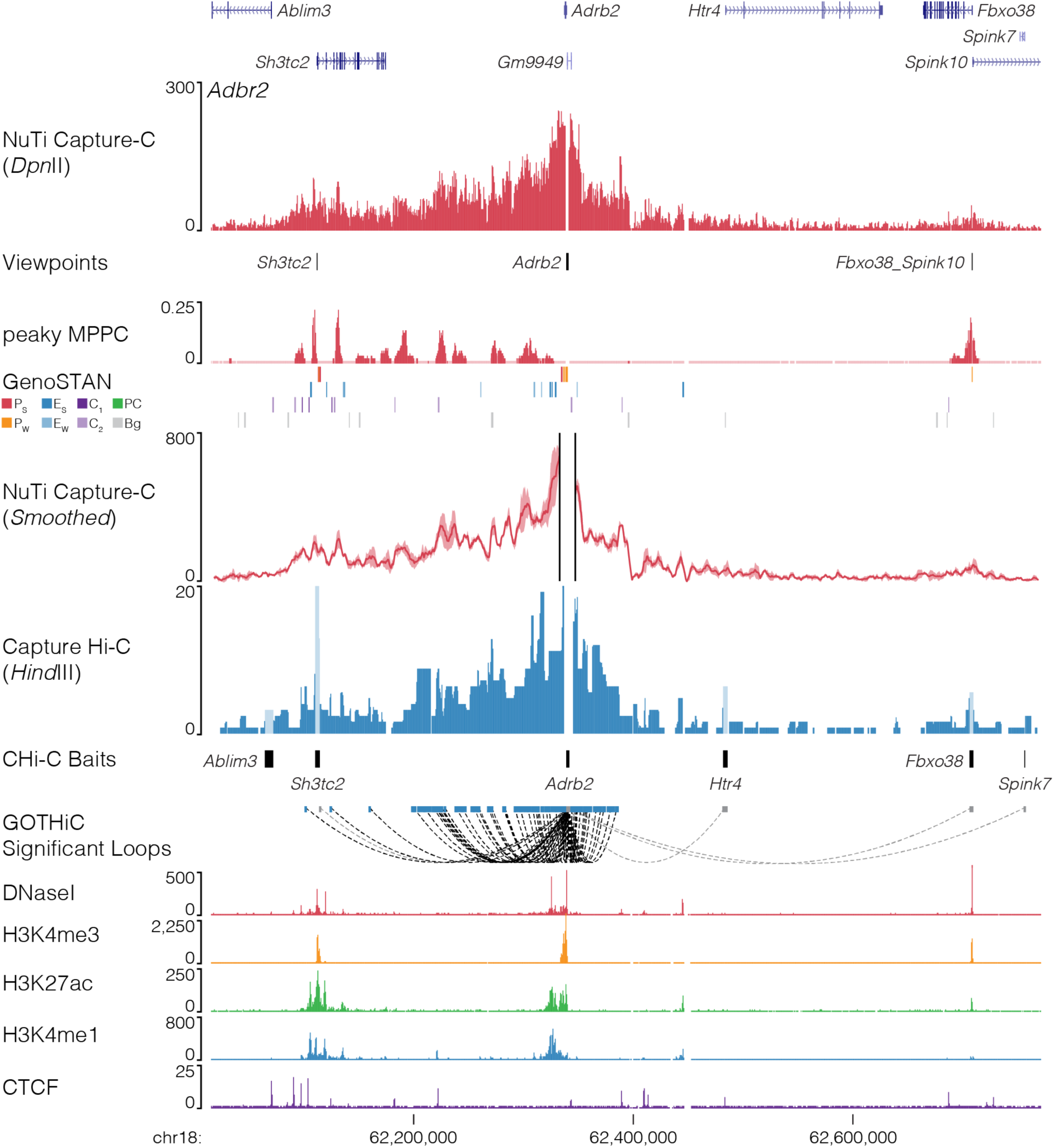
NuTi Capture-C from the *Adrb2* promoter. Sequence tracks showing the difference between high-resolution 3C (*Dpn*II, NuTi Capture-C) and low-resolution 3C (*Hind*III, Capture Hi-C) at calling interacting fragments (mm9, chr18:62,016,212-62,771,180) in erythroid cells. Tracks in order: UCSC gene annotation, *cis*-normalized mean interactions per *Dpn*II fragment using NuTi Capture-C (n=3), NuTi Capture-C viewpoints, peaky Marginal Posterior Probability of Contact (MPPC) scores with fragments with MPPC ≥0.01 darker, GenoSTAN open chromatin classification, windowed mean interactions using NuTi Capture-C, total supporting reads per *Hind*III fragment with CHi-C (n=2; co-targeted fragments are lighter in colour), CHi-C bait fragments, loops between reported significantly interacting fragments (co-targeting loops are coloured grey), erythroid tracks for open chromatin (DNaseI), promoters (H3K4me3), active transcription (H3K27ac), enhancers (H3K4me1), and boundaries (CTCF). Note overlapping MPPC signals appear darker in colour.

**Fig. 19.**
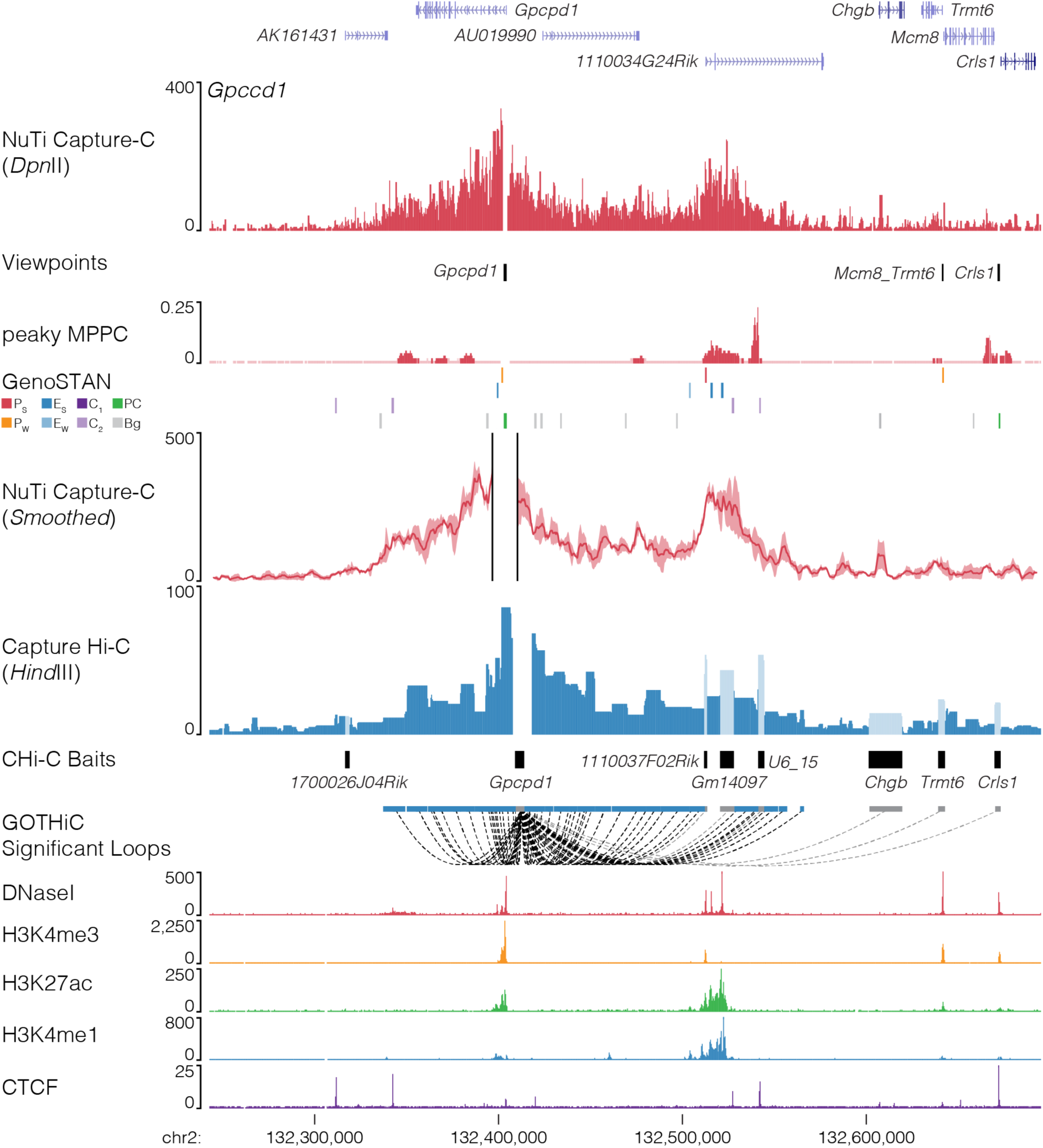
NuTi Capture-C from the *Gpcpd1* promoters. Sequence tracks showing the difference between high-resolution 3C (*Dpn*II, NuTi Capture-C) and low-resolution 3C (*Hind*III, Capture Hi-C) at calling interacting fragments (mm9, chr2:132,242,416-132,695,297) in erythroid cells. Tracks in order: UCSC gene annotation, *cis*-normalized mean interactions per *Dpn*II fragment using NuTi Capture-C (n=3), NuTi Capture-C viewpoints, peaky Marginal Posterior Probability of Contact (MPPC) scores with fragments with MPPC ≥0.01 darker, GenoSTAN open chromatin classification, windowed mean interactions using NuTi Capture-C, total supporting reads per *Hind*III fragment with CHi-C (n=2; co-targeted fragments are lighter in colour), CHi-C bait fragments, loops between reported significantly interacting fragments (co-targeting loops are coloured grey), erythroid tracks for open chromatin (DNaseI), promoters (H3K4me3), active transcription (H3K27ac), enhancers (H3K4me1), and boundaries (CTCF). Note overlapping MPPC signals appear darker in colour.

**Fig. 20.**
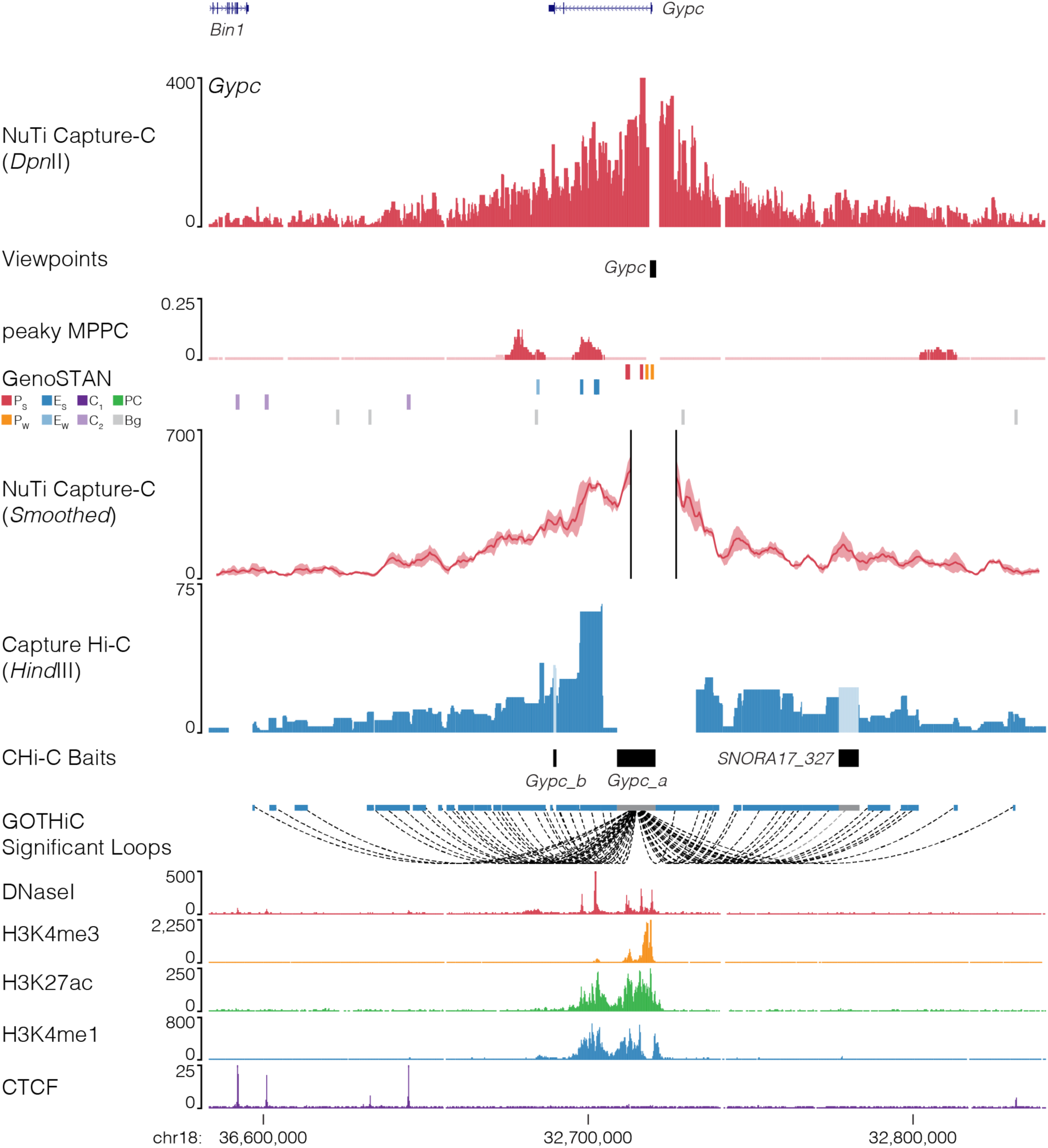
NuTi Capture-C from the *Gypc* promoter. Sequence tracks showing the difference between high-resolution 3C (*Dpn*II, NuTi Capture-C) and low-resolution 3C (*Hind*III, Capture Hi-C) at calling interacting fragments (mm9, chr18:32,583,205-32,841,048) in erythroid cells. Tracks in order: UCSC gene annotation, *cis*-normalized mean interactions per *Dpn*II fragment using NuTi Capture-C (n=3), NuTi Capture-C viewpoints, peaky Marginal Posterior Probability of Contact (MPPC) scores with fragments with MPPC ≥ 0.01 darker, GenoSTAN open chromatin classification, windowed mean interactions using NuTi Capture-C, total supporting reads per *Hind*III fragment with CHi-C (n=2; co-targeted fragments are lighter in colour), CHi-C bait fragments, loops between reported significantly interacting fragments (co-targeting loops are coloured grey), erythroid tracks for open chromatin (DNaseI), promoters (H3K4me3), active transcription (H3K27ac), enhancers (H3K4me1), and boundaries (CTCF). Note overlapping MPPC signals appear darker in colour.

**Fig. 21.**
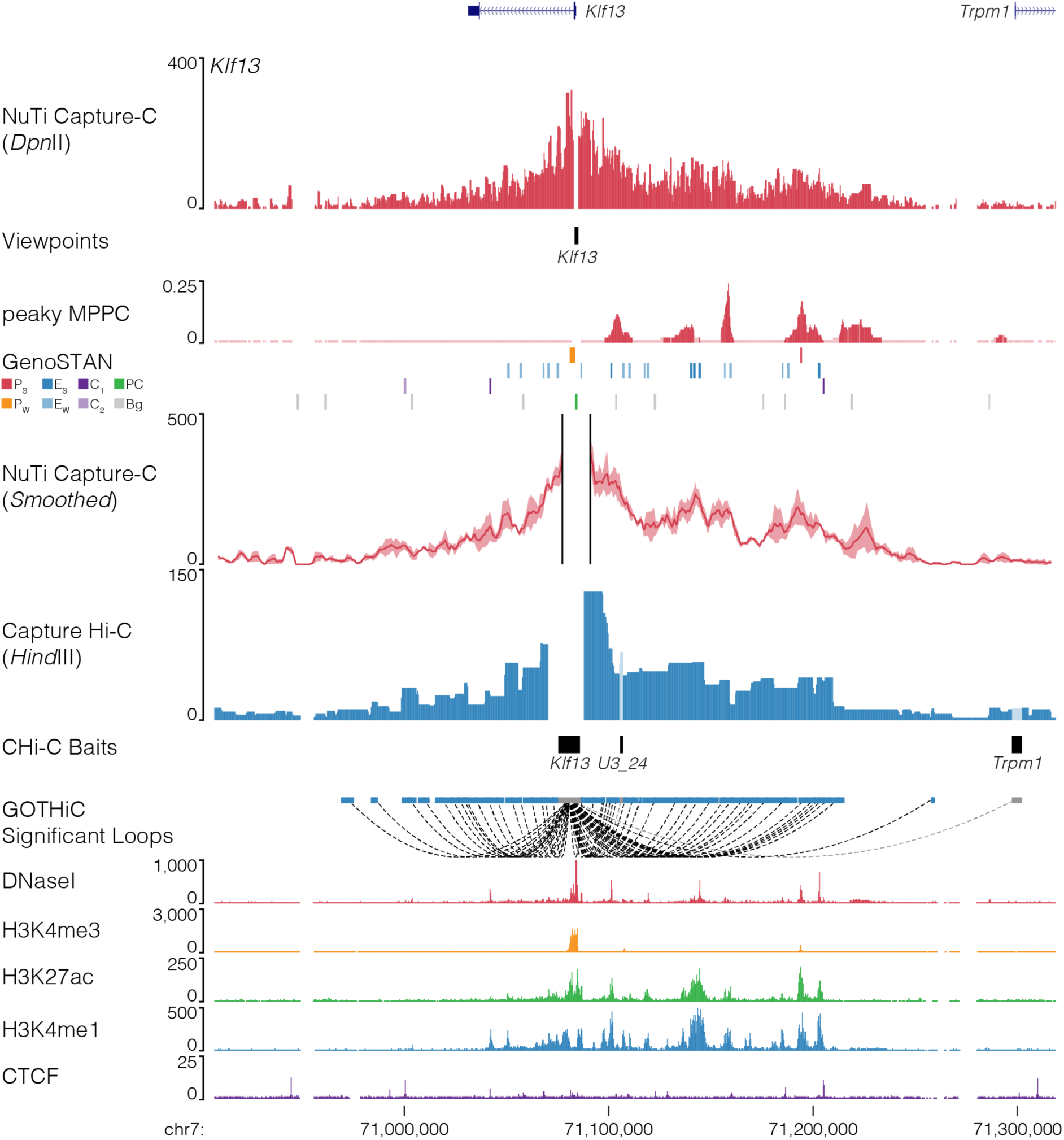
NuTi Capture-C from the *Klf13* promoters. Sequence tracks showing the difference between high-resolution 3C (*Dpn*II, NuTi Capture-C) and low-resolution 3C (*Hind*III, Capture Hi-C) at calling interacting fragments (mm9, chr7:70,906,844-71,318,843) in erythroid cells. Tracks in order: UCSC gene annotation, *cis*-normalized mean interactions per *Dpn*II fragment using NuTi Capture-C (n=3), NuTi Capture-C viewpoints, peaky Marginal Posterior Probability of Contact (MPPC) scores with fragments with MPPC ≥0.01 darker, GenoSTAN open chromatin classification, windowed mean interactions using NuTi Capture-C, total supporting reads per *Hind*III fragment with CHi-C (n=2; co-targeted fragments are lighter in colour), CHi-C bait fragments, loops between reported significantly interacting fragments (co-targeting loops are coloured grey), erythroid tracks for open chromatin (DNaseI), promoters (H3K4me3), active transcription (H3K27ac), enhancers (H3K4me1), and boundaries (CTCF). Note overlapping MPPC signals appear darker in colour.

**Fig. 22.**
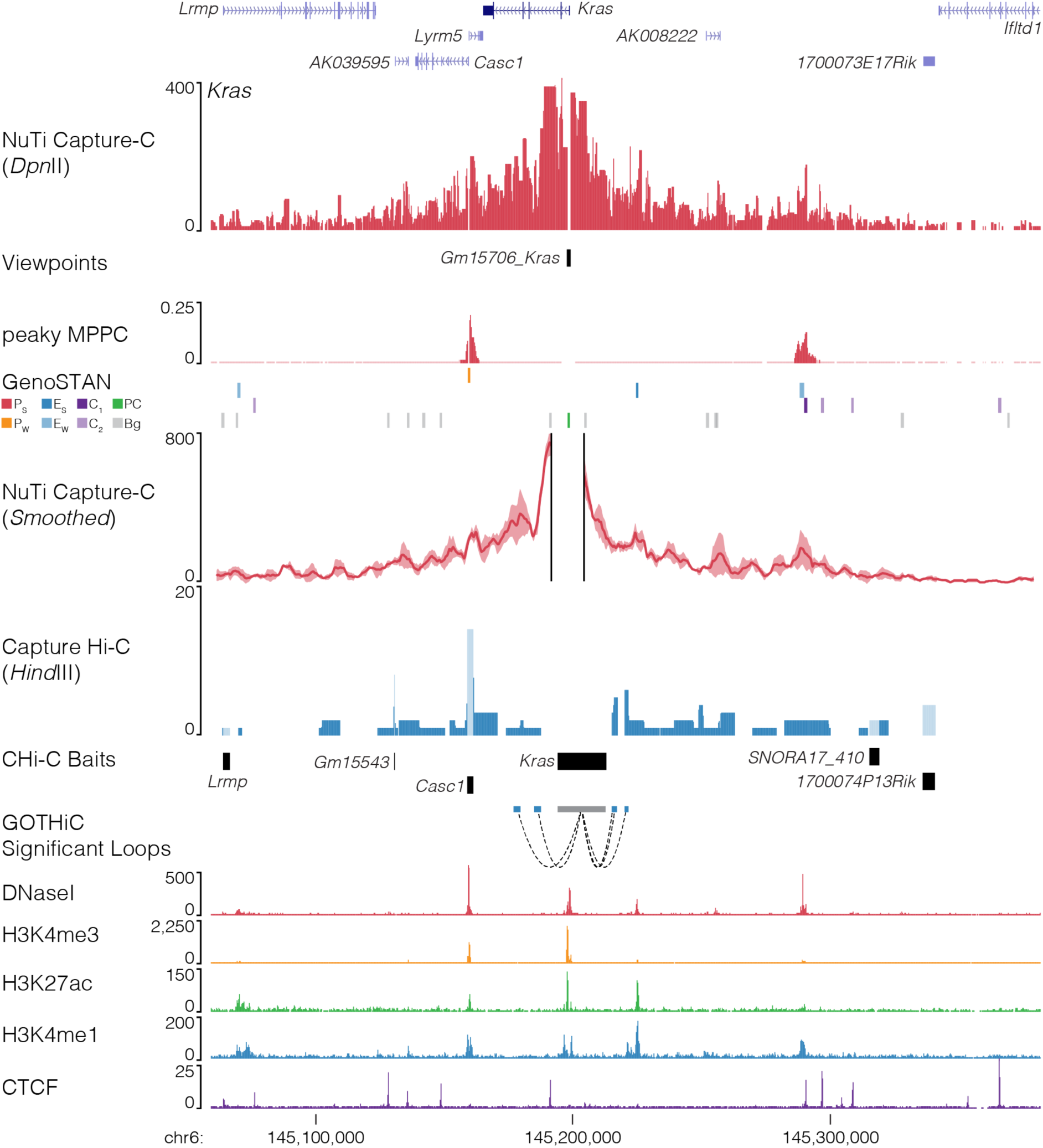
NuTi Capture-C from the *Kras* promoters. Sequence tracks showing the difference between high-resolution 3C (*Dpn*II, NuTi Capture-C) and low-resolution 3C (*Hind*III, Capture Hi-C) at calling interacting fragments (mm9, chr6:145,059,451-145,381,678) in erythroid cells. Tracks in order: UCSC gene annotation, *cis*-normalized mean interactions per *Dpn*II fragment using NuTi Capture-C (n=3), NuTi Capture-C viewpoints, peaky Marginal Posterior Probability of Contact (MPPC) scores with fragments with MPPC ≥0.01 darker, GenoSTAN open chromatin classification, windowed mean interactions using NuTi Capture-C, total supporting reads per *Hind*III fragment with CHi-C (n=2; co-targeted fragments are lighter in colour), CHi-C bait fragments, loops between reported significantly interacting fragments (co-targeting loops are coloured grey), erythroid tracks for open chromatin (DNaseI), promoters (H3K4me3), active transcription (H3K27ac), enhancers (H3K4me1), and boundaries (CTCF). Note overlapping MPPC signals appear darker in colour.

**Fig. 23.**
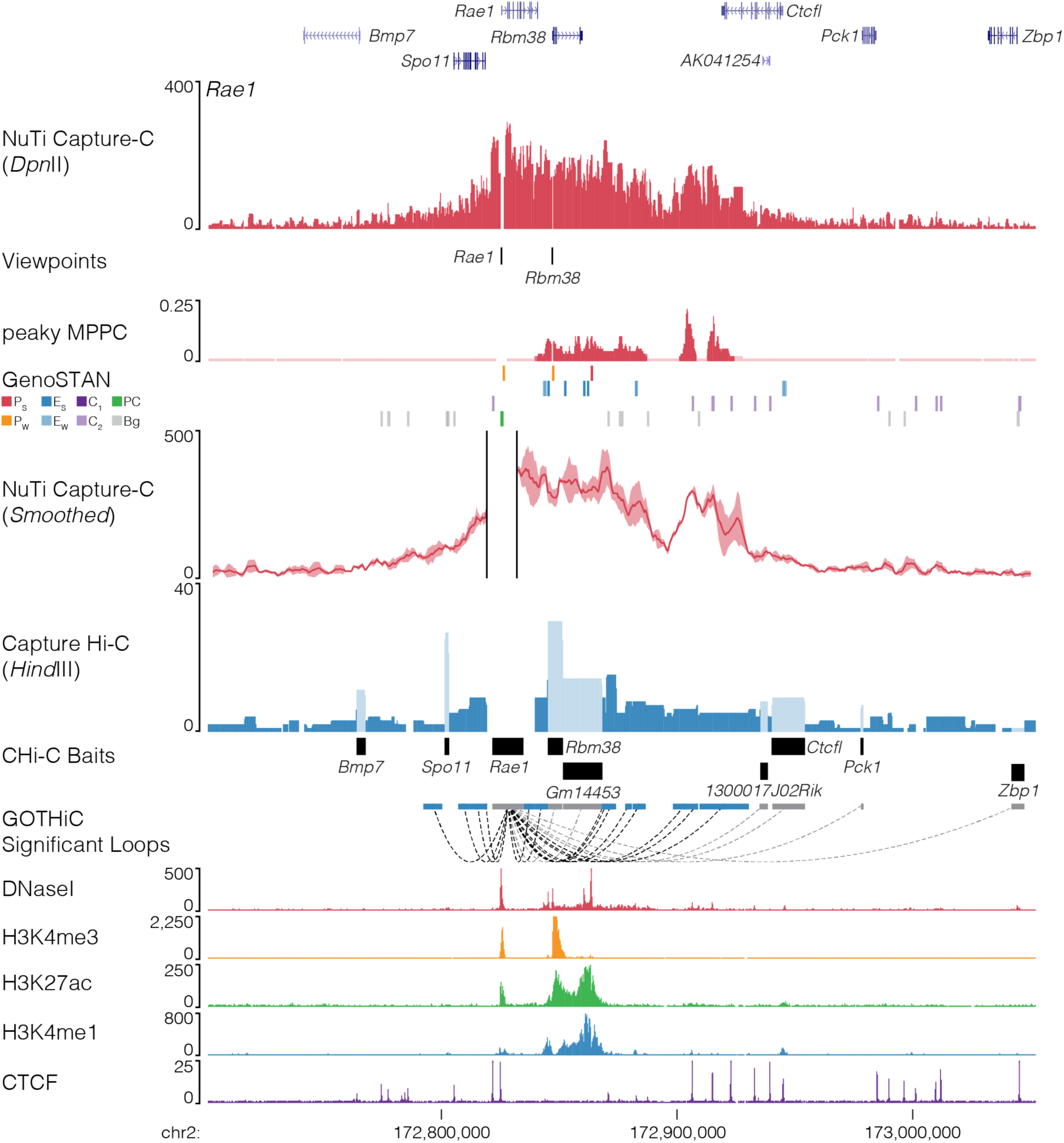
NuTi Capture-C from the *Rae1* promoter. Sequence tracks showing the difference between high-resolution 3C (*Dpn*II, NuTi Capture-C) and low-resolution 3C (*Hind*III, Capture Hi-C) at calling interacting fragments (mm9, chr2:172,701,139-173,052,278) in erythroid cells. Tracks in order: UCSC gene annotation, *cis*-normalized mean interactions per *Dpn*II fragment using NuTi Capture-C (n=3), NuTi Capture-C viewpoints, peaky Marginal Posterior Probability of Contact (MPPC) scores with fragments with MPPC ≥0.01 darker, GenoSTAN open chromatin classification, windowed mean interactions using NuTi Capture-C, total supporting reads per *Hind*III fragment with CHi-C (n=2; co-targeted fragments are lighter in colour), CHi-C bait fragments, loops between reported significantly interacting fragments (co-targeting loops are coloured grey), erythroid tracks for open chromatin (DNaseI), promoters (H3K4me3), active transcription (H3K27ac), enhancers (H3K4me1), and boundaries (CTCF). Note overlapping MPPC signals appear darker in colour.

**Fig. 24.**
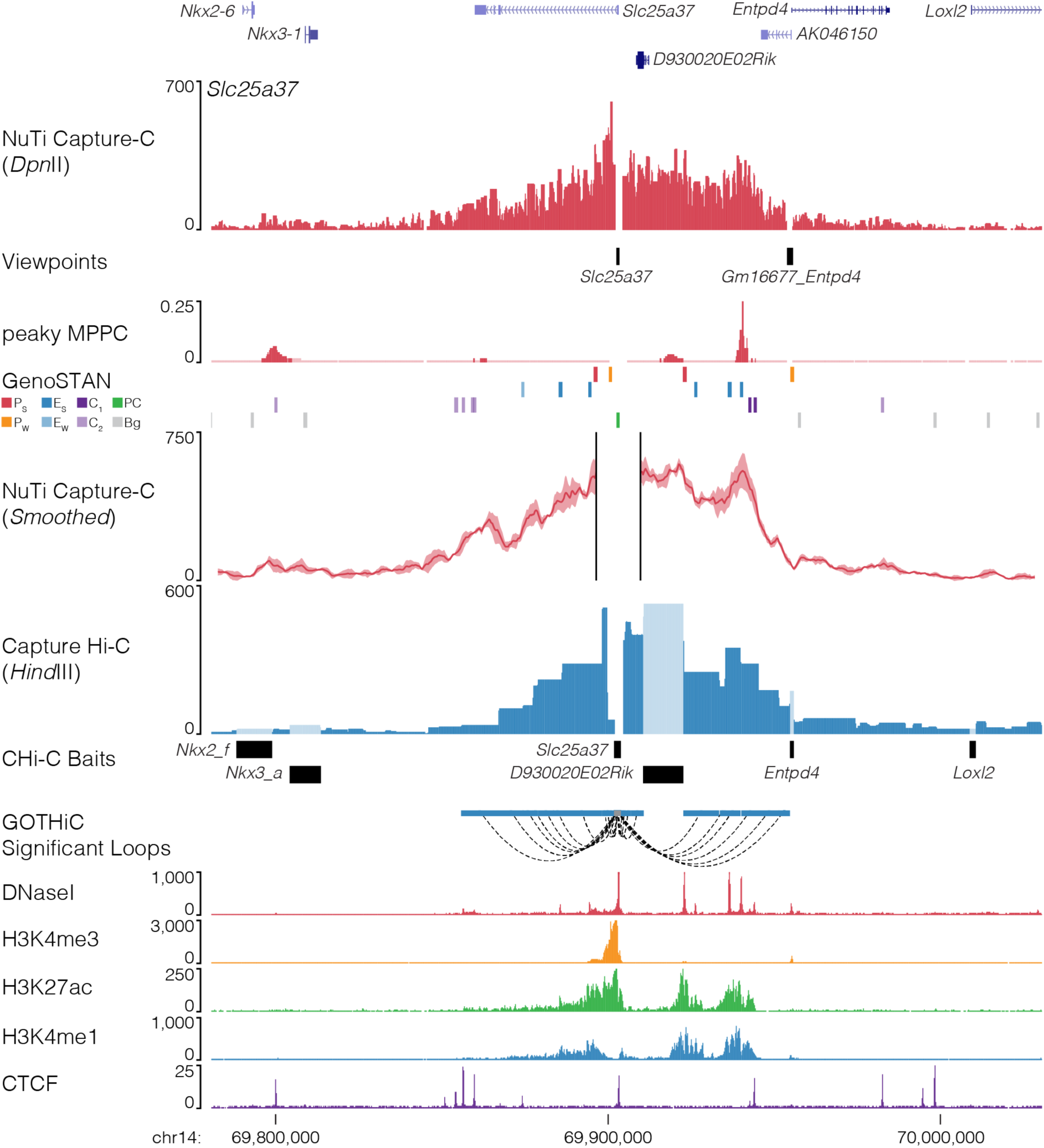
NuTi Capture-C from the *Slc25a37* promoter. Sequence tracks showing the difference between high-resolution 3C (*Dpn*II, NuTi Capture-C) and low-resolution 3C (*Hind*III, Capture Hi-C) at calling interacting fragments (mm9, chr14:69,780,624-70,030,623) in erythroid cells. Tracks in order: UCSC gene annotation, *cis*-normalized mean interactions per *Dpn*II fragment using NuTi Capture-C (n=3), NuTi Capture-C viewpoints, peaky Marginal Posterior Probability of Contact (MPPC) scores with fragments with MPPC ≥0.01 darker, GenoSTAN open chromatin classification, windowed mean interactions using NuTi Capture-C, total supporting reads per *Hind*III fragment with CHi-C (n=2; co-targeted fragments are lighter in colour), CHi-C bait fragments, loops between reported significantly interacting fragments (co-targeting loops are coloured grey), erythroid tracks for open chromatin (DNaseI), promoters (H3K4me3), active transcription (H3K27ac), enhancers (H3K4me1), and boundaries (CTCF). Note overlapping MPPC signals appear darker in colour.

**Fig. 25.**
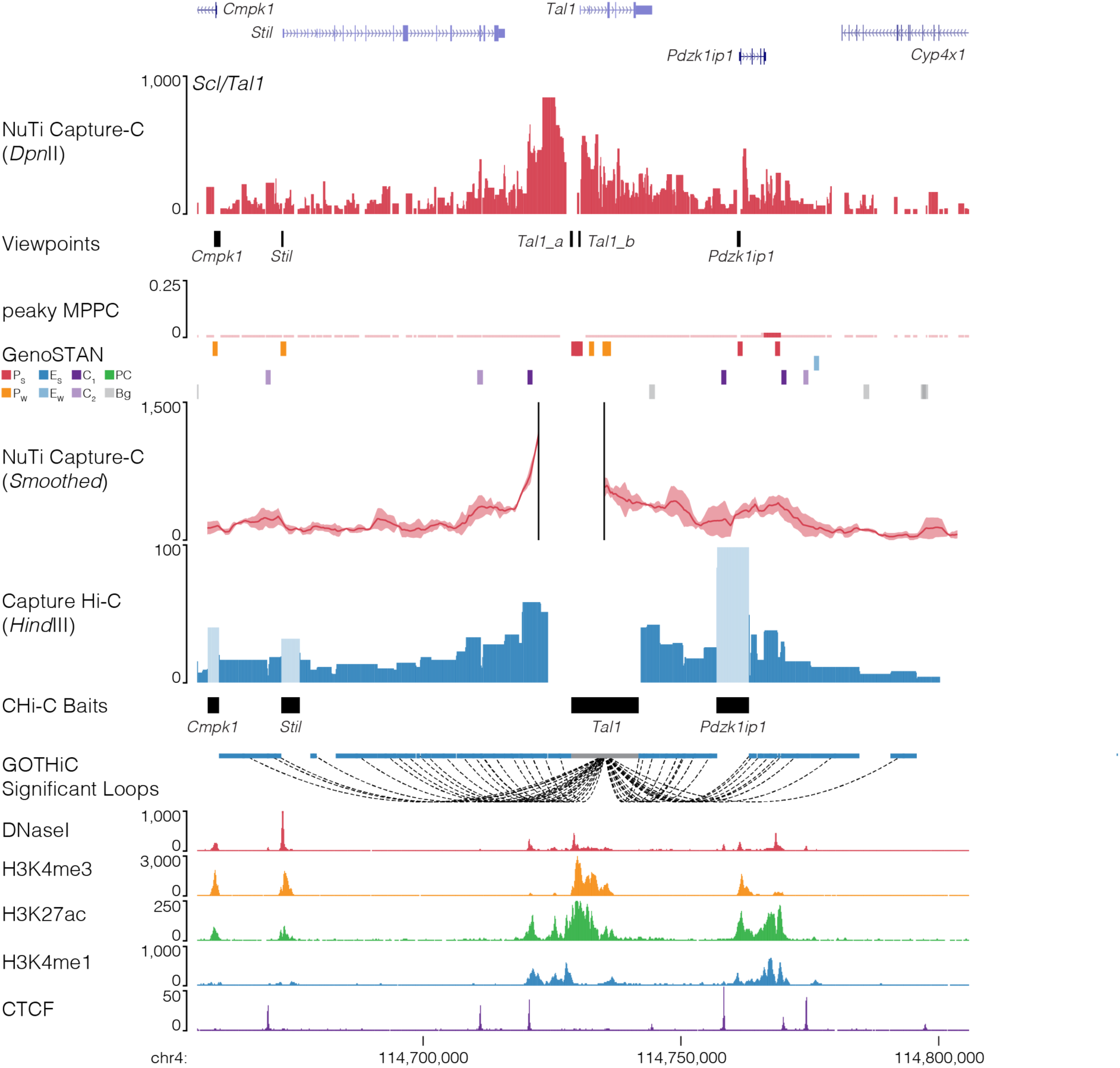
NuTi Capture-C from the *Tal1* promoter. Sequence tracks showing the difference between high-resolution 3C (*Dpn*II, NuTi Capture-C) and low-resolution 3C (*Hind*III, Capture Hi-C) at calling interacting fragments (mm9, chr4:114,656,021-114,806,020) in erythroid cells. Tracks in order: UCSC gene annotation, *cis*-normalized mean interactions per *Dpn*II fragment using NuTi Capture-C (n=3), NuTi Capture-C viewpoints, peaky Marginal Posterior Probability of Contact (MPPC) scores with fragments with MPPC ≥0.01 darker, GenoSTAN open chromatin classification, windowed mean interactions using NuTi Capture-C, total supporting reads per *Hind*III fragment with CHi-C (n=2; co-targeted fragments are lighter in colour), CHi-C bait fragments, loops between reported significantly interacting fragments (co-targeting loops are coloured grey), erythroid tracks for open chromatin (DNaseI), promoters (H3K4me3), active transcription (H3K27ac), enhancers (H3K4me1), and boundaries (CTCF). Note overlapping MPPC signals appear darker in colour.

**Fig. 26.**
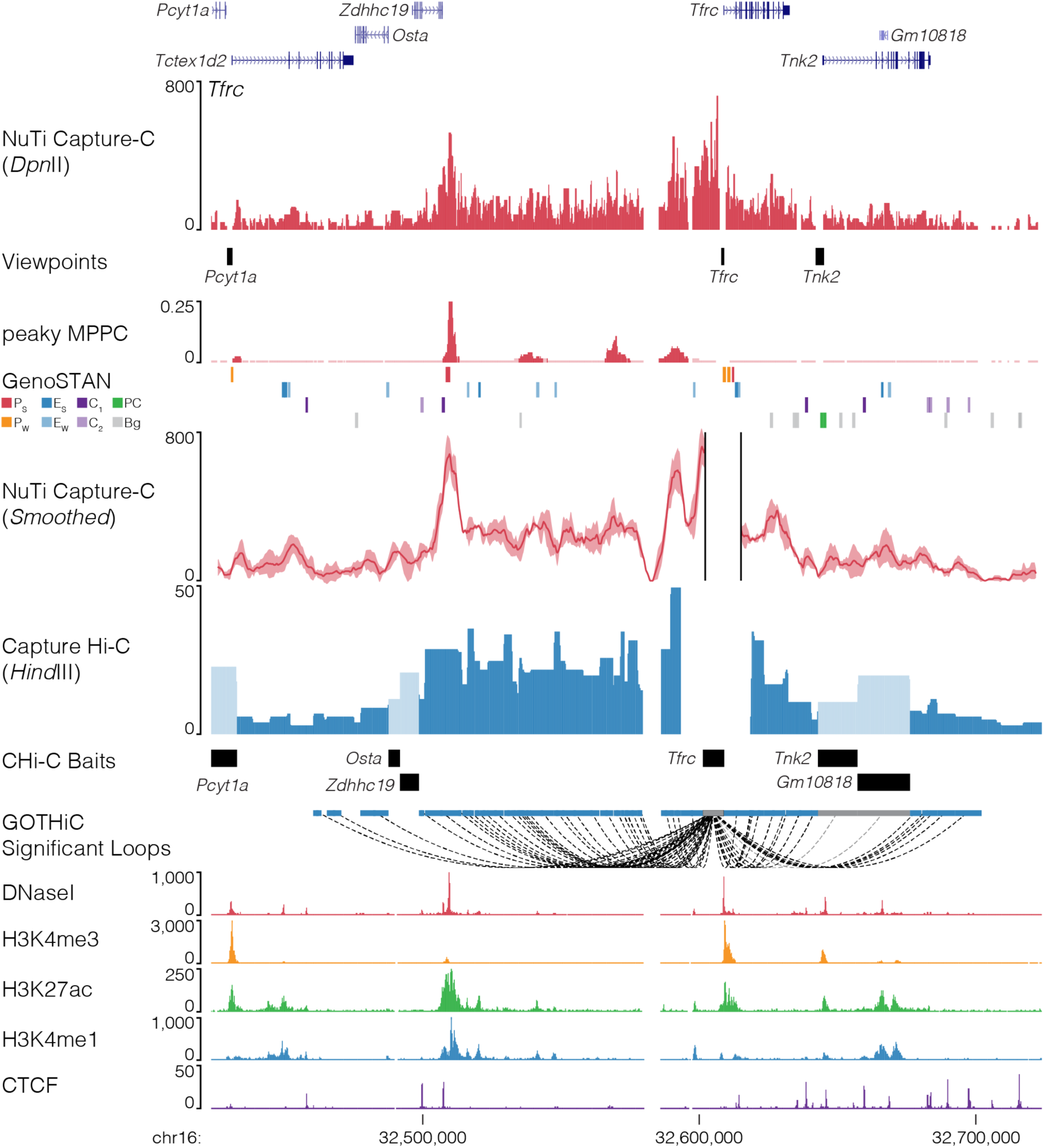
NuTi Capture-C from the *Tfrc* promoter. Sequence tracks showing the difference between high-resolution 3C (*Dpn*II, NuTi Capture-C) and low-resolution 3C (*Hind*III, Capture Hi-C) at calling interacting fragments (mm9, chr16:32,423,792-32,723,792) in erythroid cells. Tracks in order: UCSC gene annotation, *cis*-normalized mean interactions per *Dpn*II fragment using NuTi Capture-C (n=3), NuTi Capture-C viewpoints, peaky Marginal Posterior Probability of Contact (MPPC) scores with fragments with MPPC ≥0.01 darker, GenoSTAN open chromatin classification, windowed mean interactions using NuTi Capture-C, total supporting reads per *Hind*III fragment with CHi-C (n=2; co-targeted fragments are lighter in colour), CHi-C bait fragments, loops between reported significantly interacting fragments (co-targeting loops are coloured grey), erythroid tracks for open chromatin (DNaseI), promoters (H3K4me3), active transcription (H3K27ac), enhancers (H3K4me1), and boundaries (CTCF). Note overlapping MPPC signals appear darker in colour.

**Fig. 27.**
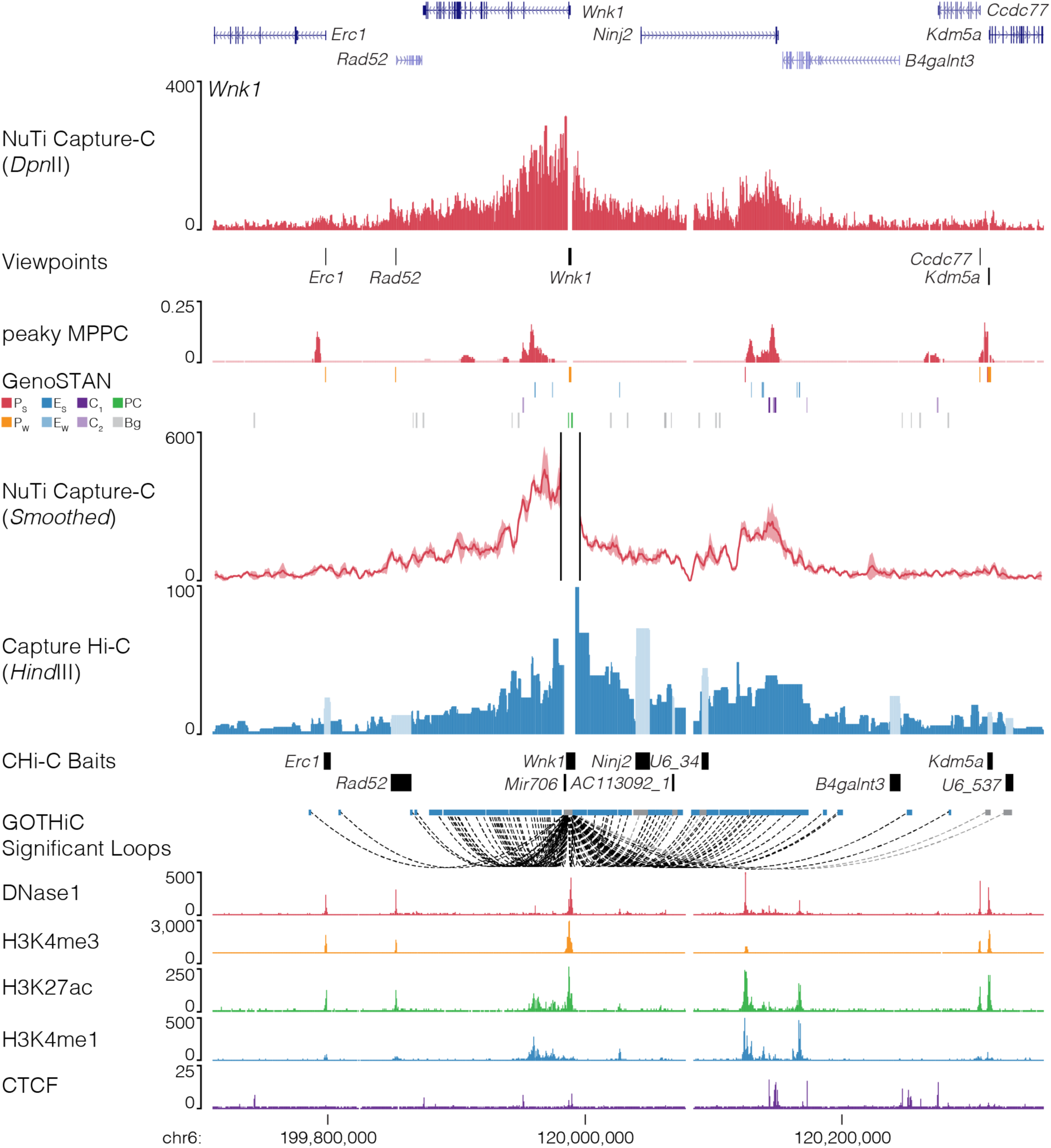
NuTi Capture-C from the *Wnk1* promoter. Sequence tracks showing the difference between high-resolution 3C (*Dpn*II, NuTi Capture-C) and low-resolution 3C (*Hind*III, Capture Hi-C) at calling interacting fragments (mm9, chr6:119,710,118-120,356,868) in erythroid cells. Tracks in order: UCSC gene annotation, *cis*-normalized mean interactions per *Dpn*II fragment using NuTi Capture-C (n=3), NuTi Capture-C viewpoints, peaky Marginal Posterior Probability of Contact (MPPC) scores with fragments with MPPC ≥0.01 darker, GenoSTAN open chromatin classification, windowed mean interactions using NuTi Capture-C, total supporting reads per *Hind*III fragment with CHi-C (n=2; co-targeted fragments are lighter in colour), CHi-C bait fragments, loops between reported significantly interacting fragments (co-targeting loops are coloured grey), erythroid tracks for open chromatin (DNaseI), promoters (H3K4me3), active transcription (H3K27ac), enhancers (H3K4me1), and boundaries (CTCF). Note overlapping MPPC signals appear darker in colour.

**Fig. 28.**
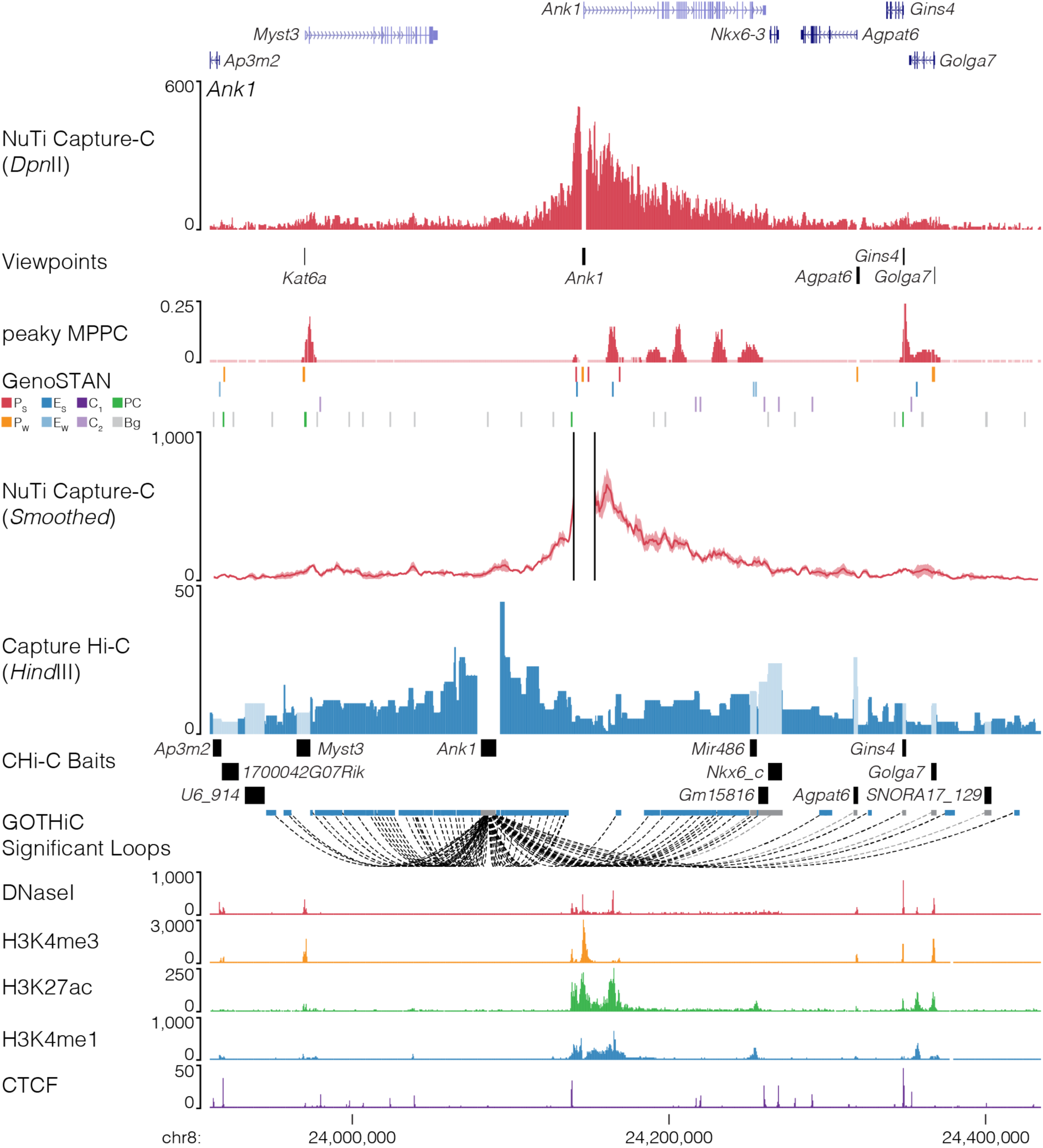
NuTi Capture-C from the *Ank1* promoter. Sequence tracks showing the importance of tissue specific probe design when performing promoter capture with either high-resolution 3C (*Dpn*II, NuTi Capture-C) or low-resolution 3C (*Hind*III, Capture Hi-C), particularly for genes with multiple promoters (mm9, chr8:23,910,000-24,435,000) in erythroid cells. Tracks in order: UCSC gene annotation, *cis*-normalized mean interactions per *Dpn*II fragment using NuTi Capture-C (n=3), NuTi Capture-C viewpoints, peaky Marginal Posterior Probability of Contact (MPPC) scores with fragments with MPPC ≥ 0.01 darker, GenoSTAN open chromatin classification, windowed mean interactions using NuTi Capture-C, total supporting reads per *Hind*III fragment with CHi-C (n=2; co-targeted fragments are lighter in colour), CHi-C bait fragments, loops between reported significantly interacting fragments (co-targeting loops are coloured grey), erythroid tracks for open chromatin (DNaseI), promoters (H3K4me3), active transcription (H3K27ac), enhancers (H3K4me1), and boundaries (CTCF). Note overlapping MPPC signals appear darker in colour.

**Fig. 29.**
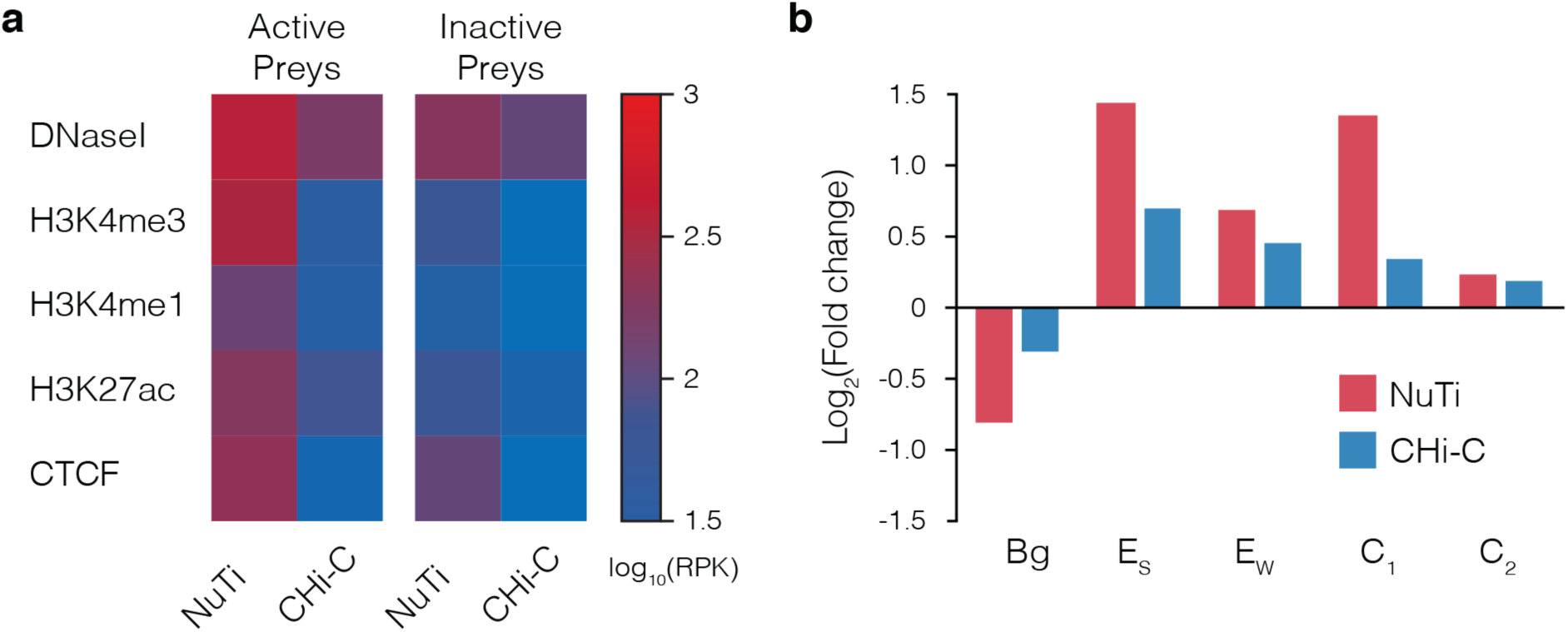
Comparison of interactions identified by NuTi Capture-C and Capture Hi-C. **a,** Average chromatin signature in mouse erythroid cells over fragments identified as being significantly interacting with either active or inactive promoters by NuTi Capture-C (Nu-3C) and Capture Hi-C (CHi-C). **b,** Enrichment of different classes of open chromatin element in fragments identified as being significantly interacting with active promoters. E_S_: Enhancer (Strong H3K27ac), E_W_: Enhancer (Weak H3K27ac), C_1_: CTCF near promoter/enhancer, C_2_: CTCF, Bg: Background, RPK: Reads per kilobase.

### SUPPLEMENTARY NOTE: Mathematical modelling of 3C enrichment bias

The number of interactions in 3C experiments are constrained by the fact each fragment can only ligate to two other fragments. Therefore, the total number of interactions is limited by the total number of cells, in effect, it is a closed system. To explore the effects of enrichment for multiple targets on 3C libraries we created a small closed system of fragments where each fragment has 5,000 interactions. Within this system, interactions involving three fragments, “A”, “B”, and “C”, can be sampled (i.e. enriched) with varying levels of efficiency. The remaining fragments can be collectively considered “X”, with “D” being one of these remaining fragments. The absolute number of interactions between each fragment within this system and example interaction profiles are demonstrated below (Supp. Note Table 1, Supp. Note Fig. 1). To demonstrate the effect on interaction calling, within this system “significant interactions” are simply those observed at a frequency of greater than 1 in 50 (>0.02). The significant interactions in this system are A-B, A-D, and B-D.

**Supp. Note Fig. 1.**
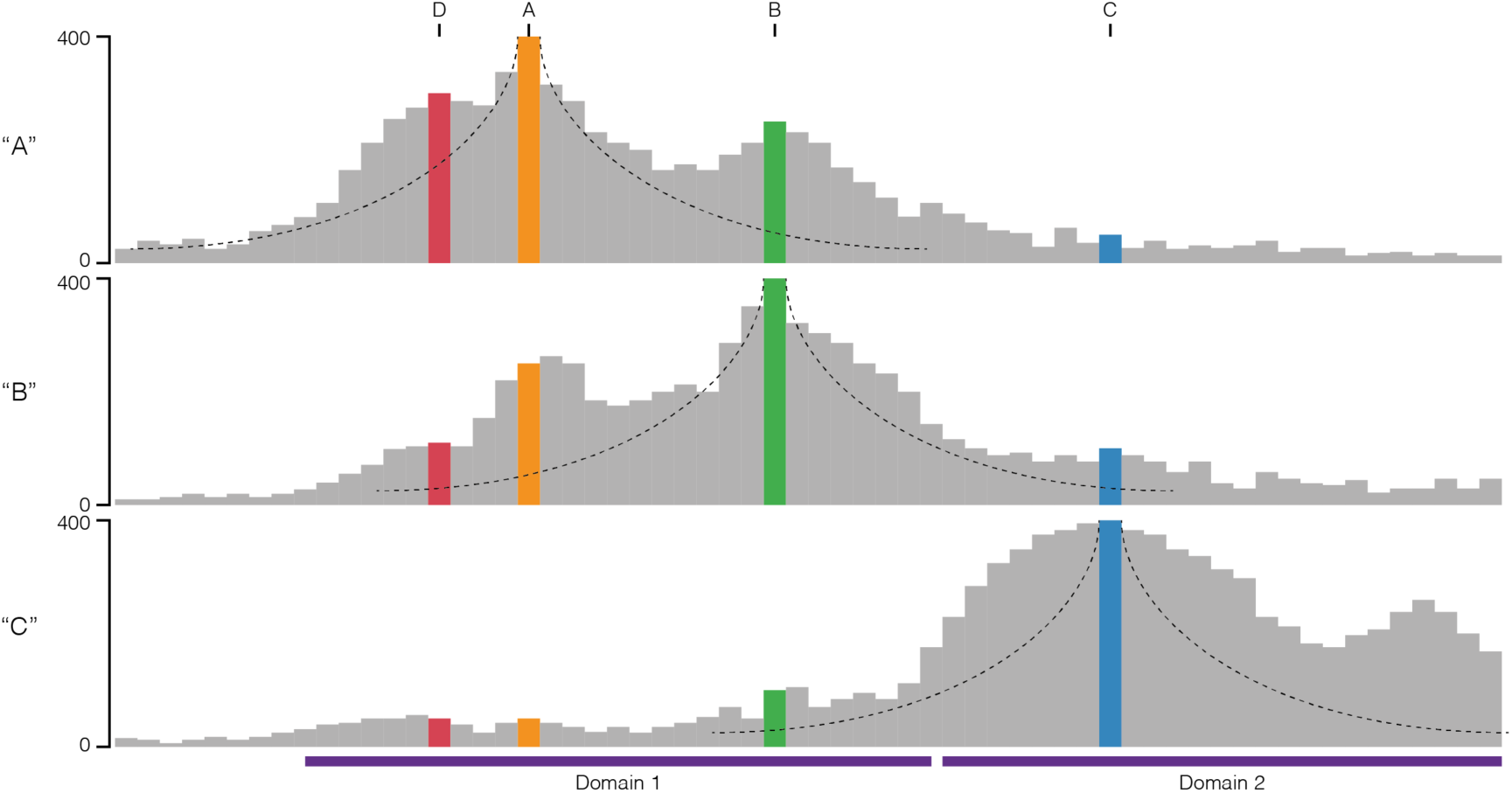
A closed system of interactions. Profile of absolute interaction counts for three sampleable fragments “A”, “B” and “C” within a closed system. The fragments separate into two interacting domains. Monotonic decay curves associated with the polymer models of interaction are shown.

**Supp. Note Table 1.**
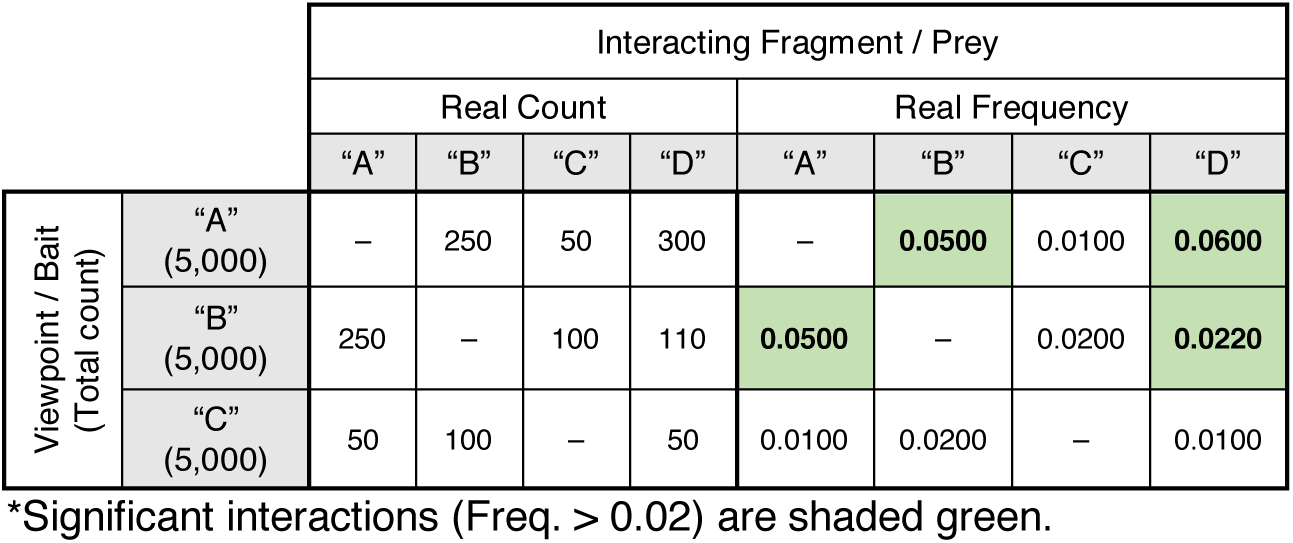
A closed system of interactions.

The sampling within this system represents the targeted enrichment of 3C methods (e.g. probe hybridisation or immunoprecipitation). For 3C enrichment, efficiency can be affected by, among other things, the number of individual probes targeting each viewpoint and the melting points (e.g. Capture-C, CHi-C), and the level of target signal (e.g. Hi-ChIP, Hi-ChIRP, ChIA-PET). To represent this diversity in these processes, we assigned sampling of “A” and “C” to be highly efficient at 80% and 90% respectively. Whereas enrichment of “B” is relatively low at 10%. When each of these sampling efficiencies is applied to any one fragment at time, the total number of observed interactions decreases, however the frequency of interaction within the system remains constant (Note Fig. 2, Note Table 2). At this point, it is important to note that while the B-to-C count is lower than the C-to-B count, their proportional frequencies are still equal, and the same set of “significantly interacting” fragments are detected.

**Supp. Note Fig. 2.**
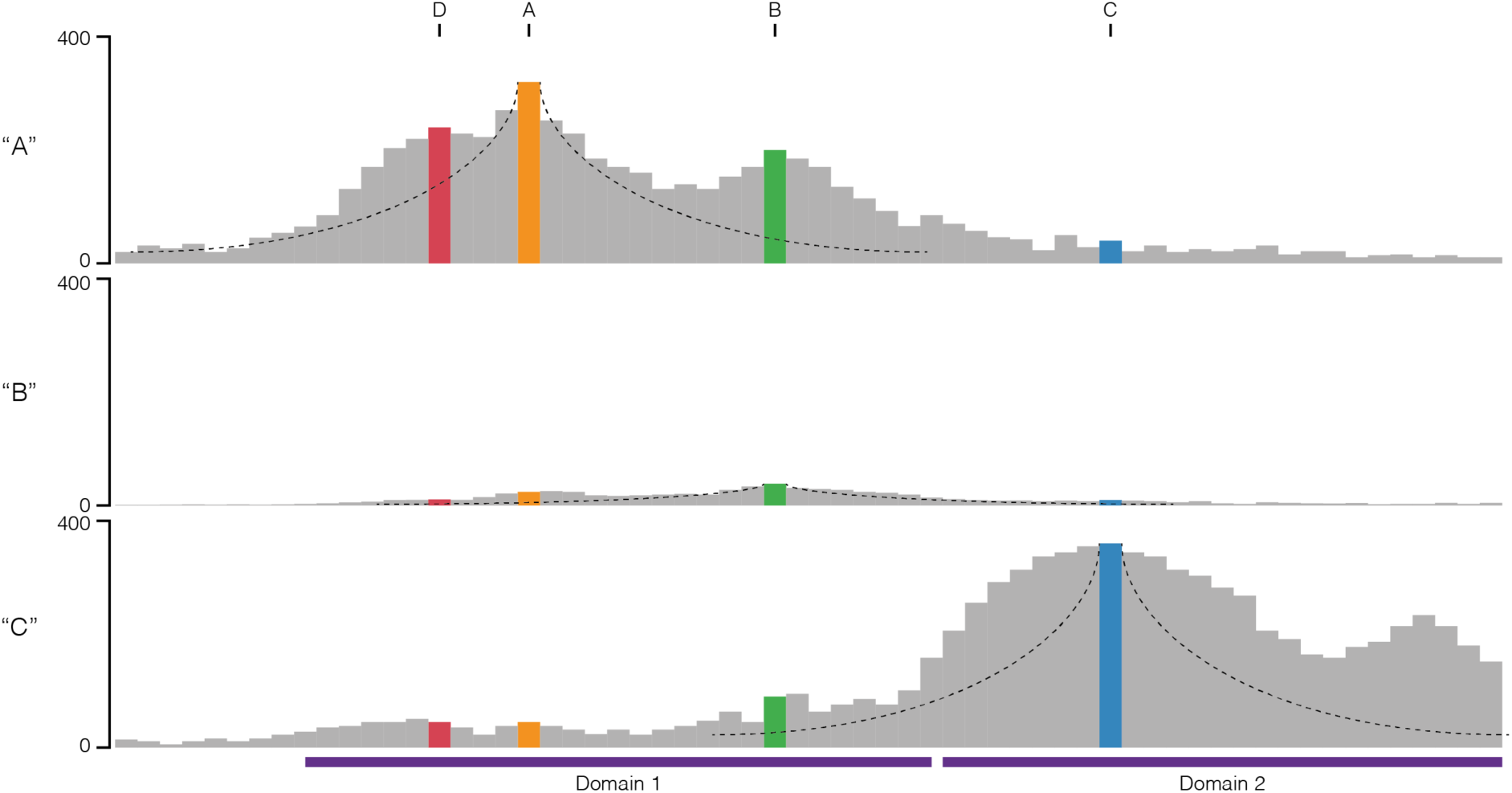
Independent sampling of interactions. Profile of observed interaction counts for three independently sampled fragments “A”, “B” and “C” within a closed system.

**Supp. Note Table 2.**
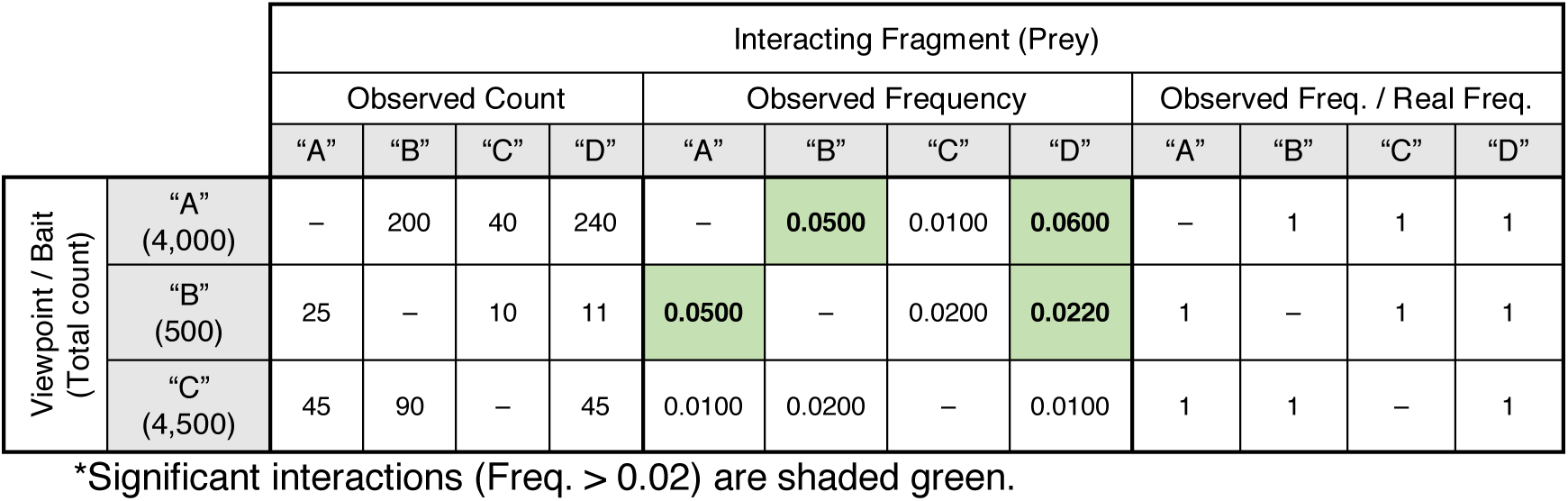
Interaction counts following independent sampling.

When we consider sampling fragment “A”, at 80% efficiency, we fail to see 50 “A-B” interactions, ten “A-C” interactions, and 60 “A-D” interactions. If we were to simultaneously sample (co-sample) the remaining viewpoints, from the missed interactions would can recover five “A-B” interactions (at 10% “B” sampling efficiency), nine “A-C” interactions (at 90% “B” sampling efficiency), and zero “A-D” interactions as it is unsampled. This recovery can be applied to each fragment as the first or second fragment sampled and leads to as much as a 9.1-fold increase in observed interaction counts. When the co-sampled fragments are presented as frequency values significant divergence from the true values within the system is observed (Supp. Note Fig. 3, Supp. Note Table 3). This divergence in frequency varies across each interacting pair and ranges from a 0.65-fold decrease (B-D) to a near 6-fold increase (B-C).

**Supp. Note Fig. 3.**
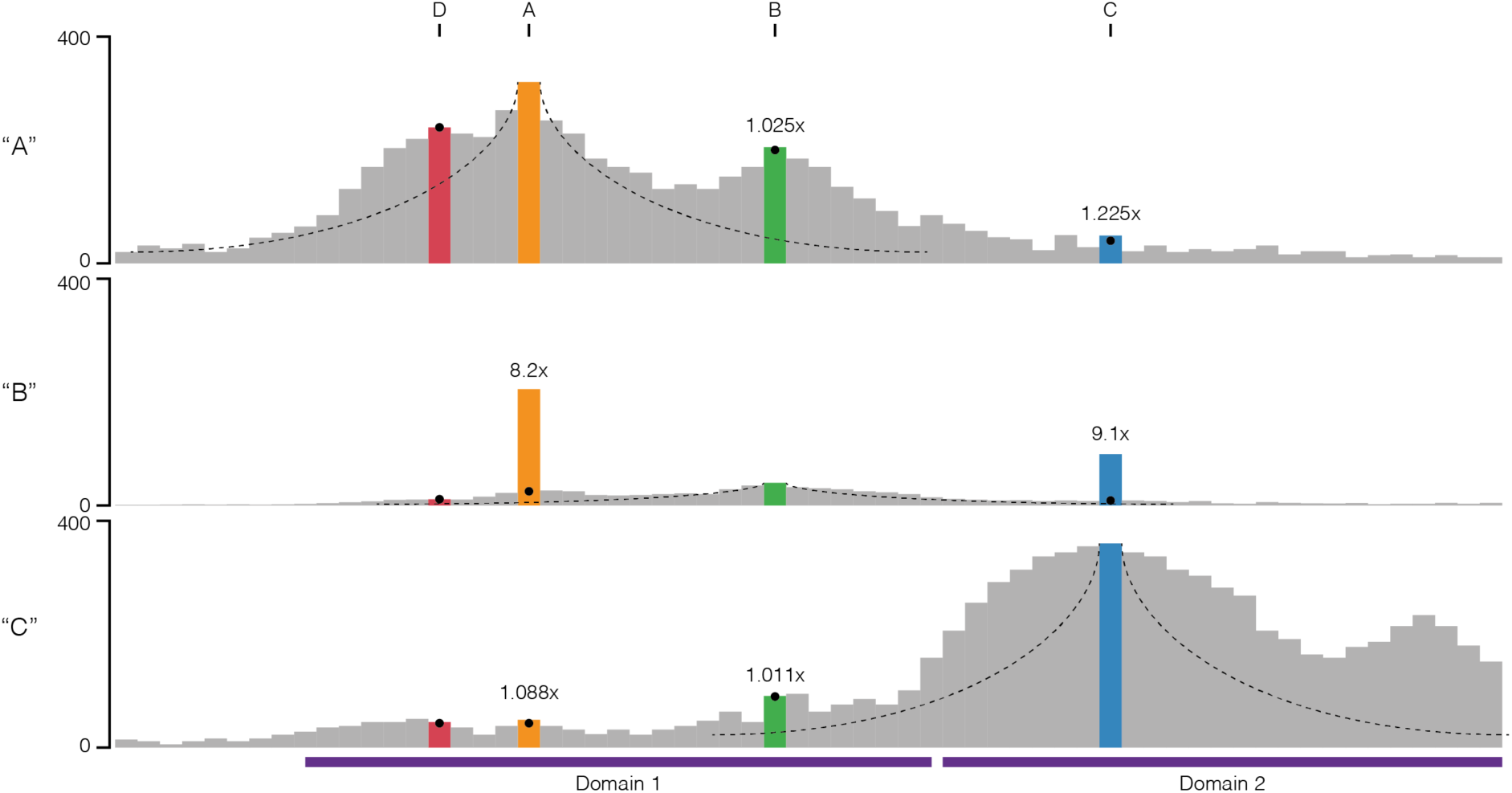
Co-sampling of interactions. Profile of observed interaction counts for three co-sampled fragments “A”, “B” and “C” within a closed system. Black circles represent counts observed with independent sampling, and the observed fold difference in raw counts is shown.

**Note Table 3.**
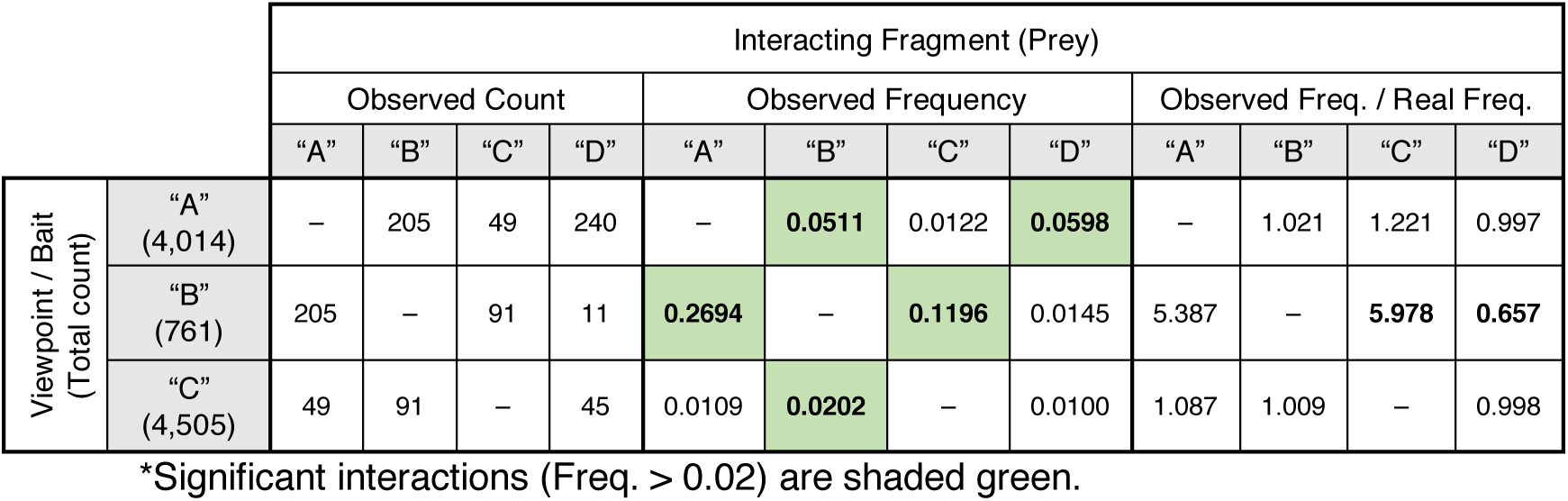
Interaction counts following co-sampling.

Interestingly, where previously the frequency of B-to-C matched C-to-B, it is now the count of interactions which is equal, and the observed frequency is unequal. This change in frequency also results in different significant interactions, B-D is no longer called (false negative), while B-C is now called (false positive). The effect on relative frequency of co-sampling is bi-directional: the observed frequency of interaction with co-sampled fragments increases, and the observed frequency of interaction with un-sampled fragments decreases. Therefore, significant and variable bias is introduced by co-targeting.

To determine the effect of this divergence we formalized this bias into a polynomial equation (Supp. Note Fig. 4) describing the observed interaction frequency of two co-targeted fragments (O_AB_) within all interactions containing “A”. For the numerator, the efficiency of targeting “A” (E_A_) and the real frequency of “A-B” (f_AB_) determines the number of interactions sampled by the A probe and the number of “lost interactions” available for the B probe (1-E_A_). These lost interactions are then recovered at the efficiency of the B probe (E_B_). The denominator, which describes the total observations involving A, can be simply denoted as E_A_ plus the number of recovered events (E_B_ x f_AB_ x [1-E_A_]). The level of bias can then be calculated as O_AB_ divided by f_AB_.

**Supp. Note Fig. 4.**
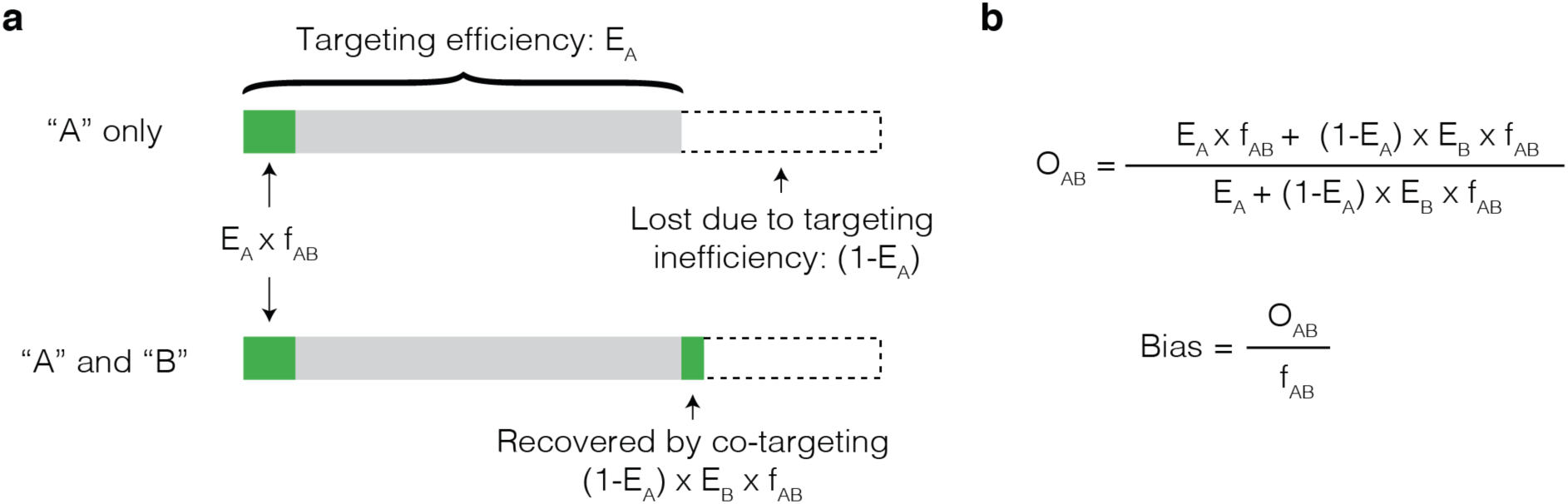
Model for the effect on observed frequency caused by co-targeting. **a,** Diagram of the total number of interactions containing A (entire bar), which includes A-B (green) and the effect of incompletely efficient targeting. Un-enriched (or lost) interactions are in dotted lines, but can be recovered by capture with additional probes. **b,** Equations for the observed frequency of A-B interaction (O_AB_) when both A and B are samples and calculation of bias.

We used this equation to model the effects of variable efficiency of enrichment (0.05-1.0), and variable underlying interaction frequencies (1/200,000, 1/25,000 and 1/40; based on the min, median, and max interactions frequencies associated with *Hba-1* capture). Under these tested parameters the highest level of bias was a ∼20-fold increase in frequency (Supp. Note Fig. 5), seen when the primary target had a low enrichment efficiency, and the secondary target had a high enrichment efficiency. Notably, for any given level of enrichment efficiency the level of bias was variable across the interaction frequencies, with infrequent interactions more affected then frequent interactions.

**Supp. Note Fig. 5.**
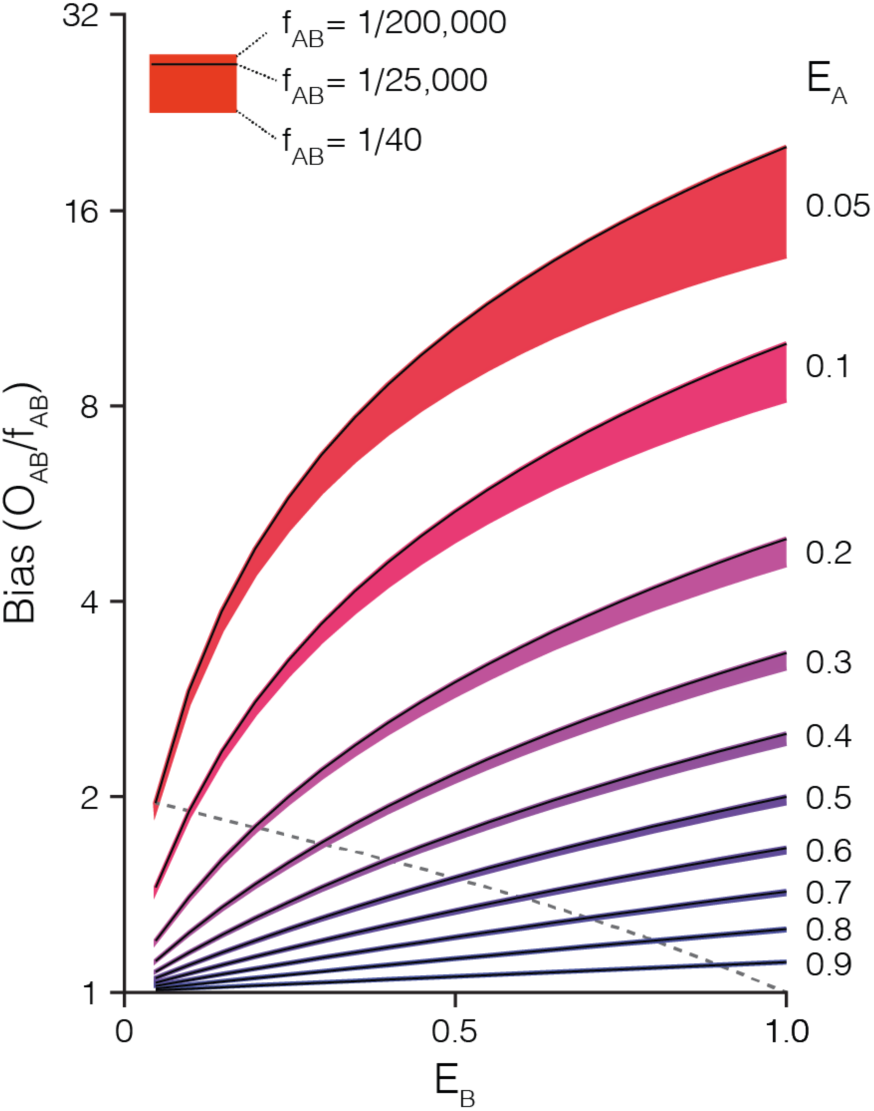
Levels of observed frequency bias caused by co-targeting. Variable levels of bias are observed in the observed frequency of A-B interaction (O_AB_) when altering the efficiency of targeting fragment A (E_A_), targeting fragment B (E_B_), and the real frequency of A-B interaction (f_AB_). The Dashed line shows when efficiency of targeting A and B is equal (E_A_ = E_B_). Note that bias is avoided only when E_A_ is equal to either zero or one.

The implication of these results are quite striking and two-fold. Firstly, when investigating a viewpoint with very low enrichment (say ChIA-PET for a poorly bound PolII site, or Hi-ChIP at a weak H3K27ac peak) then significant enrichment bias is likely to be seen at strong PolII or H3K27ac sites, regardless of whether or not they are actually interacting. In fact, the rarer an interaction is, the stronger the bias effect. Secondly, because all three parameters (enrichment at targets, and the underlying interaction frequency) contribute significantly to the observed bias, proper data correction depends upon having accurate values for all three parameters.

