## Supplementary Figures for "Targeted high-resolution chromosome conformation capture at genome-wide scale"

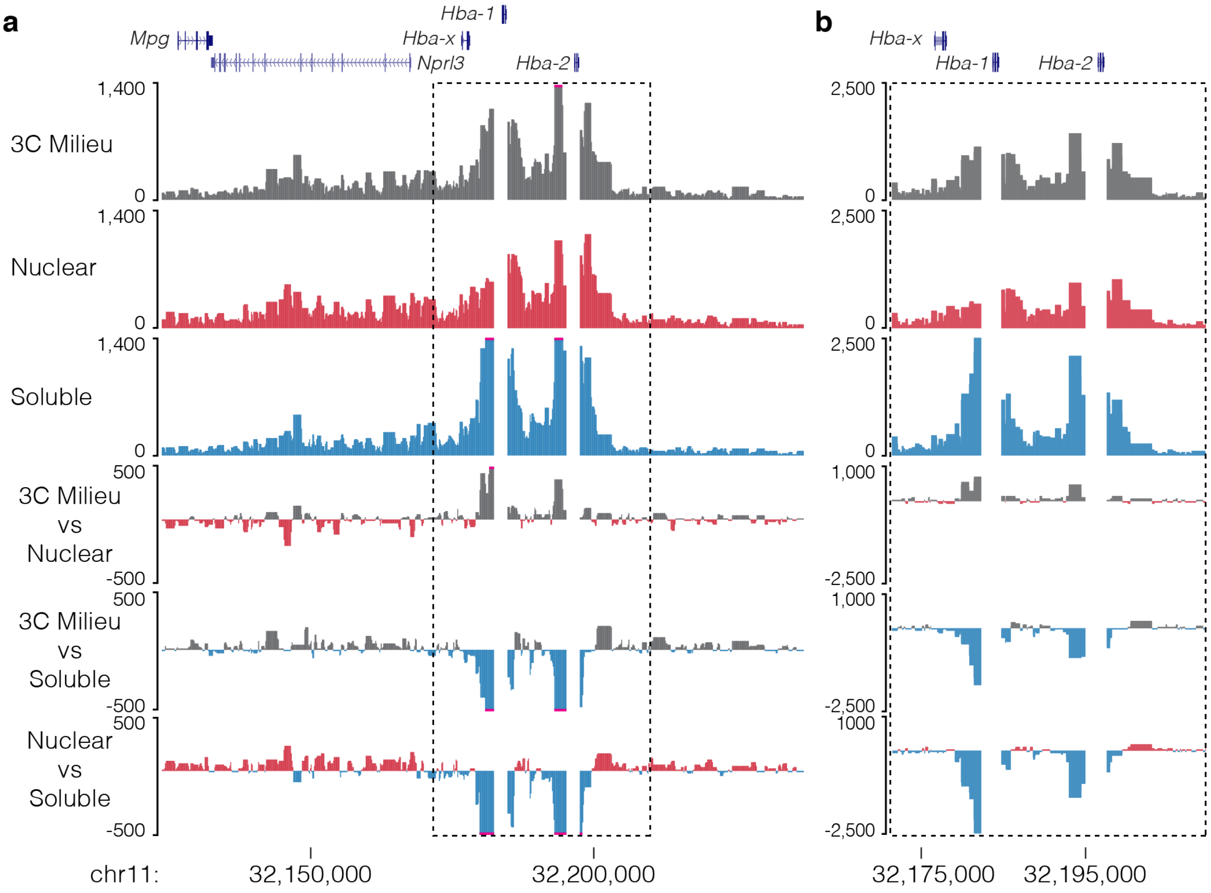


**Supp. Fig. 1 | Soluble 3C material has a higher proximity signal.** Capture profiles and comparison tracks for *Hba-1/2* capture in mouse erythroid cells **(a)** from total 3C library Milieu or its fractionated nuclear and soluble fractions shows soluble material has a higher proximity signal **(b)**, likely from small diffusing chunks of digested crosslinked chromatin.

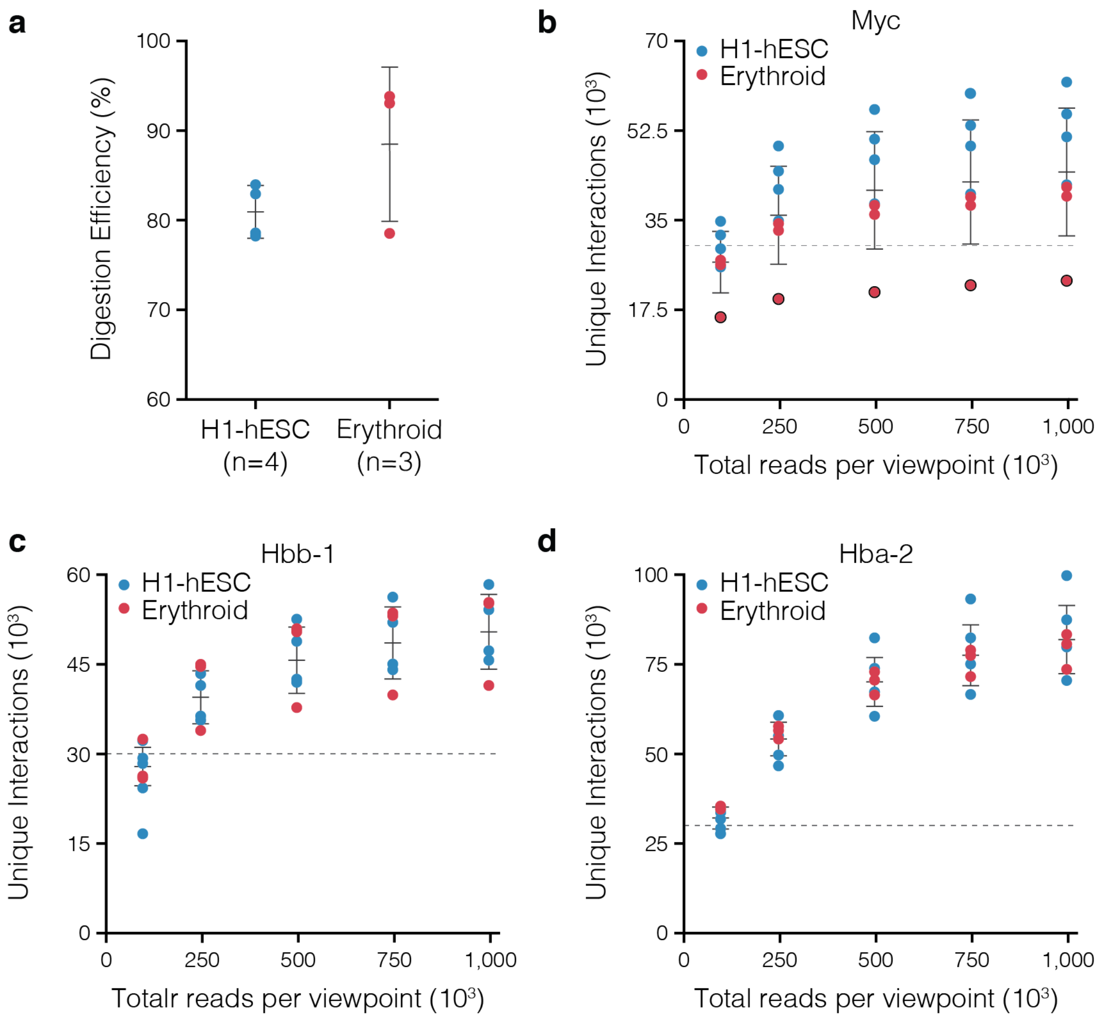


**Supp. Fig. 2 | Reporter sensitivity through sequencing depth. a,** Digestion efficiency for Nu-3C libraries from human embryonic stem cells (H1-hESC) and erythroid cells. NuTi Capture-C was performed for the seven multiplexed libraries targeting *Myc* **(b)**, *Hbb-b1/2* **(c)**, and *Hba-1/2* **(d)** and sequenced to over $10^6$ reads per viewpoint per library. Sequence files were subsampled and analyzed to determine number of unique reporters. Dashed lines represent 30,000 unique reporters, or extremely high-sensitivity capture. For *Myc*, one donor has a polymorphism that removes one of the two *Dpn*II sites on the targeted fragment (black outline) – illustrating the effect of using a single probe. Bars show mean and one standard deviation.

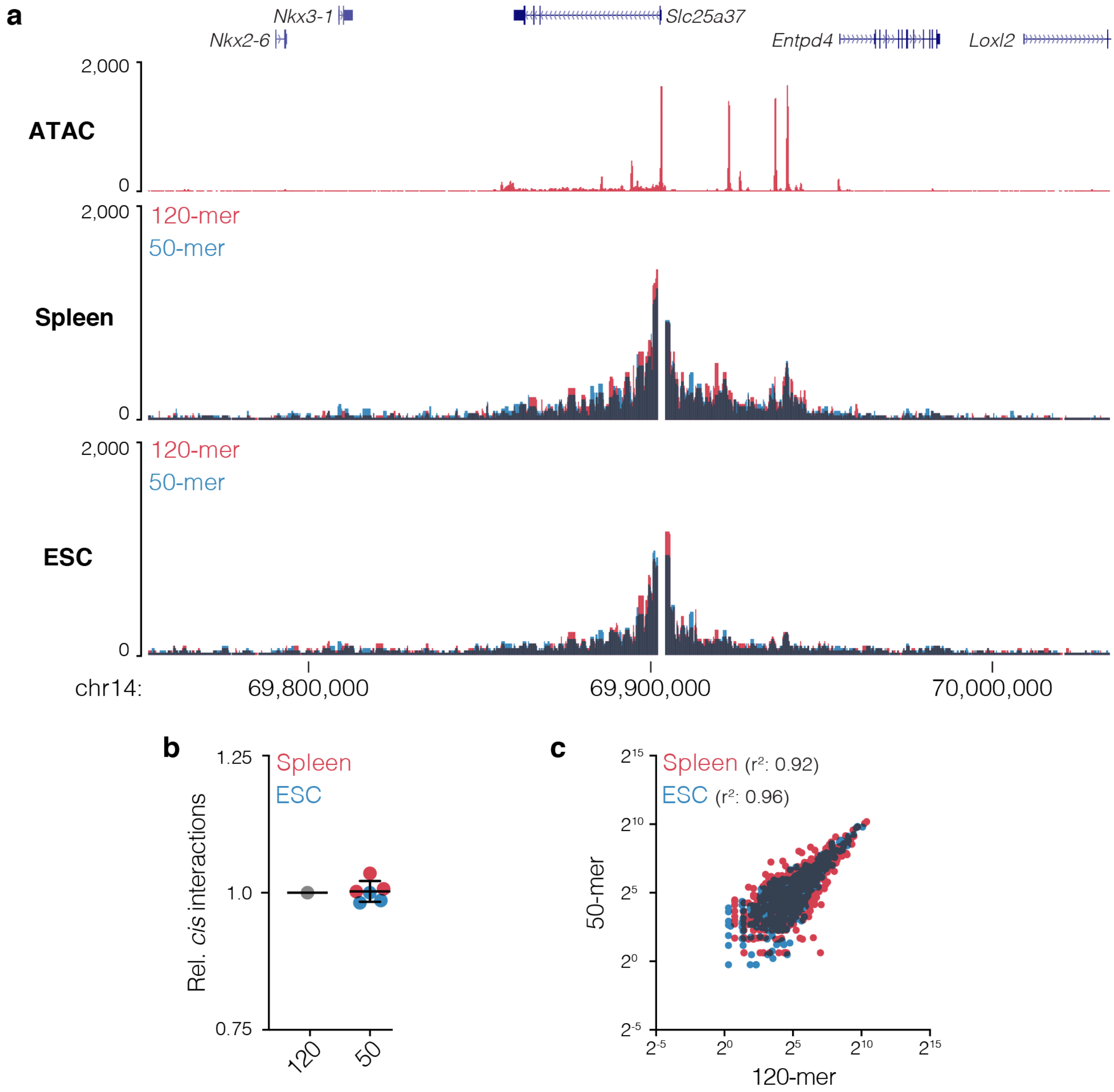


**Supp. Fig. 3 I Capture of *Slc25a37* with short probes. a,** Overlaid 3C interaction profile for *Slc25a37*, which encodes mitoferrin, from mouse erythroid (n=3) and embryonic stem cells (ESC, n=3) captured with either 120-mer or 50-mer probes. Darkened areas show overlapping signals. **b**, Number of cis reporters relative to 120-mer capture. **c**, Comparison of interactions counts from using long or short probes for fragments displayed in panel **a** with Pearson's correlation.

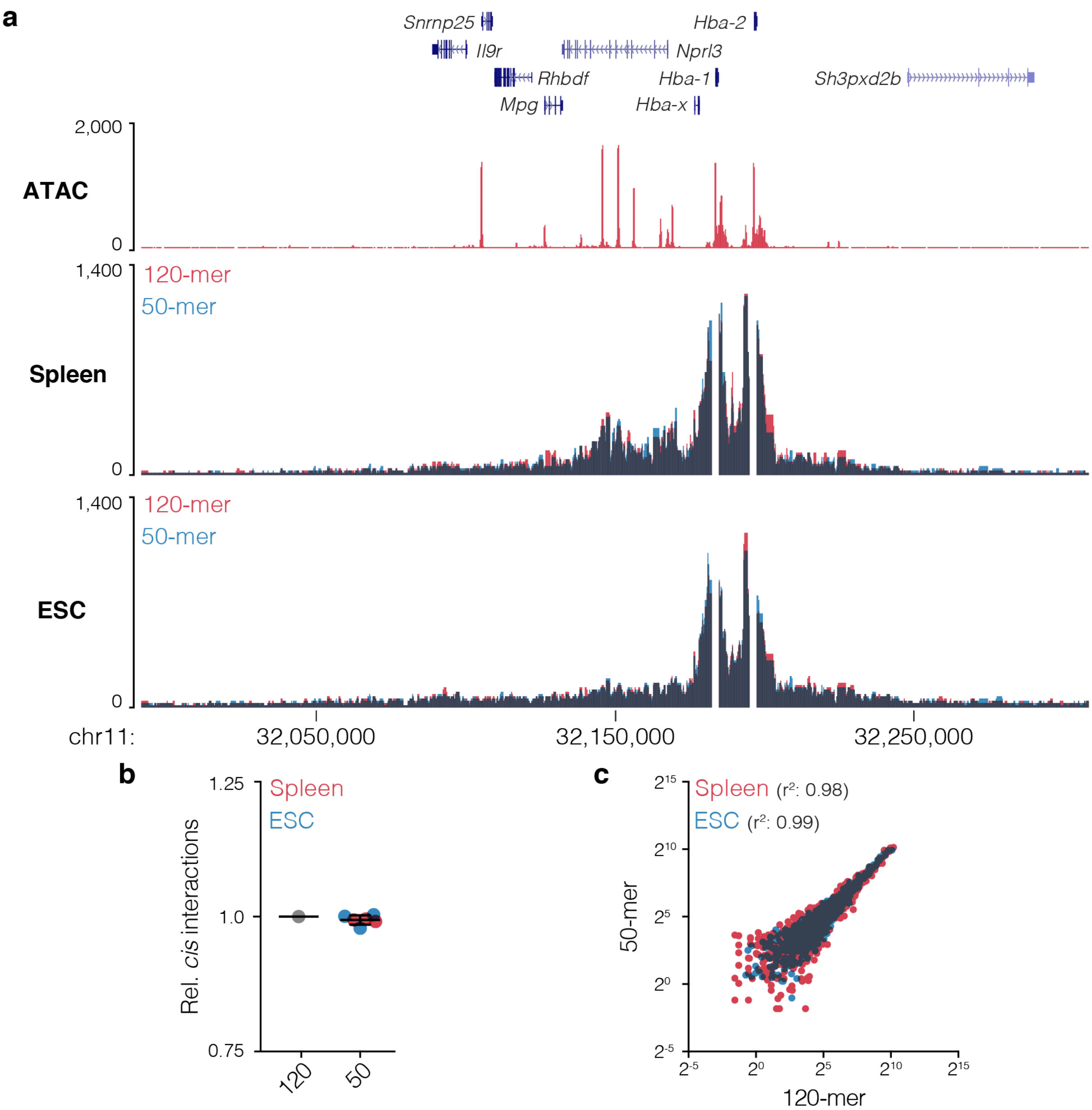


**Supp. Fig. 4 | Capture of $\alpha$-globin locus with short probes. a,** Overlaid 3C interaction profile for *Hba-1 and Hba-2*, which encode $\alpha$-globin, from mouse erythroid (n=3) and embryonic stem cells (ESC, n=3) captured with either 120-mer or 50-mer probes. Darkened areas show overlapping signals. **b**, Number of cis reporters relative to 120-mer capture. **c**, Comparison of interactions counts from using long or short probes for fragments displayed in panel **a** with Pearson's correlation.

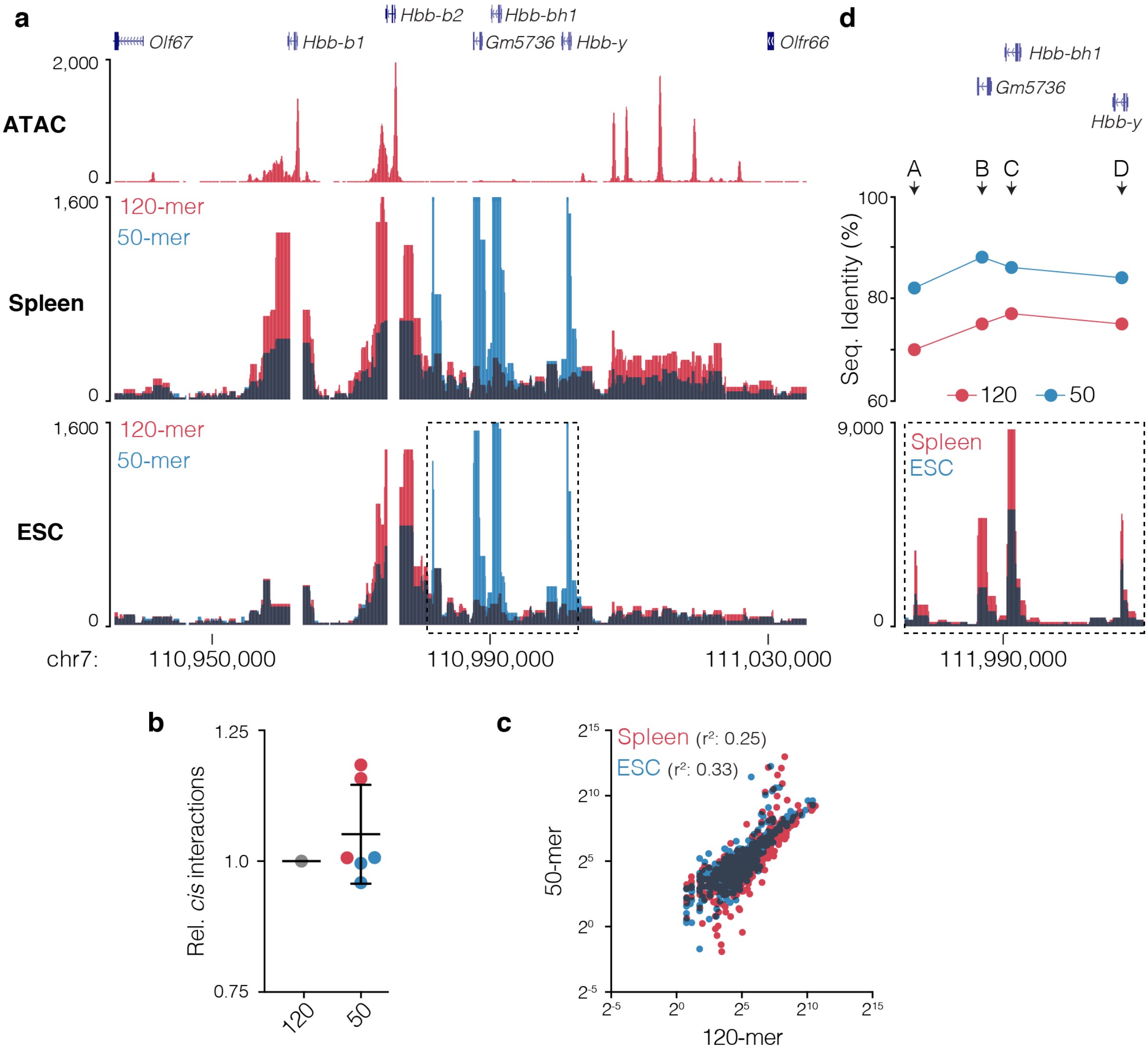


**Supp. Fig. 5 | Capture of β-globin locus with short probes. a,** Overlaid *cis*-normalized 3C interaction profile for *Hbb-b1 and Hbb-b2*, which encode β-globin, from mouse erythroid (n=3) and embryonic stem cells (ESC, n=3) captured with either 120-mer or 50-mer probes. Darkened areas show overlapping signals. **b**, Number of cis reporters relative to 120-mer capture. **c**, Comparison of interactions counts from using long or short probes for fragments displayed in panel **a** with Pearson's correlation. **d,** Clustalω determined sequence identity for 120-mer and 50-mer probes with the four novel peaks of interactions seen with the shorter probes.

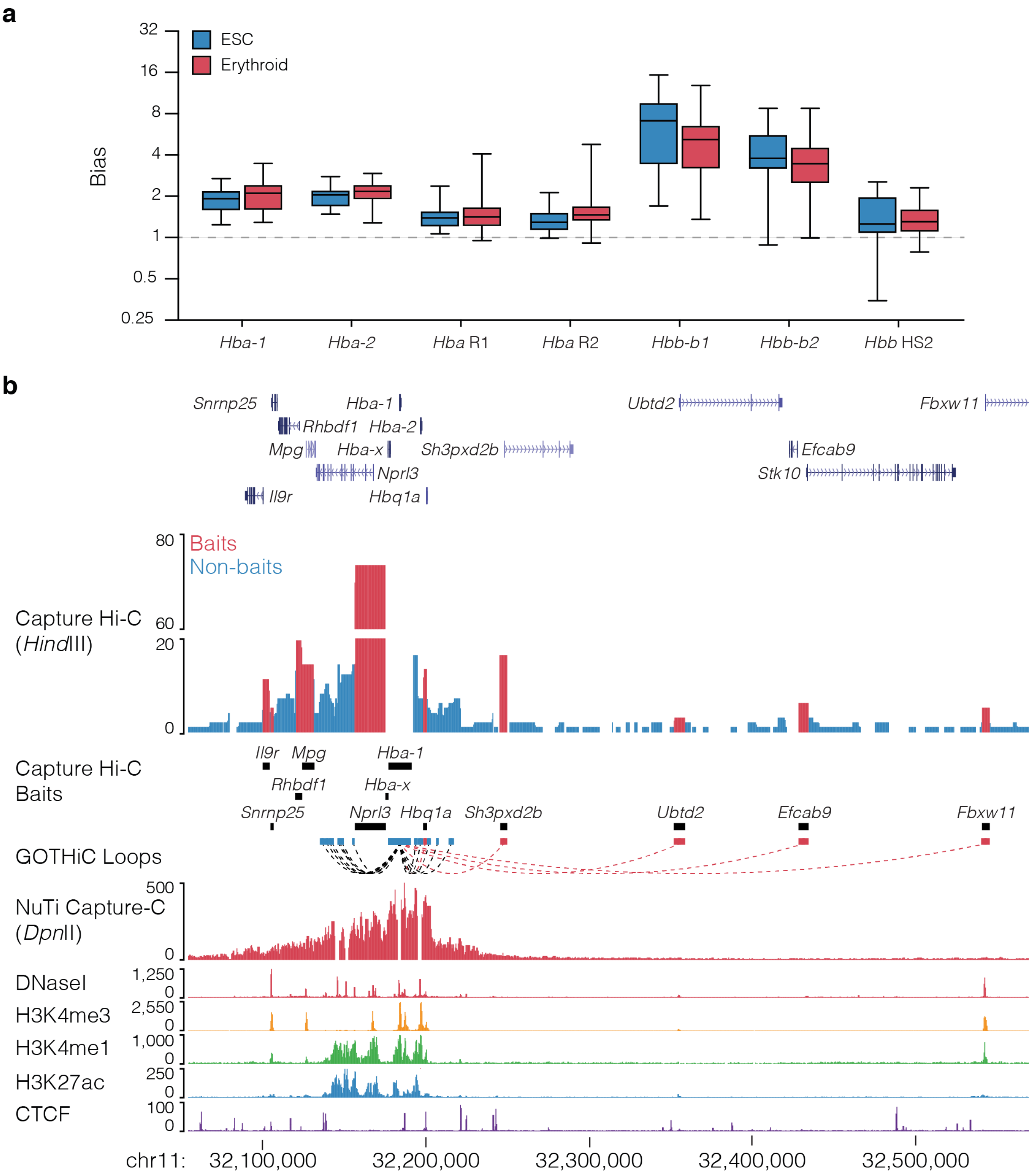


**Supp. Fig. 6 I Co-targeting bias observed in Capture-C and Capture Hi-C. a,** Per viewpoint levels of bias at co-targeted viewpoints around the α-globin locus (*Hba-1* and *Hba-2* promoters, R1 and R2 enhancers) and the β-globin locus (*Hbb-b1* and *Hbb-b2* promoters, HS2 enhancer). Levels of bias varies across viewpoints and co-targeted fragments but not between erythroid and embryonic stem cells (ESC) indicating bias is primarily caused through the identity of the targeted fragment rather than by cell type signal. **b,** Comparison of the 3C interaction profiles for *Hba-1* generated with Capture Hi-C (targeting all promoters) and NuTi Capture-C (targeting specifically *Hba-1/2* and their two main enhancers – excluded from analysis and seen as gaps in the signal). Total interaction counts for CHi-C in erythroid cells are shown (n=2), fragments and reported significant interactions involving co-targeting coloured red. Note that the peaks over reported long-range significant interactions are not present in NuTi Capture-C and occur specifically at co-targeted fragments (and not adjacent fragments). Erythroid tracks show open chromatin (DNaseI), promoters (H3K4me3), active transcription (H3K27ac), enhancers (H3K4me1), and boundaries (CTCF).

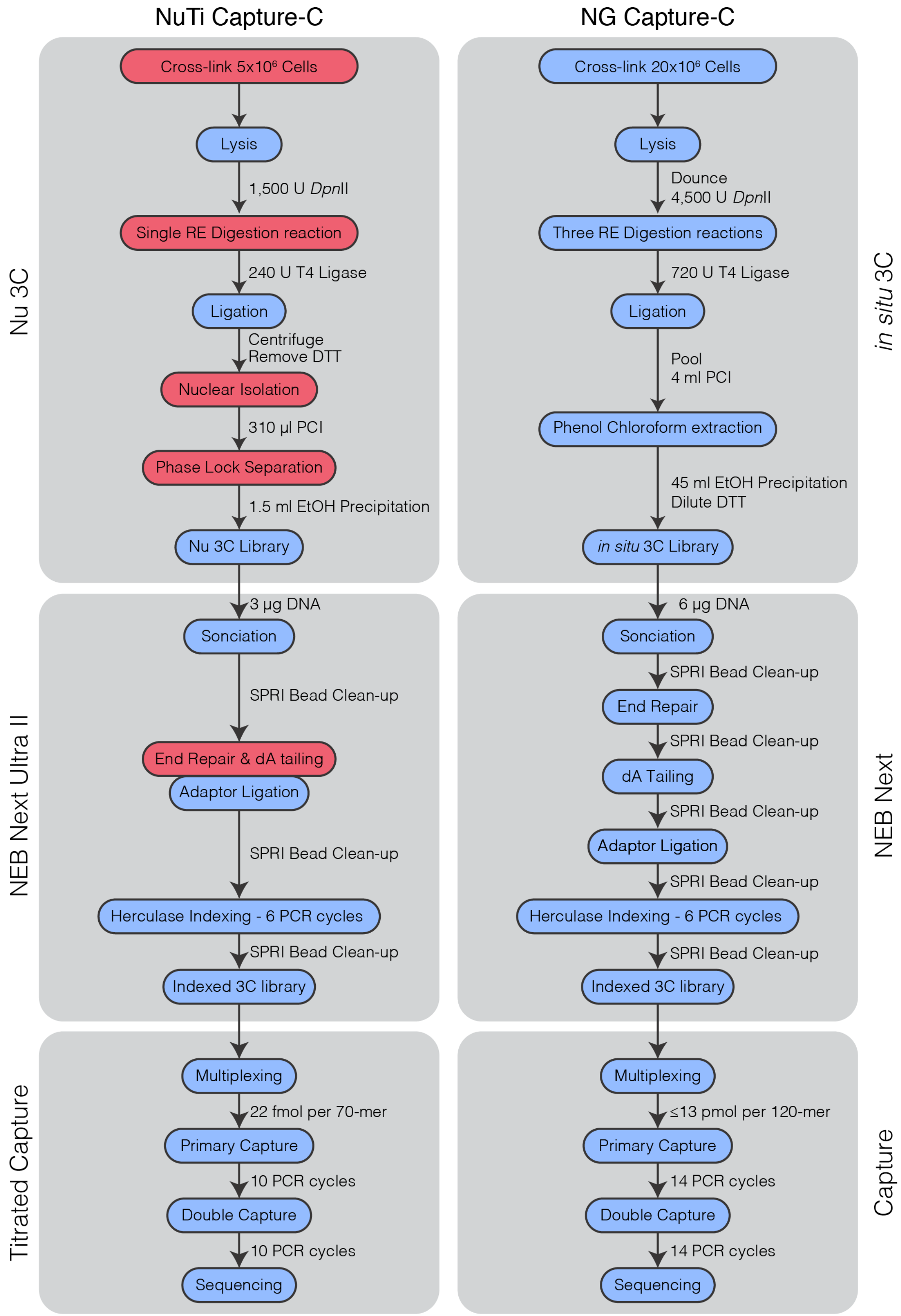


**Supp. Fig. 7 I Capture-C workflows.** Comparison of experimental workflows for Nuclear-Titratred (NuTi) and Next Generation (NG) Capture-C. Main steps are in blue bubbles, with key innovations for NuTi Capture-C highlighted by red bubbles. Differences in reagents and PCR cycles are shown at individual steps. DTT: Dithiothreitol, PCI: Phenol-Chloroform Isoamyl-alcohol, EtOH: ethanol, SPRI: solid phase reversible immobilisation.

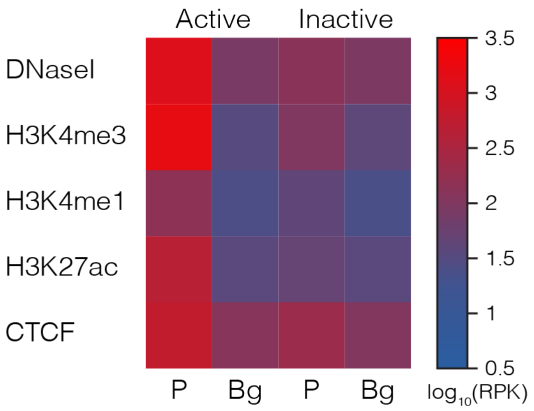


**Supp. Fig. 8 I Chromatin signature of captured promoters.** Average sequence coverage signature of promoter (P) containing fragments (±1kb) classified as active (n=7,014) or inactive (n=181). Chromatin marks from mouse erythroid cells show open chromatin (DNaseI), promoters (H3K4me3), active transcription (H3K27ac), enhancers (H3K4me1), and boundaries (CTCF). Background (Bg) signal was calculated by generating random peaks of the same number and size using BEDtools shuffle. RPK: reads per kilobase.

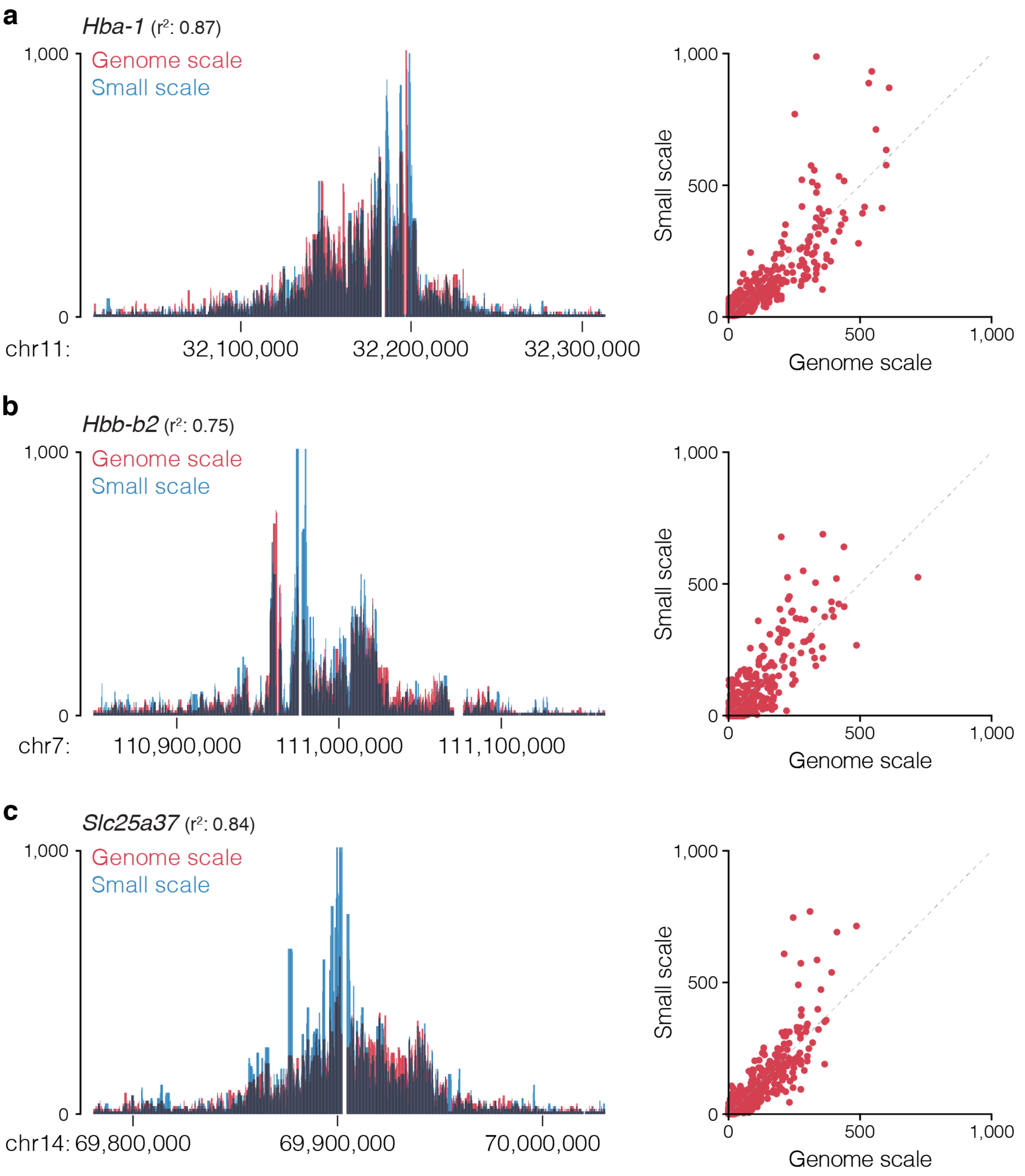


**Supp. Fig. 9 | Genome scale capture closely matches designs with fewer probes.** Overlaid 3C profiles, Pearson correlation values, and per fragment count correlation plots for the *Hba-1* (**a**), *Hbb-b2* (**b**) and *Slc25a37* (**c**) promoters in mouse erythroid cells when targeting <10 (small scale or >7000 (genome scale) viewpoints with NuTi Capture-C. Note overlaid track go dark where signals overlap, seven values >1,000 not shown.

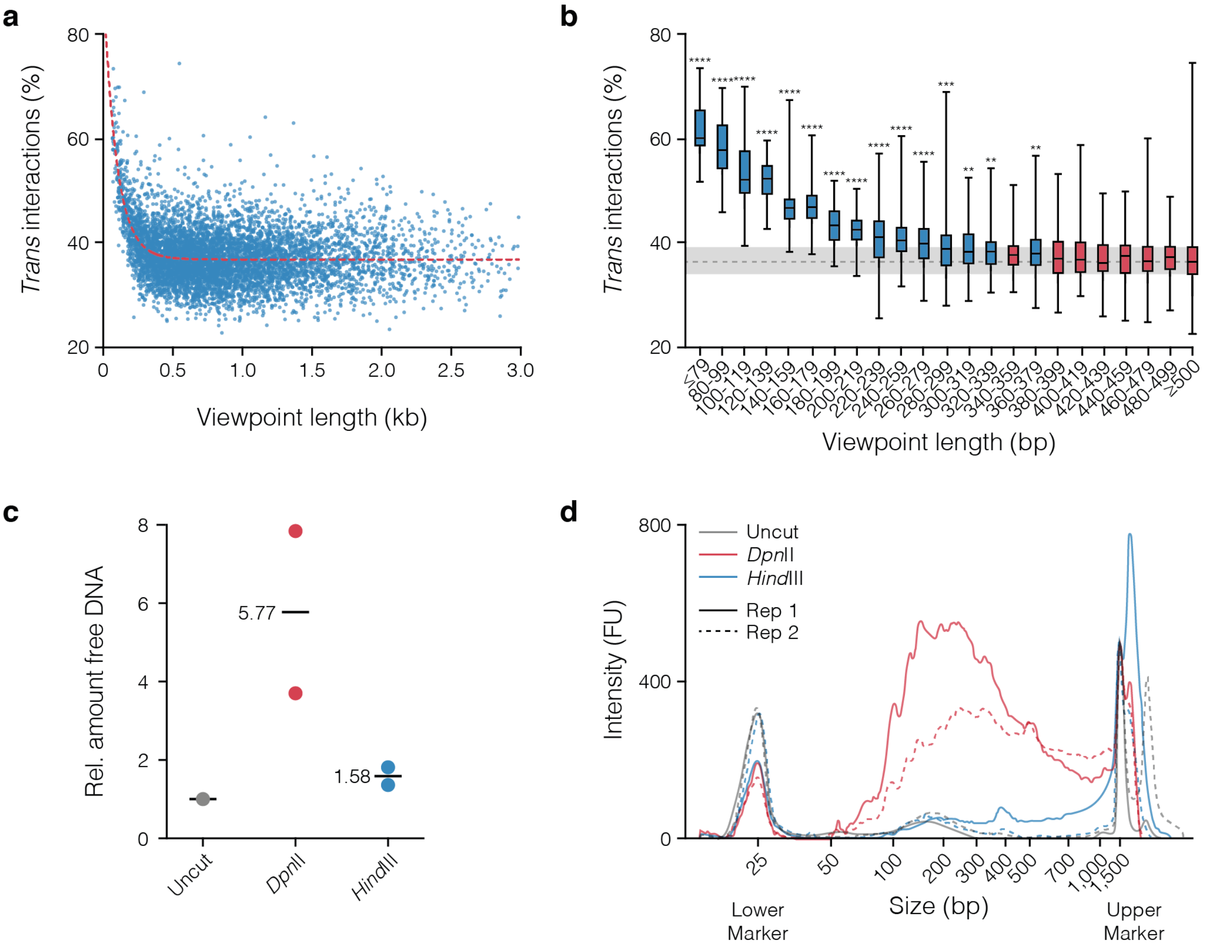


**Supp. Fig. 10 I Short fragments have higher levels of *trans* interactions. a,** Plot of mean percent of *trans* interactions (n=3) for all viewpoints shorter than 3 kb (n=6,659). Red line shows a non-linear fit to the data ($r^2$=0.2150, d.f. 19,972). **b,** Box and whisker plot of viewpoints shorter than 500 bp in 20bp bins (n≥12). A one-way ANOVA was carried out with multiple comparisons for each bin against all viewpoints over 500 bp (n=5,017). Significantly different bins identified by a Dunnett's multiple comparisons test are coloured blue (**p<0.005, ***p<0.0005, ****p<0.0001). Relative amount **(c)** and D1000 tapestation profile **(d)** of DNA recovered from the soluble (non-nuclear) fractions of two 3C samples divided across three tubes each and digested overnight with no enzyme (Uncut), a 4-bp cutter (*Dpn*II), and a 6-bp cutter (*Hind*III). FU: Fluorescent units.

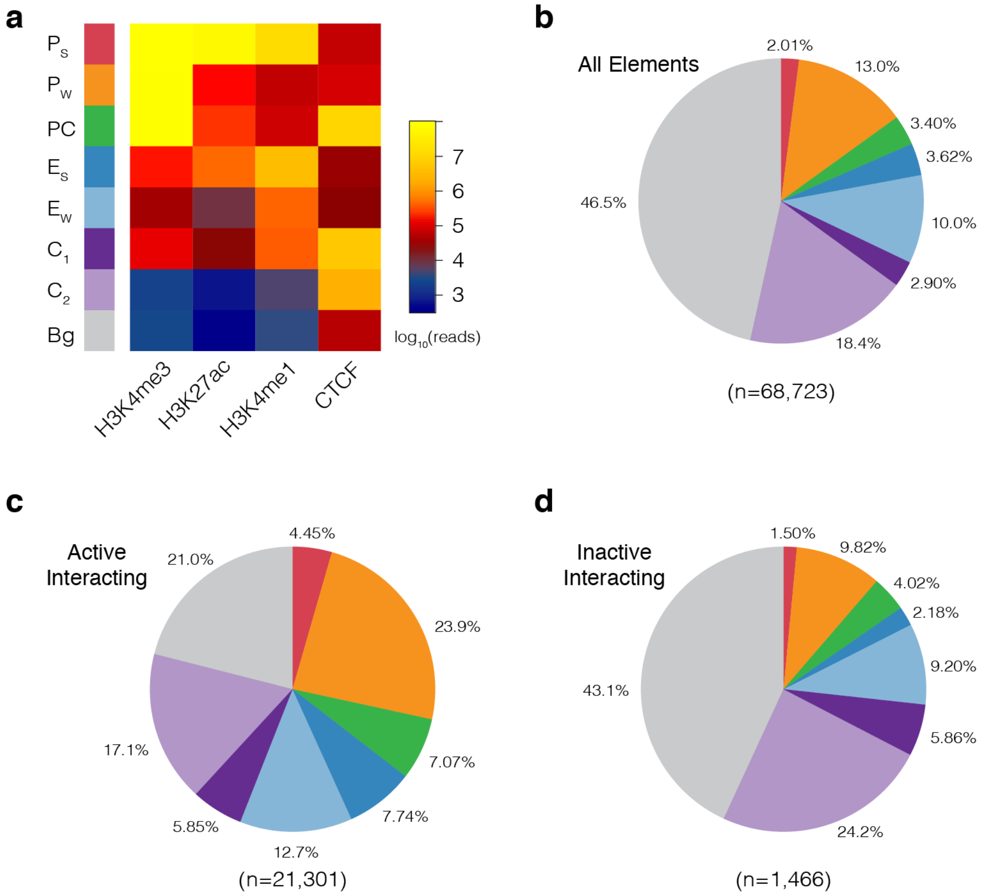


**Supp. Fig. 11 | GenoSTAN annotation of the mouse genome in erythroid cells. a,** Following curation for similar signal profiles, the GenoSTAN Hidden Markov Model identified eight states for 1 kb erythroid open chromatin regions using ChIP-seq for marks associated with promoters (H3K4me3), active transcription (H3K27ac), enhancers (H3K4me1), and boundaries (CTCF). Identified states were named based on average chromatin profile (shown) as: $P_S$: Promoter (Strong H3K27ac), $P_W$: Promoter (Weak H3K27ac), PC: Promoter/CTCF, $E_S$: Enhancer (Strong H3K27ac), $E_W$: Enhancer (Weak H3K27ac), $C_1$: CTCF near promoter/enhancer, $C_2$: CTCF, Bg: Background. Pie charts showing the proportion of unique annotations for all open chromatin regions **(b)**, fragments significantly interacting with active promoters **(c)**, and fragments significantly interacting with inactive promoters **(d).** Pie chart colours and order match the key in panel a.

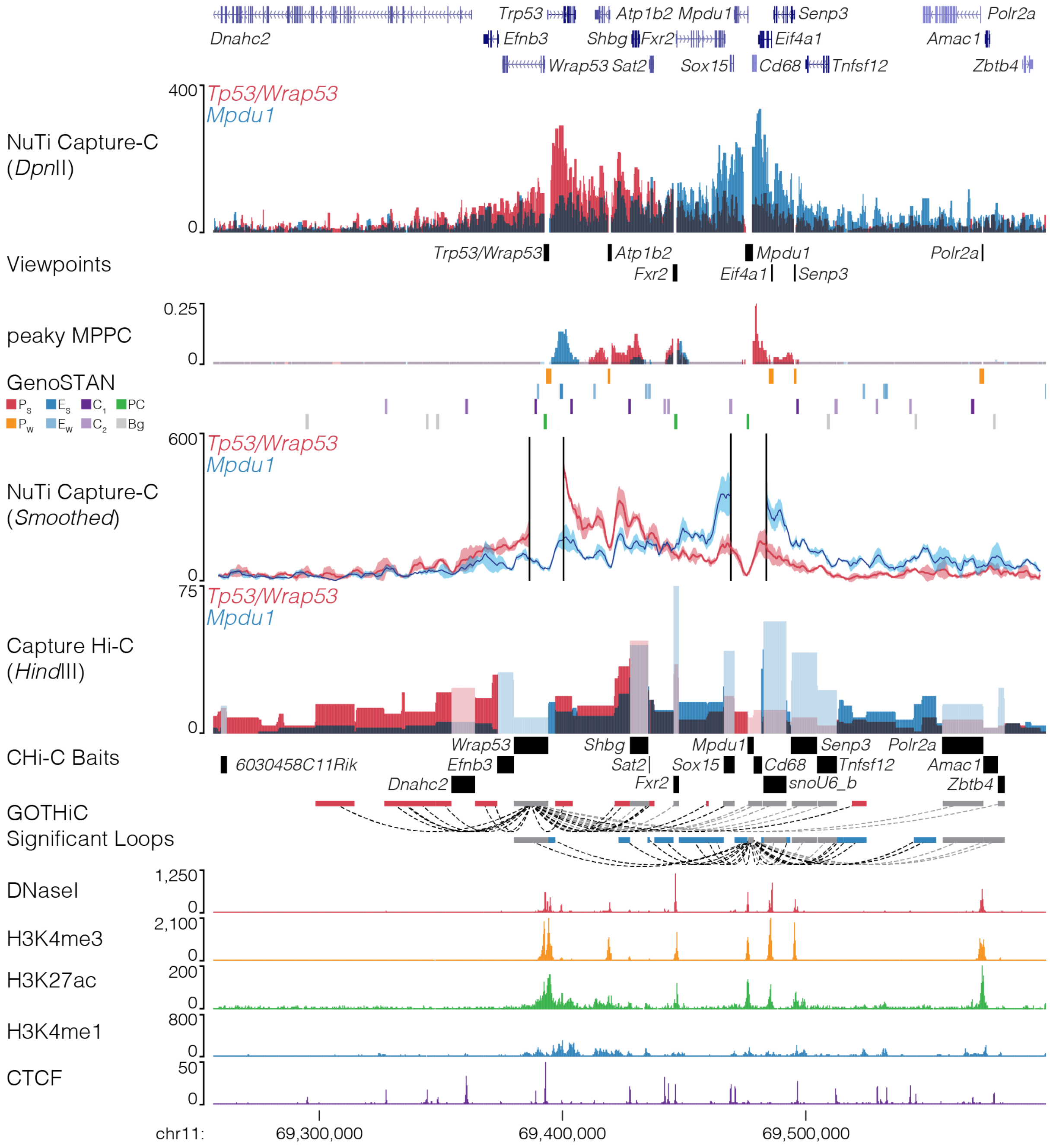


**Supp. Fig. 12 | NuTi Capture-C from the *Tp53, Wrap53* and *Mpdu1* promoters.** Sequence tracks showing the difference between high-resolution 3C (*Dpn*II, NuTi Capture-C) and low-resolution 3C (*Hind*III, Capture Hi-C) at gene promoters in the same regulatory domain in erythroid cells (mm9, chr11:69,256,536-69,598,480). Tracks in order: UCSC gene annotation, *cis*-normalized mean interactions per *Dpn*II fragment using NuTi Capture-C (n=3), NuTi Capture-C viewpoints, peaky Marginal Posterior Probability of Contact (MPPC) scores with fragments with MPPC ≥0.01 darker, GenoSTAN open chromatin classification, windowed mean interactions using NuTi Capture-C, total supporting reads per *Hind*III fragment with CHi-C (n=2; co-targeted fragments are lighter in colour), CHi-C bait fragments, loops between reported significantly interacting fragments (co-targeting loops are coloured grey), erythroid tracks for open chromatin (DNaseI), promoters (H3K4me3), active transcription (H3K27ac), enhancers (H3K4me1), and boundaries (CTCF). Note overlapping blue and red signals appear darker in colour (NuTi Capture-C, peaky MPPC, CHi-C).

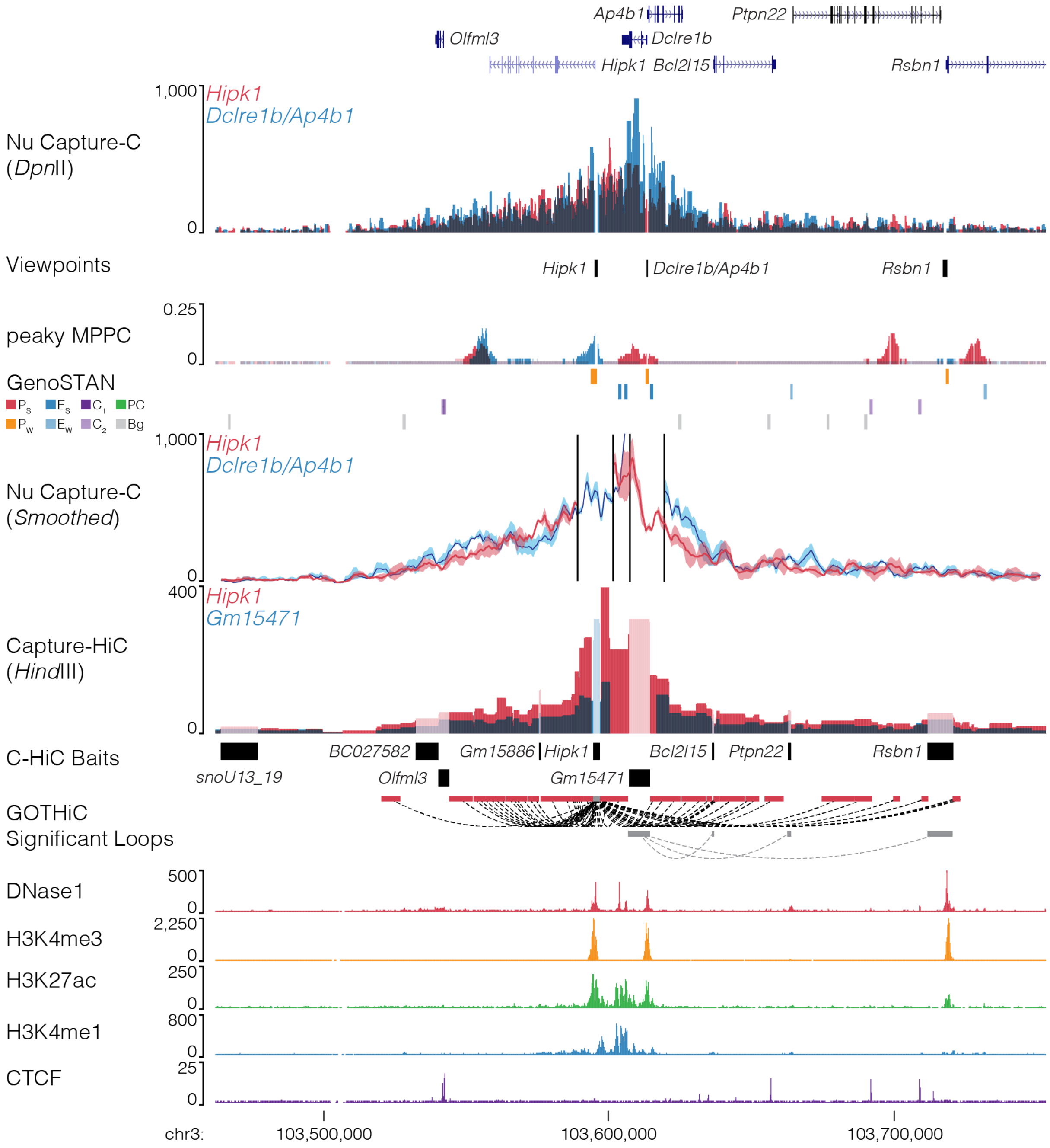


**Fig. 13 l NuTi Capture-C from the *Hipk1, Dclre1b* and *Ap4b1* promoters.** Sequence tracks showing the difference between high-resolution 3C (*Dpn*II, NuTi Capture-C) and low-resolution 3C (*Hind*III, Capture Hi-C) at gene promoters in the same regulatory domain in erythroid cells (mm9, chr3:103,462,115-103,753,122). Tracks in order: UCSC gene annotation, *cis*-normalized mean interactions per *Dpn*II fragment using NuTi Capture-C (n=3), NuTi Capture-C viewpoints, peaky Marginal Posterior Probability of Contact (MPPC) scores with fragments with MPPC ≥0.01 darker, GenoSTAN open chromatin classification, windowed mean interactions using NuTi Capture-C, total supporting reads per *Hind*III fragment with CHi-C (n=2; co-targeted fragments are lighter in colour), CHi-C bait fragments, loops between reported significantly interacting fragments (co-targeting loops are coloured grey), erythroid tracks for open chromatin (DNaseI), promoters (H3K4me3), active transcription (H3K27ac), enhancers (H3K4me1), and boundaries (CTCF). Note overlapping blue and red signals appear darker in colour (NuTi Capture-C, peaky MPPC, CHi-C).

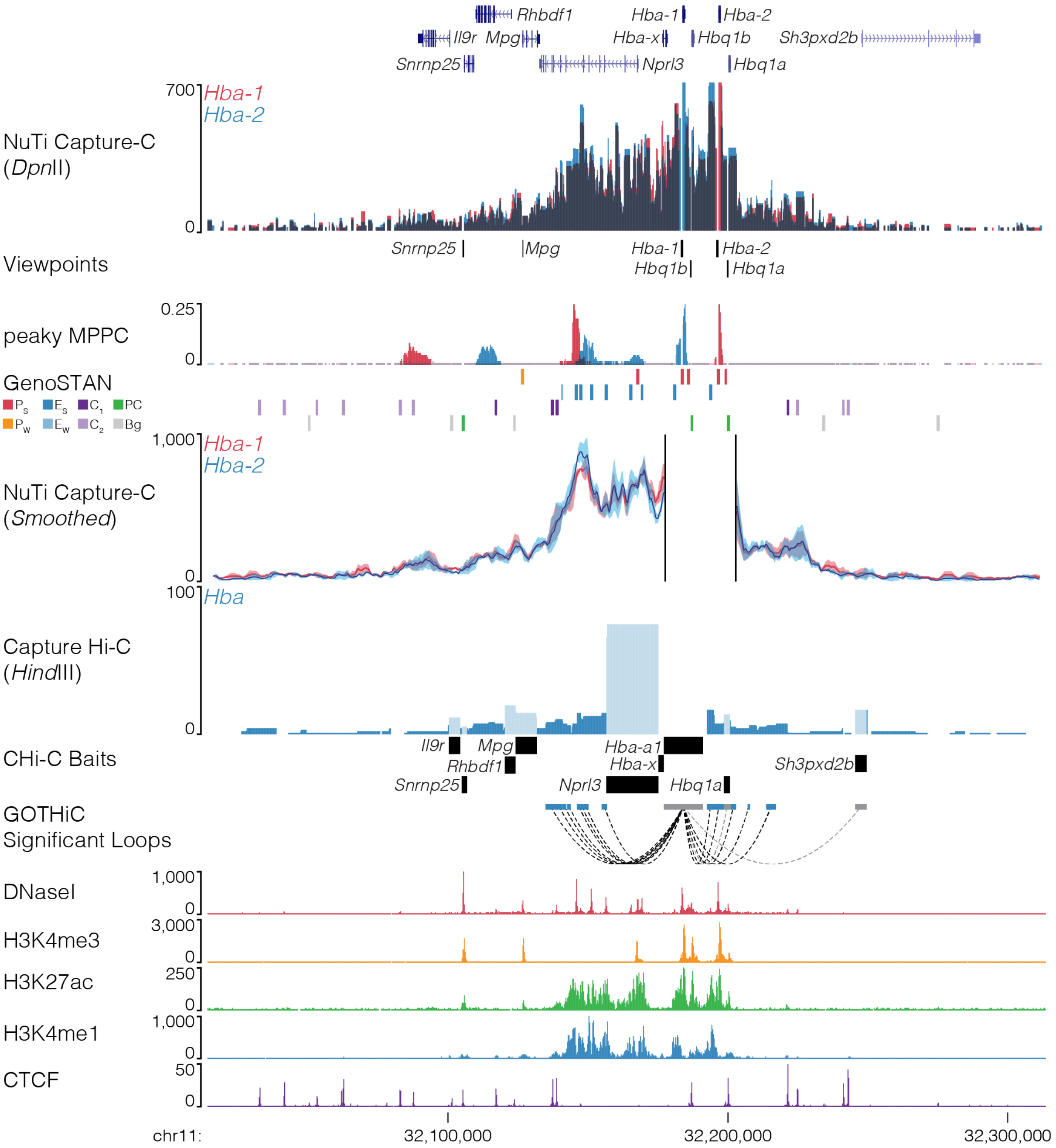


**Fig. 14 | NuTi Capture-C from the *Hba-1* and *Hba-2* promoters.** Sequence tracks showing the difference between high-resolution 3C (*Dpn*II, NuTi Capture-C) and low-resolution 3C (*Hind*III, Capture Hi-C) at gene promoters in the same regulatory domain in erythroid cells (mm9, chr3:103,462,115-103,753,122). Tracks in order: UCSC gene annotation, *cis*-normalized mean interactions per *Dpn*II fragment using NuTi Capture-C (n=3), NuTi Capture-C viewpoints, peaky Marginal Posterior Probability of Contact (MPPC) scores with fragments with MPPC ≥0.01 darker, GenoSTAN open chromatin classification, windowed mean interactions using NuTi Capture-C, total supporting reads per *Hind*III fragment with CHi-C (n=2; co-targeted fragments are lighter in colour), CHi-C bait fragments, loops between reported significantly interacting fragments (co-targeting loops are coloured grey), erythroid tracks for open chromatin (DNaseI), promoters (H3K4me3), active transcription (H3K27ac), enhancers (H3K4me1), and boundaries (CTCF). Note overlapping blue and red signals appear darker in colour (NuTi Capture-C, peaky MPPC, CHi-C).

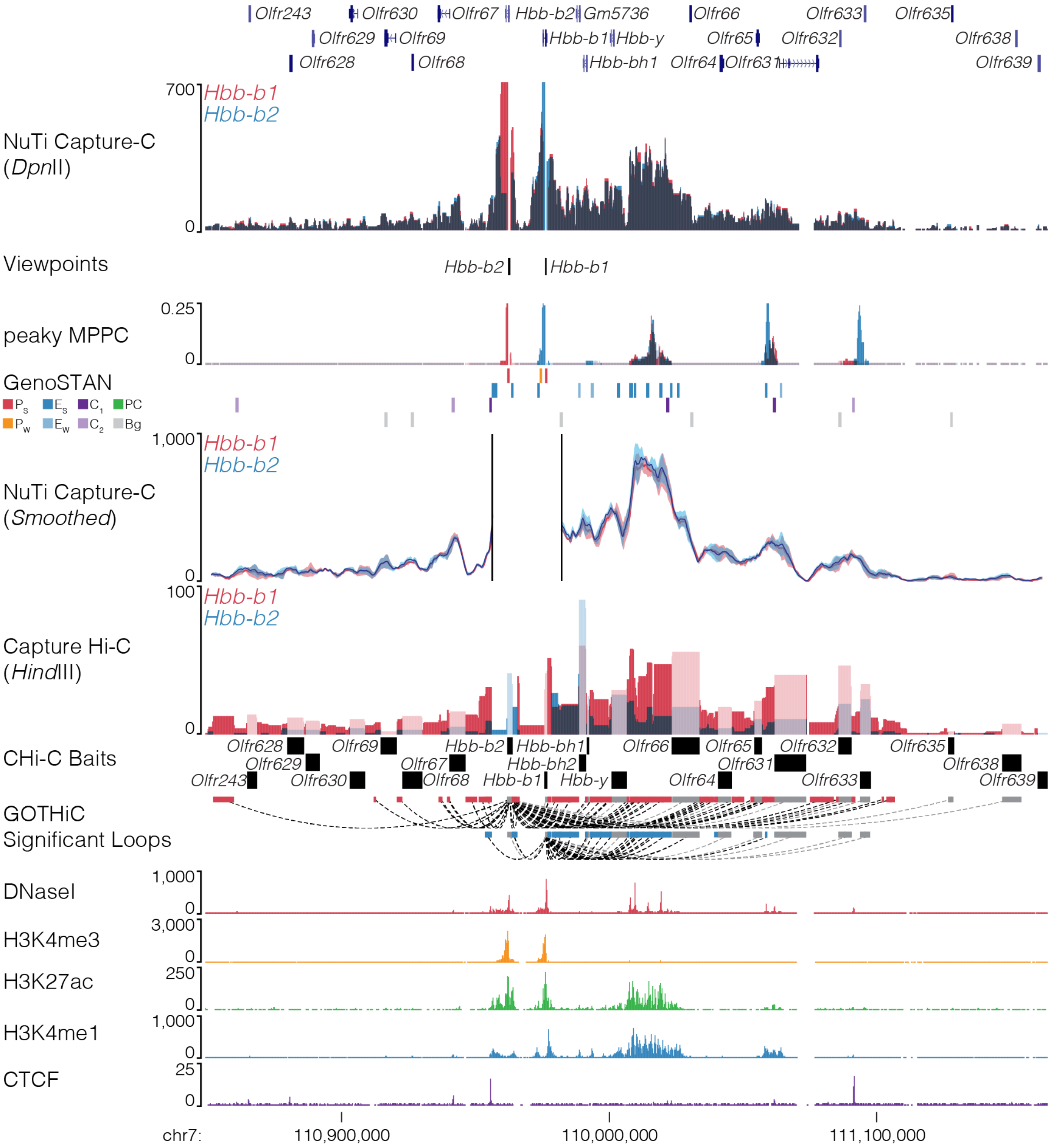


**Fig. 15 | NuTi Capture-C from the *Hbb-b1* and *Hbb-b2* promoters.** Sequence tracks showing the difference between high-resolution 3C (*Dpn*II, NuTi Capture-C) and low-resolution 3C (*Hind*III, Capture Hi-C) at gene promoters in the same regulatory domain in erythroid cells (mm9, chr7:110,848,909-111,163,908). Tracks in order: UCSC gene annotation, *cis*-normalized mean interactions per *Dpn*II fragment using NuTi Capture-C (n=3), NuTi Capture-C viewpoints, peaky Marginal Posterior Probability of Contact (MPPC) scores with fragments with MPPC ≥0.01 darker, GenoSTAN open chromatin classification, windowed mean interactions using NuTi Capture-C, total supporting reads per *Hind*III fragment with CHi-C (n=2; co-targeted fragments are lighter in colour), CHi-C bait fragments, loops between reported significantly interacting fragments (co-targeting loops are coloured grey), erythroid tracks for open chromatin (DNaseI), promoters (H3K4me3), active transcription (H3K27ac), enhancers (H3K4me1), and boundaries (CTCF). Note overlapping blue and red signals appear darker in colour (NuTi Capture-C, peaky MPPC, CHi-C).

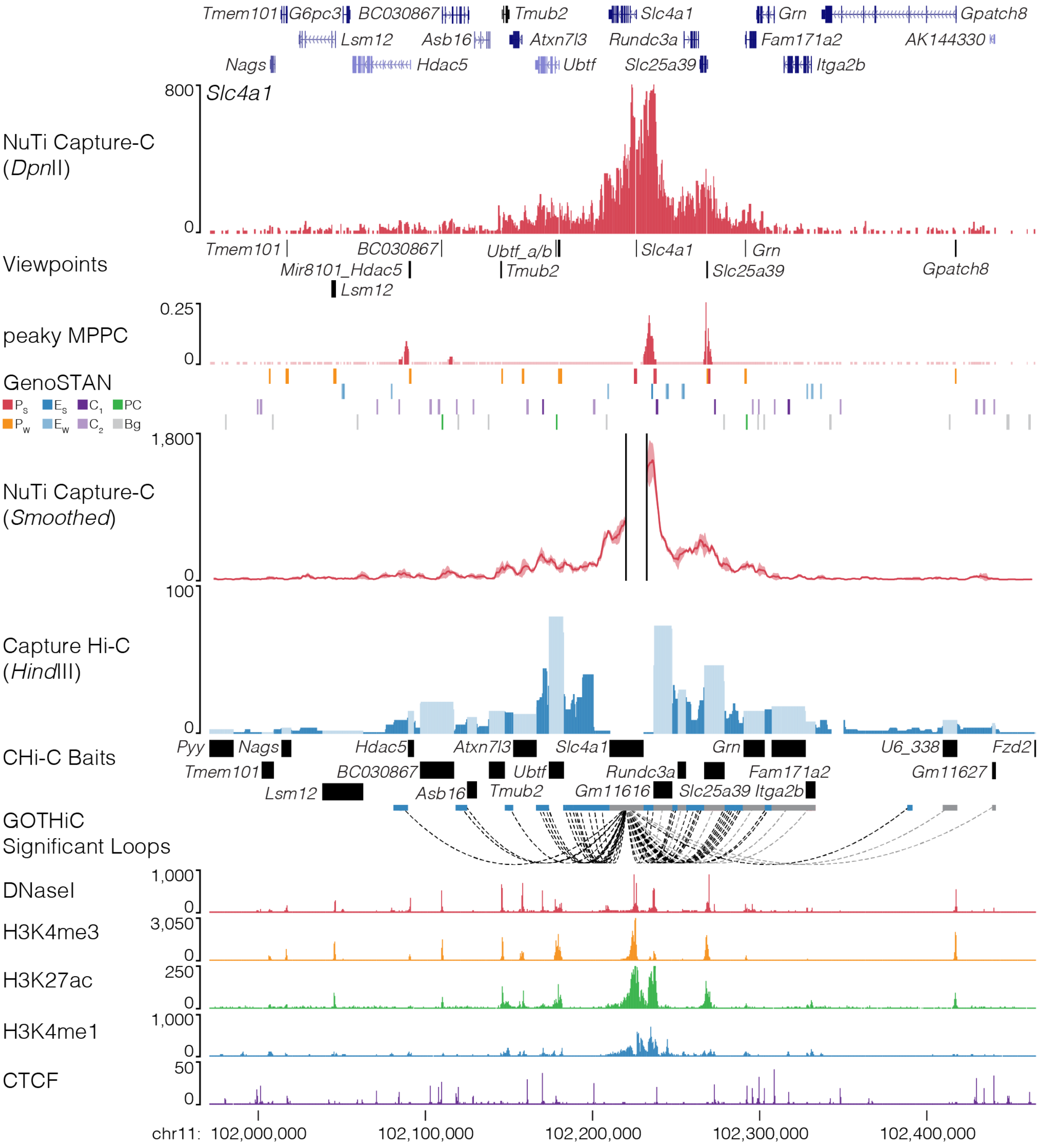


**Fig. 16 | NuTi Capture-C from the *Slc4a1* promoter.** Sequence tracks showing the difference between high-resolution 3C (*Dpn*II, NuTi Capture-C) and low-resolution 3C (*Hind*III, Capture Hi-C) at calling interacting fragments (mm9, chr11:101,971,435-102,465,294) in erythroid cells. Tracks in order: UCSC gene annotation, *cis*-normalized mean interactions per *Dpn*II fragment using NuTi Capture-C (n=3), NuTi Capture-C viewpoints, peaky Marginal Posterior Probability of Contact (MPPC) scores with fragments with MPPC ≥0.01 darker, GenoSTAN open chromatin classification, windowed mean interactions using NuTi Capture-C, total supporting reads per *Hind*III fragment with CHi-C (n=2; co-targeted fragments are lighter in colour), CHi-C bait fragments, loops between reported significantly interacting fragments (co-targeting loops are coloured grey), erythroid tracks for open chromatin (DNaseI), promoters (H3K4me3), active transcription (H3K27ac), enhancers (H3K4me1), and boundaries (CTCF). Note overlapping MPPC signals appear darker in colour.

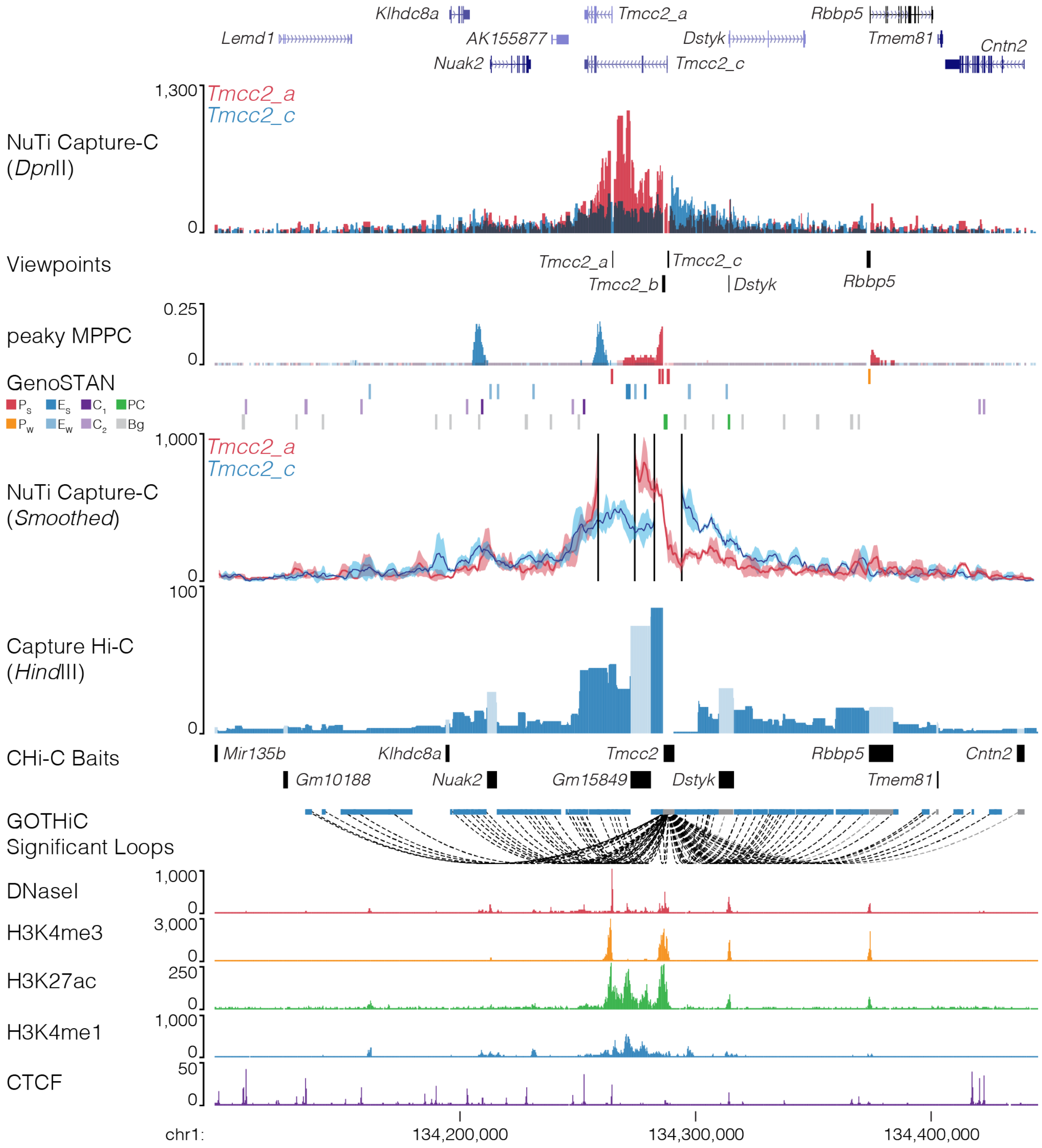


**Fig. 17 | NuTi Capture-C from alternative *Tmcc2* promoters.** Sequence tracks showing the difference between high-resolution 3C (*Dpn*II, NuTi Capture-C) and low-resolution 3C (*Hind*III, Capture Hi-C) from alternative *Tmmc2* promoters (mm9, chr1:134,095,540-134,445,539) in erythroid cells. Tracks in order: UCSC gene annotation, *cis*-normalized mean interactions per *Dpn*II fragment using NuTi Capture-C (n=3), NuTi Capture-C viewpoints, peaky Marginal Posterior Probability of Contact (MPPC) scores with fragments with MPPC ≥0.01 darker, GenoSTAN open chromatin classification, windowed mean interactions using NuTi Capture-C, total supporting reads per *Hind*III fragment with CHi-C (n=2; co-targeted fragments are lighter in colour), CHi-C bait fragments, loops between reported significantly interacting fragments (co-targeting loops are coloured grey), erythroid tracks for open chromatin (DNaseI), promoters (H3K4me3), active transcription (H3K27ac), enhancers (H3K4me1), and boundaries (CTCF). Note overlapping blue and red signals appear darker in colour (NuTi Capture-C, peaky MPPC, CHi-C).

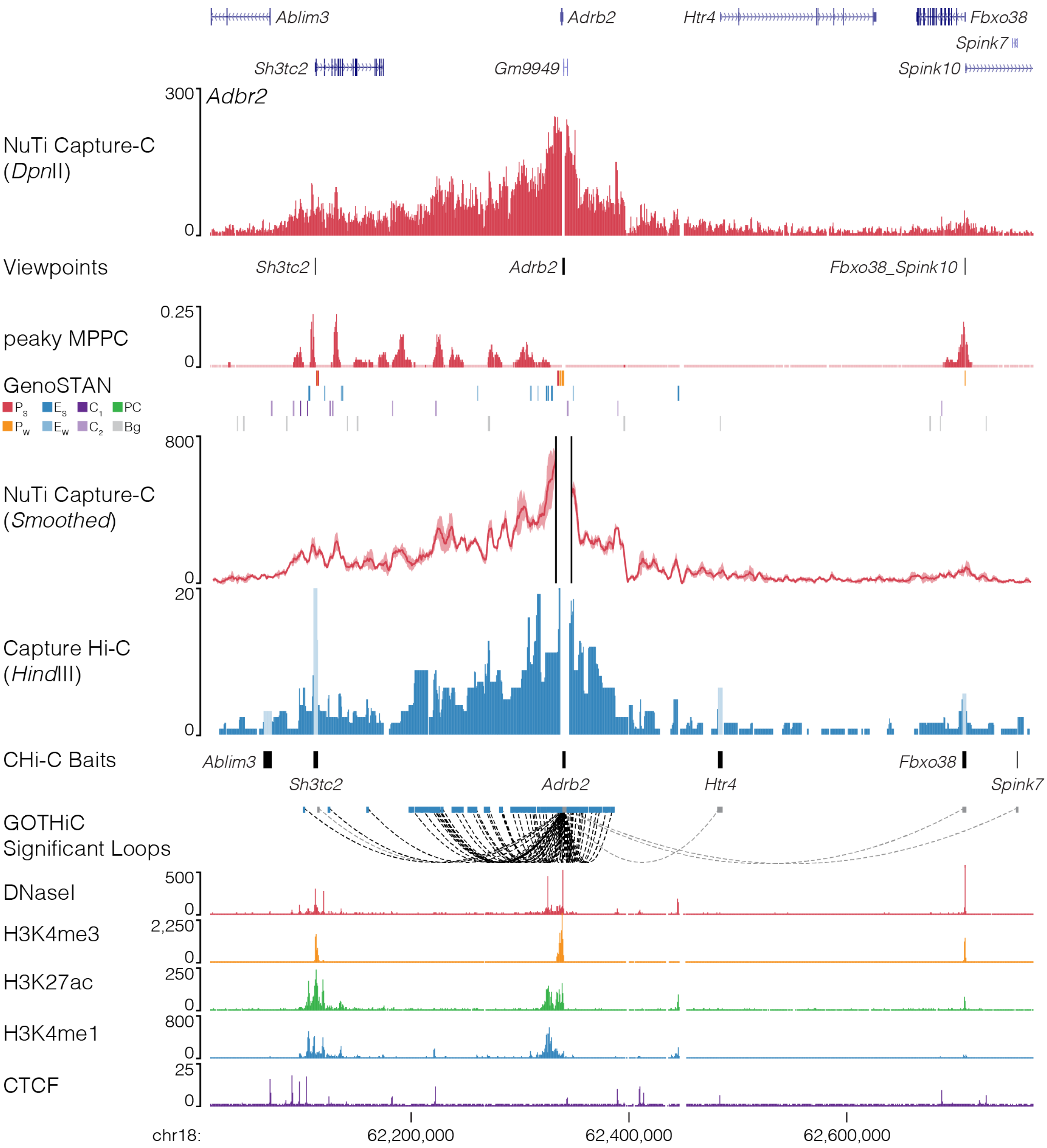


**Fig. 18 | NuTi Capture-C from the *Adrb2* promoter.** Sequence tracks showing the difference between high-resolution 3C (*Dpn*II, NuTi Capture-C) and low-resolution 3C (*Hind*III, Capture Hi-C) at calling interacting fragments (mm9, chr18:62,016,212-62,771,180) in erythroid cells. Tracks in order: UCSC gene annotation, *cis*-normalized mean interactions per *Dpn*II fragment using NuTi Capture-C (n=3), NuTi Capture-C viewpoints, peaky Marginal Posterior Probability of Contact (MPPC) scores with fragments with MPPC ≥0.01 darker, GenoSTAN open chromatin classification, windowed mean interactions using NuTi Capture-C, total supporting reads per *Hind*III fragment with CHi-C (n=2; co-targeted fragments are lighter in colour), CHi-C bait fragments, loops between reported significantly interacting fragments (co-targeting loops are coloured grey), erythroid tracks for open chromatin (DNaseI), promoters (H3K4me3), active transcription (H3K27ac), enhancers (H3K4me1), and boundaries (CTCF). Note overlapping MPPC signals appear darker in colour.

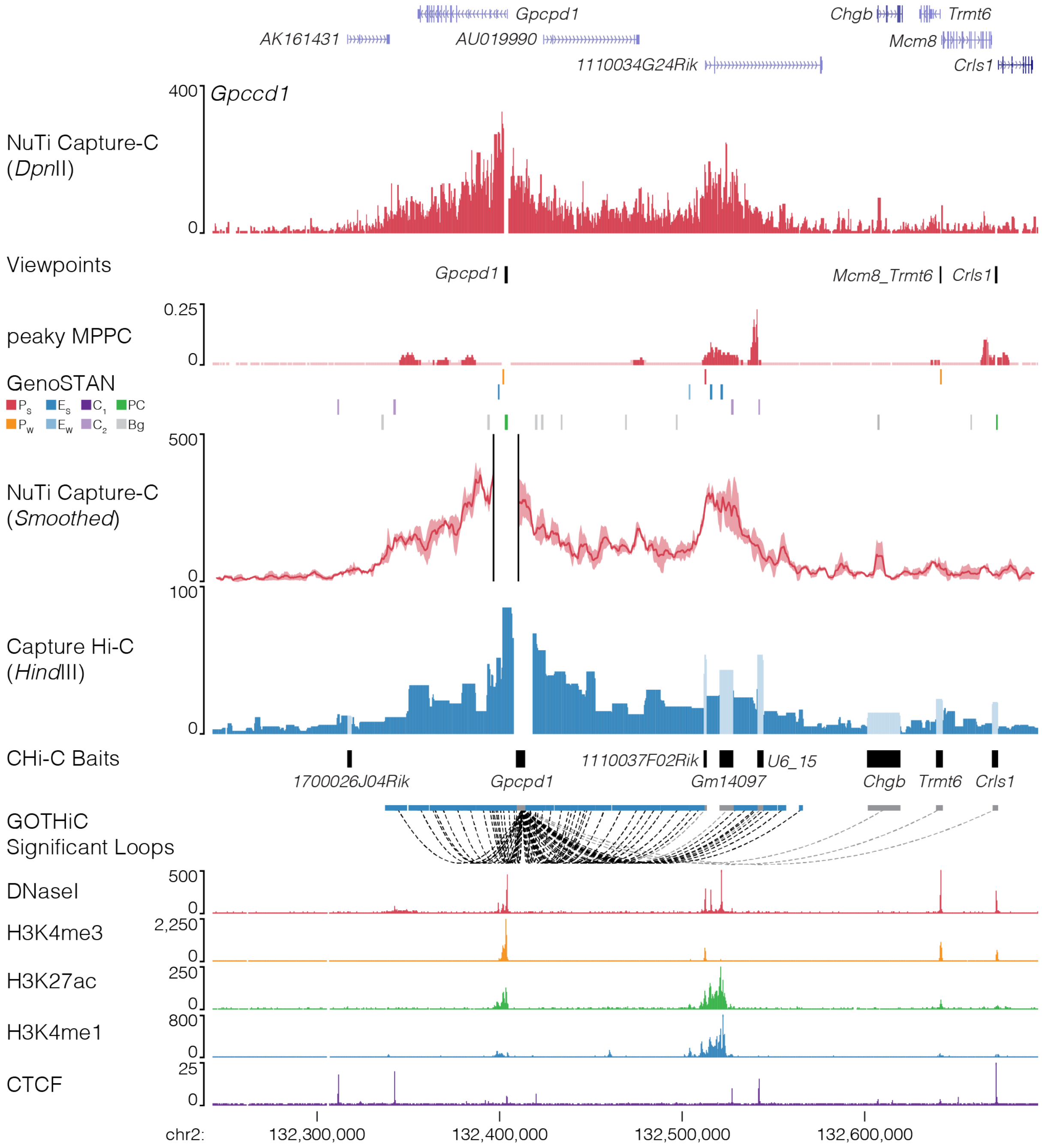


**Fig. 19 l NuTi Capture-C from the *Gpcpd1* promoters.** Sequence tracks showing the difference between high-resolution 3C (*Dpn*II, NuTi Capture-C) and low-resolution 3C (*Hind*III, Capture Hi-C) at calling interacting fragments (mm9, chr2:132,242,416-132,695,297) in erythroid cells. Tracks in order: UCSC gene annotation, *cis*-normalized mean interactions per *Dpn*II fragment using NuTi Capture-C (n=3), NuTi Capture-C viewpoints, peaky Marginal Posterior Probability of Contact (MPPC) scores with fragments with MPPC $\geq$0.01 darker, GenoSTAN open chromatin classification, windowed mean interactions using NuTi Capture-C, total supporting reads per *Hind*III fragment with CHi-C (n=2; co-targeted fragments are lighter in colour), CHi-C bait fragments, loops between reported significantly interacting fragments (co-targeting loops are coloured grey), erythroid tracks for open chromatin (DNaseI), promoters (H3K4me3), active transcription (H3K27ac), enhancers (H3K4me1), and boundaries (CTCF). Note overlapping MPPC signals appear darker in colour.

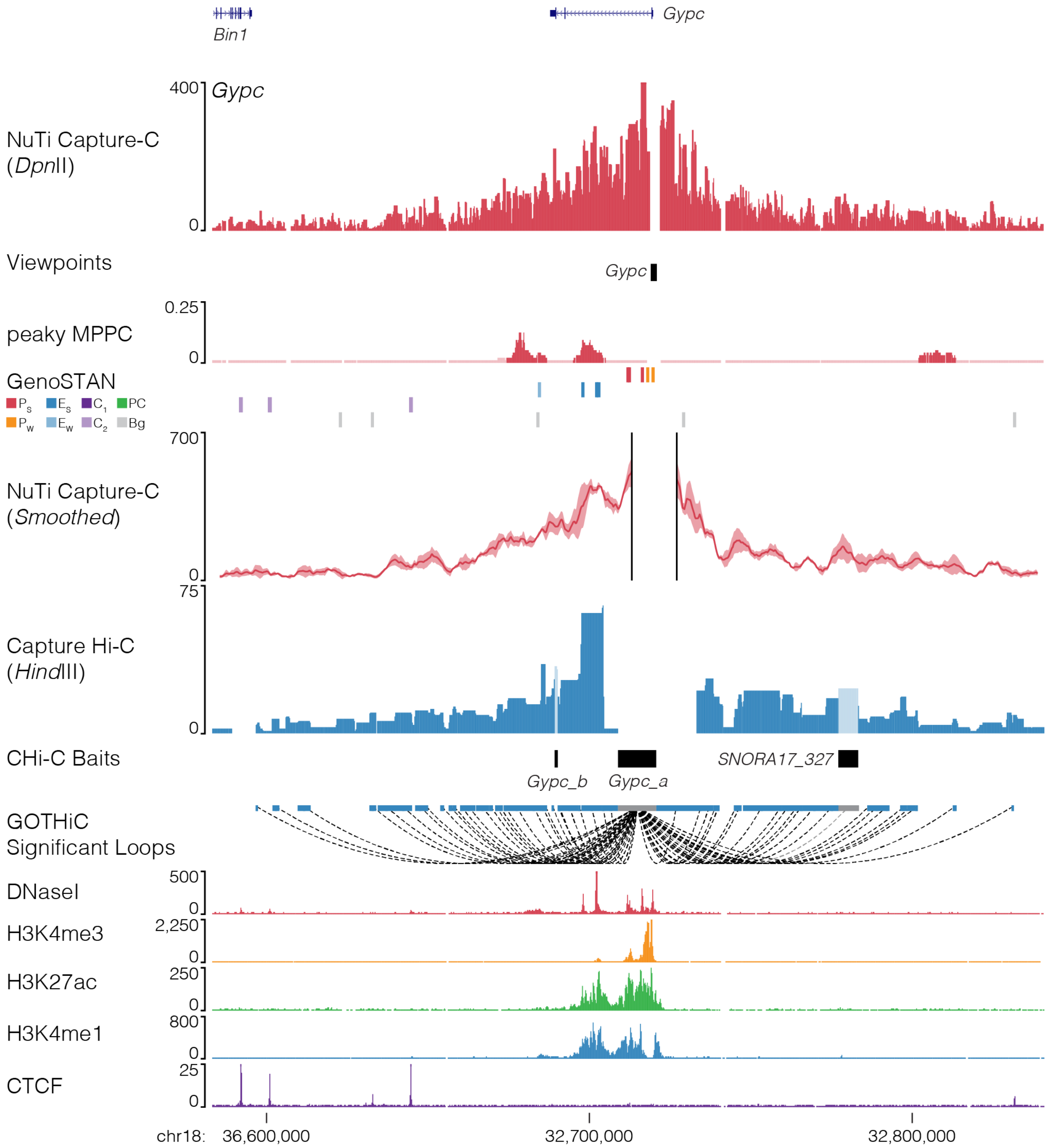


**Fig. 20 | NuTi Capture-C from the *Gypc* promoter.** Sequence tracks showing the difference between high-resolution 3C (*Dpn*II, NuTi Capture-C) and low-resolution 3C (*Hind*III, Capture Hi-C) at calling interacting fragments (mm9, chr18:32,583,205-32,841,048) in erythroid cells. Tracks in order: UCSC gene annotation, *cis*-normalized mean interactions per *Dpn*II fragment using NuTi Capture-C (n=3), NuTi Capture-C viewpoints, peaky Marginal Posterior Probability of Contact (MPPC) scores with fragments with MPPC ≥0.01 darker, GenoSTAN open chromatin classification, windowed mean interactions using NuTi Capture-C, total supporting reads per *Hind*III fragment with CHi-C (n=2; co-targeted fragments are lighter in colour), CHi-C bait fragments, loops between reported significantly interacting fragments (co-targeting loops are coloured grey), erythroid tracks for open chromatin (DNaseI), promoters (H3K4me3), active transcription (H3K27ac), enhancers (H3K4me1), and boundaries (CTCF). Note overlapping MPPC signals appear darker in colour.

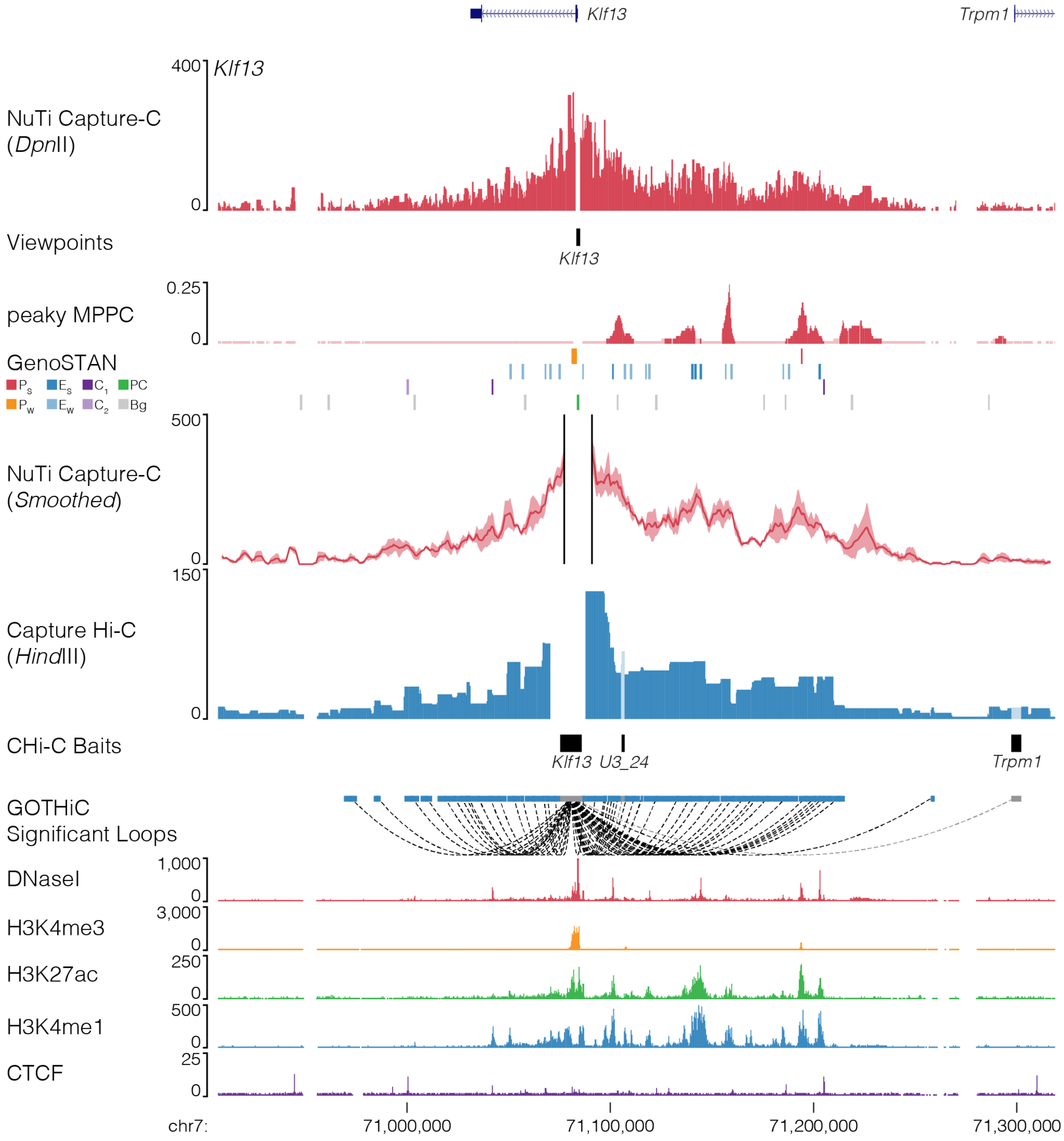


**Fig. 21 | NuTi Capture-C from the *Klf13* promoters.** Sequence tracks showing the difference between high-resolution 3C (*Dpn*II, NuTi Capture-C) and low-resolution 3C (*Hind*III, Capture Hi-C) at calling interacting fragments (mm9, chr7:70,906,844-71,318,843) in erythroid cells. Tracks in order: UCSC gene annotation, *cis*-normalized mean interactions per *Dpn*II fragment using NuTi Capture-C (n=3), NuTi Capture-C viewpoints, peaky Marginal Posterior Probability of Contact (MPPC) scores with fragments with MPPC ≥0.01 darker, GenoSTAN open chromatin classification, windowed mean interactions using NuTi Capture-C, total supporting reads per *Hind*III fragment with CHi-C (n=2; co-targeted fragments are lighter in colour), CHi-C bait fragments, loops between reported significantly interacting fragments (co-targeting loops are coloured grey), erythroid tracks for open chromatin (DNaseI), promoters (H3K4me3), active transcription (H3K27ac), enhancers (H3K4me1), and boundaries (CTCF). Note overlapping MPPC signals appear darker in colour.

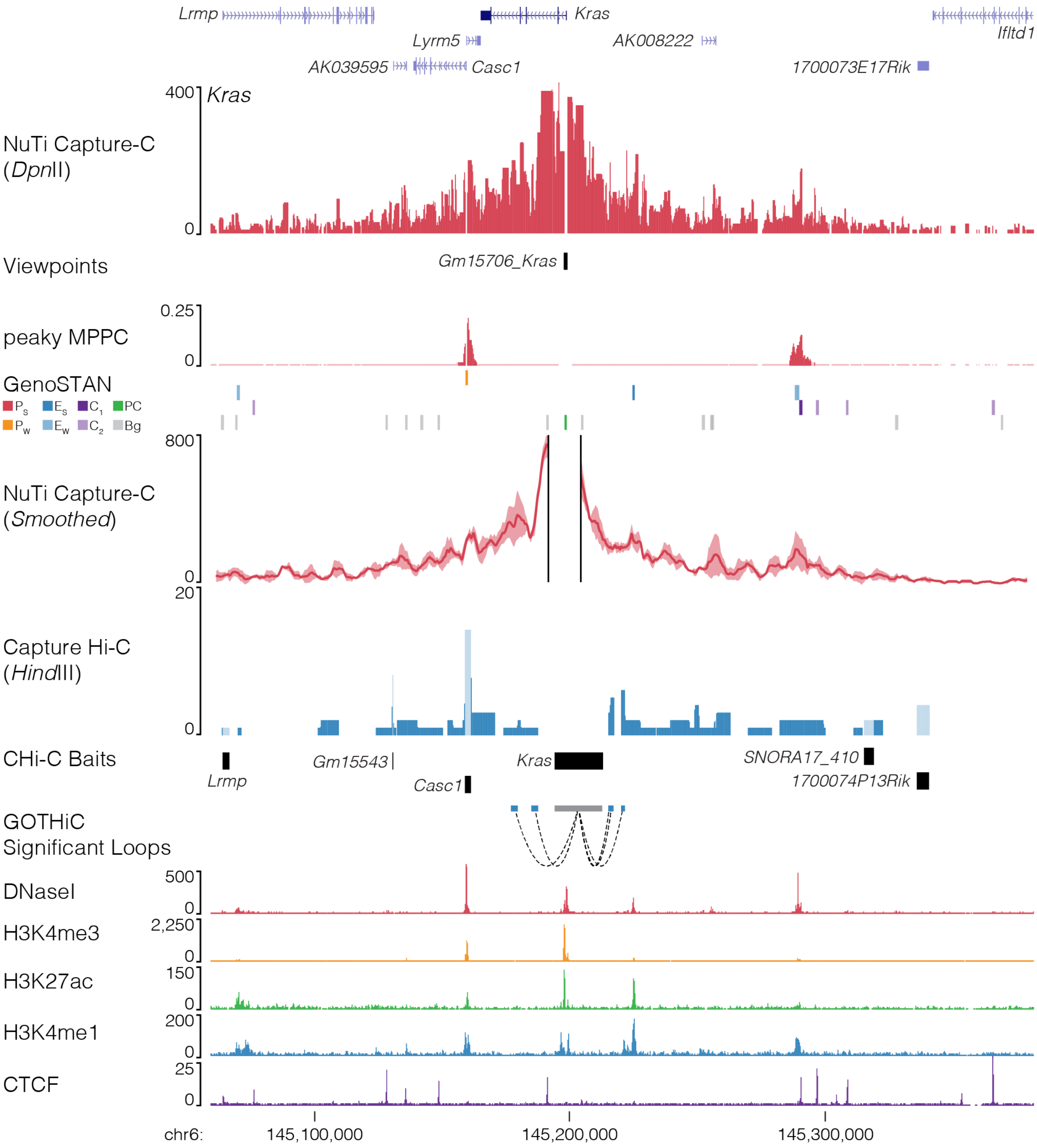


**Fig. 22 | NuTi Capture-C from the *Kras* promoters** Sequence tracks showing the difference between high-resolution 3C (*Dpn*II, NuTi Capture-C) and low-resolution 3C (*Hind*III, Capture Hi-C) at calling interacting fragments (mm9, chr6:145,059,451-145,381,678) in erythroid cells. Tracks in order: UCSC gene annotation, *cis*-normalized mean interactions per *Dpn*II fragment using NuTi Capture-C (n=3), NuTi Capture-C viewpoints, peaky Marginal Posterior Probability of Contact (MPPC) scores with fragments with MPPC ≥0.01 darker, GenoSTAN open chromatin classification, windowed mean interactions using NuTi Capture-C, total supporting reads per *Hind*III fragment with CHi-C (n=2; co-targeted fragments are lighter in colour), CHi-C bait fragments, loops between reported significantly interacting fragments (co-targeting loops are coloured grey), erythroid tracks for open chromatin (DNaseI), promoters (H3K4me3), active transcription (H3K27ac), enhancers (H3K4me1), and boundaries (CTCF). Note overlapping MPPC signals appear darker in colour.

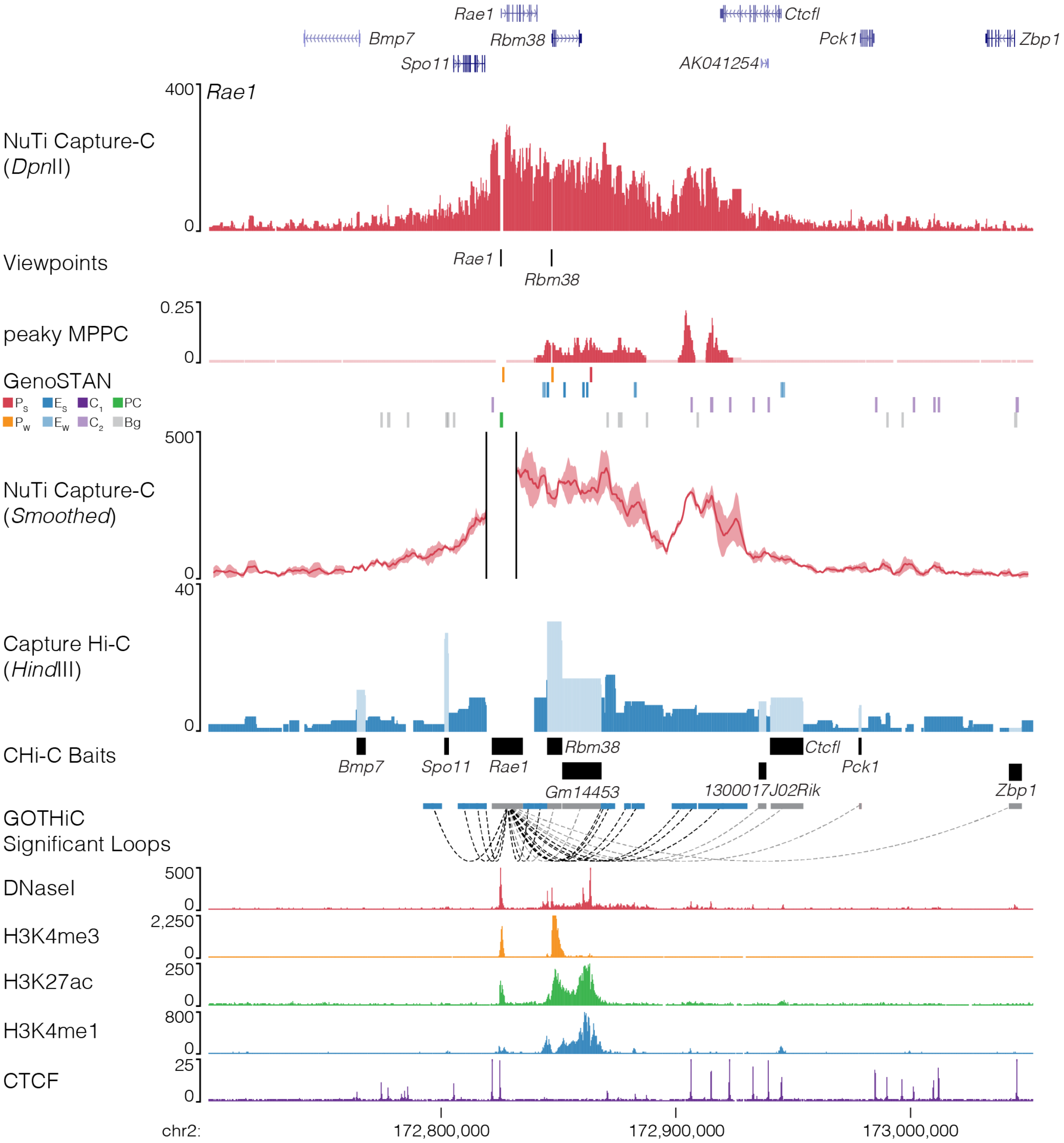


**Fig. 23 | NuTi Capture-C from the *Rae1* promoter.** Sequence tracks showing the difference between high-resolution 3C (*Dpn*II, NuTi Capture-C) and low-resolution 3C (*Hind*III, Capture Hi-C) at calling interacting fragments (mm9, chr2:172,701,139-173,052,278) in erythroid cells. Tracks in order: UCSC gene annotation, *cis*-normalized mean interactions per *Dpn*II fragment using NuTi Capture-C (n=3), NuTi Capture-C viewpoints, peaky Marginal Posterior Probability of Contact (MPPC) scores with fragments with MPPC ≥0.01 darker, GenoSTAN open chromatin classification, windowed mean interactions using NuTi Capture-C, total supporting reads per *Hind*III fragment with CHi-C (n=2; co-targeted fragments are lighter in colour), CHi-C bait fragments, loops between reported significantly interacting fragments (co-targeting loops are coloured grey), erythroid tracks for open chromatin (DNaseI), promoters (H3K4me3), active transcription (H3K27ac), enhancers (H3K4me1), and boundaries (CTCF). Note overlapping MPPC signals appear darker in colour.

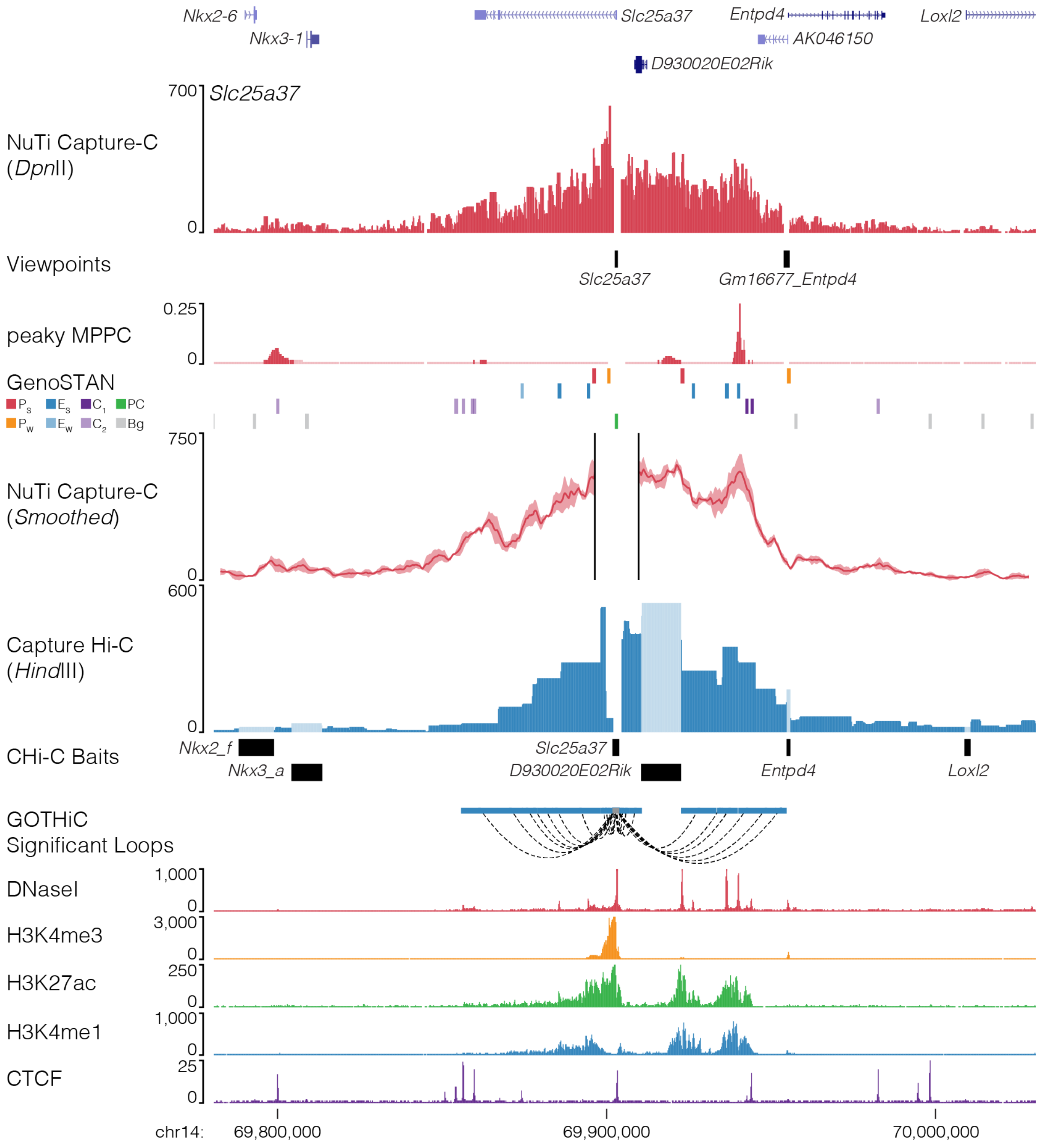


**Fig. 24 | NuTi Capture-C from the *Slc25a37* promoter.** Sequence tracks showing the difference between high-resolution 3C (*Dpn*II, NuTi Capture-C) and low-resolution 3C (*Hind*III, Capture Hi-C) at calling interacting fragments (mm9, chr14:69,780,624-70,030,623) in erythroid cells. Tracks in order: UCSC gene annotation, *cis*-normalized mean interactions per *Dpn*II fragment using NuTi Capture-C (n=3), NuTi Capture-C viewpoints, peaky Marginal Posterior Probability of Contact (MPPC) scores with fragments with MPPC ≥0.01 darker, GenoSTAN open chromatin classification, windowed mean interactions using NuTi Capture-C, total supporting reads per *Hind*III fragment with CHi-C (n=2; co-targeted fragments are lighter in colour), CHi-C bait fragments, loops between reported significantly interacting fragments (co-targeting loops are coloured grey), erythroid tracks for open chromatin (DNaseI), promoters (H3K4me3), active transcription (H3K27ac), enhancers (H3K4me1), and boundaries (CTCF). Note overlapping MPPC signals appear darker in colour.

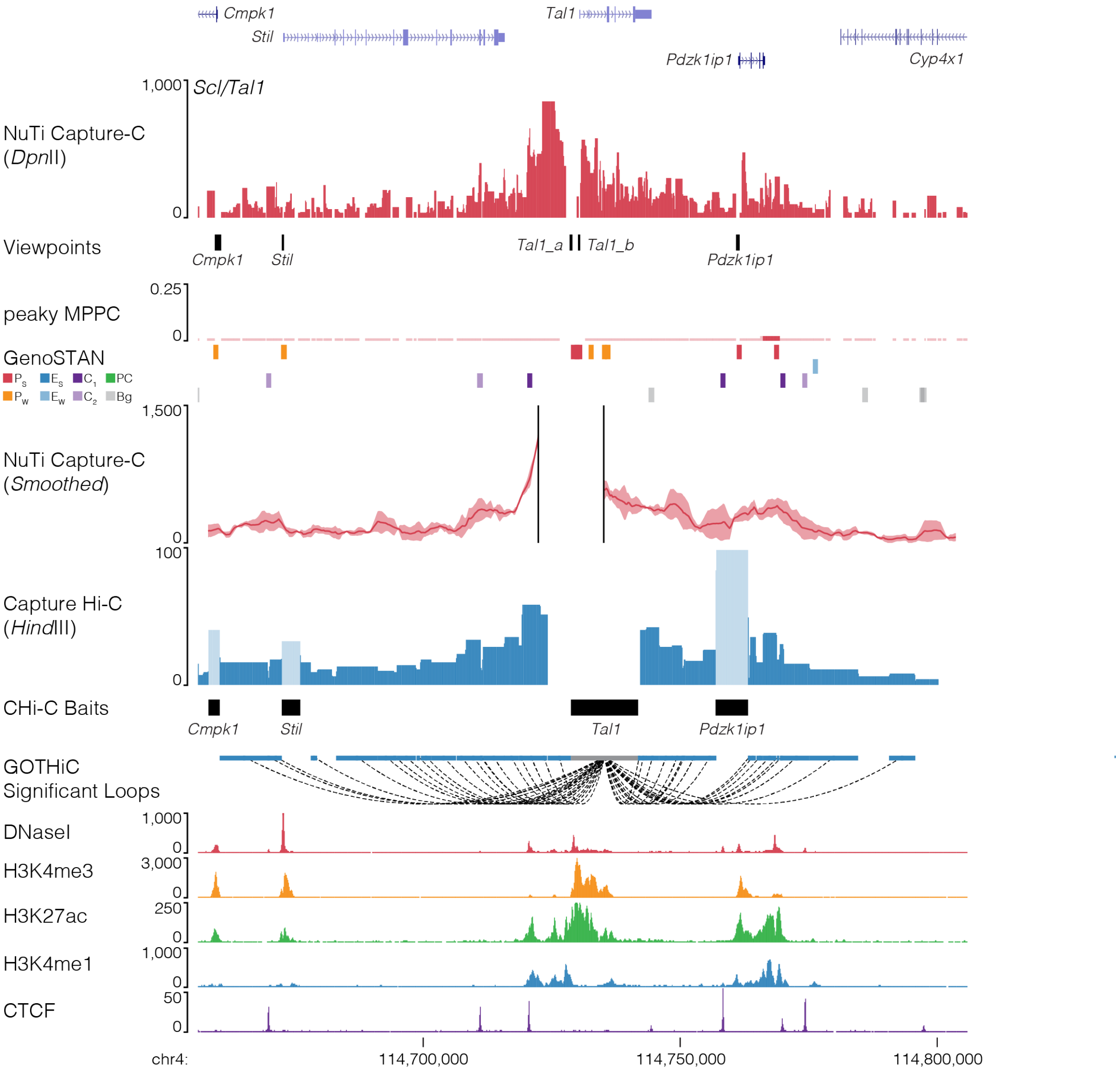


**Fig. 25 | NuTi Capture-C from the *Tal1* promoter.** Sequence tracks showing the difference between high-resolution 3C (*Dpn*II, NuTi Capture-C) and low-resolution 3C (*Hind*III, Capture Hi-C) at calling interacting fragments (mm9, chr4:114,656,021-114,806,020) in erythroid cells. Tracks in order: UCSC gene annotation, *cis*-normalized mean interactions per *Dpn*II fragment using NuTi Capture-C (n=3), NuTi Capture-C viewpoints, peaky Marginal Posterior Probability of Contact (MPPC) scores with fragments with MPPC ≥0.01 darker, GenoSTAN open chromatin classification, windowed mean interactions using NuTi Capture-C, total supporting reads per *Hind*III fragment with CHi-C (n=2; co-targeted fragments are lighter in colour), CHi-C bait fragments, loops between reported significantly interacting fragments (co-targeting loops are coloured grey), erythroid tracks for open chromatin (DNaseI), promoters (H3K4me3), active transcription (H3K27ac), enhancers (H3K4me1), and boundaries (CTCF). Note overlapping MPPC signals appear darker in colour.

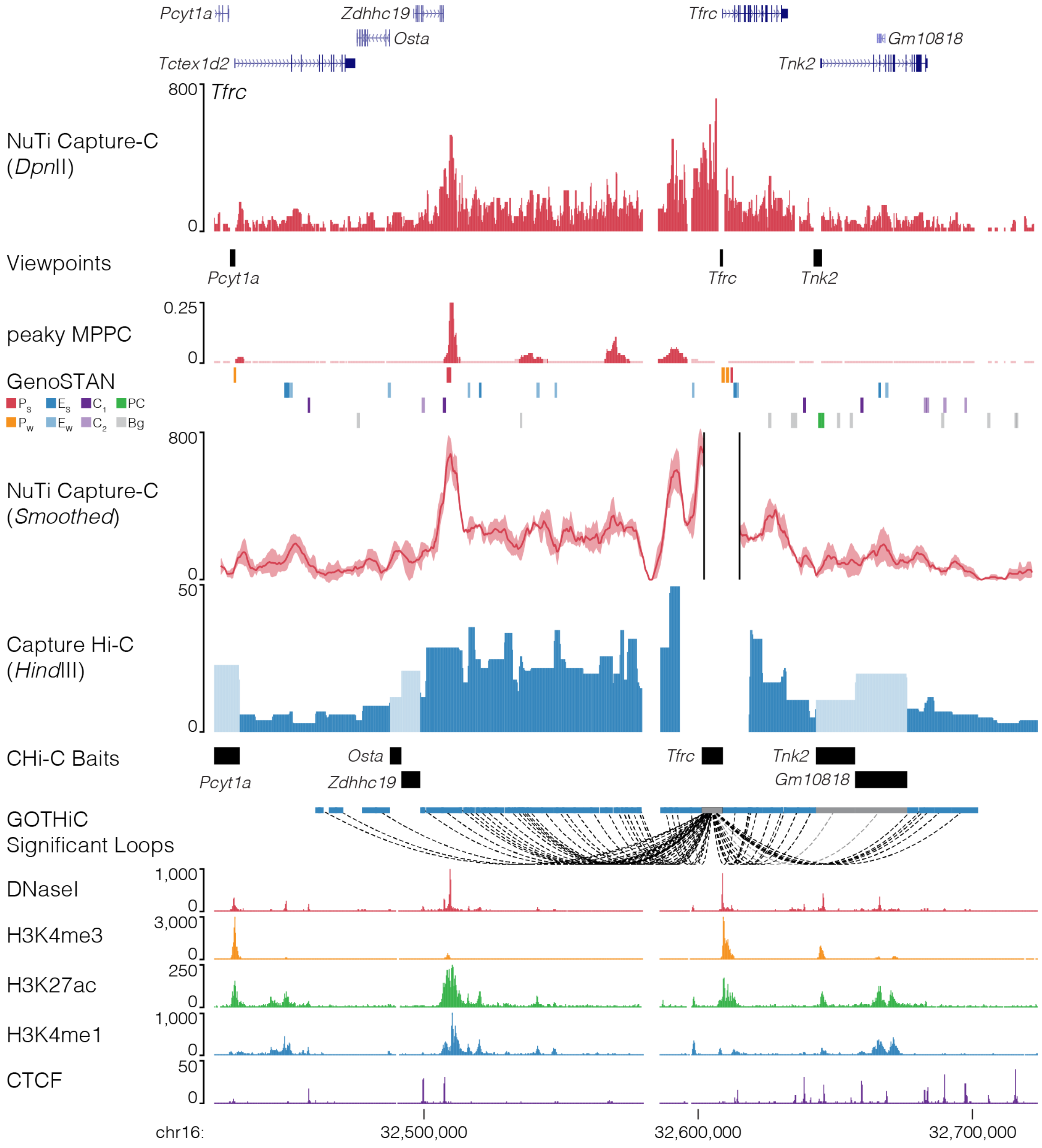


**Fig. 26 | NuTi Capture-C from the *Tfrc* promoter.** Sequence tracks showing the difference between high-resolution 3C (*Dpn*II, NuTi Capture-C) and low-resolution 3C (*Hind*III, Capture Hi-C) at calling interacting fragments (mm9, chr16:32,423,792-32,723,792) in erythroid cells. Tracks in order: UCSC gene annotation, *cis*-normalized mean interactions per *Dpn*II fragment using NuTi Capture-C (n=3), NuTi Capture-C viewpoints, peaky Marginal Posterior Probability of Contact (MPPC) scores with fragments with MPPC ≥0.01 darker, GenoSTAN open chromatin classification, windowed mean interactions using NuTi Capture-C, total supporting reads per *Hind*III fragment with CHi-C (n=2; co-targeted fragments are lighter in colour), CHi-C bait fragments, loops between reported significantly interacting fragments (co-targeting loops are coloured grey), erythroid tracks for open chromatin (DNaseI), promoters (H3K4me3), active transcription (H3K27ac), enhancers (H3K4me1), and boundaries (CTCF). Note overlapping MPPC signals appear darker in colour.

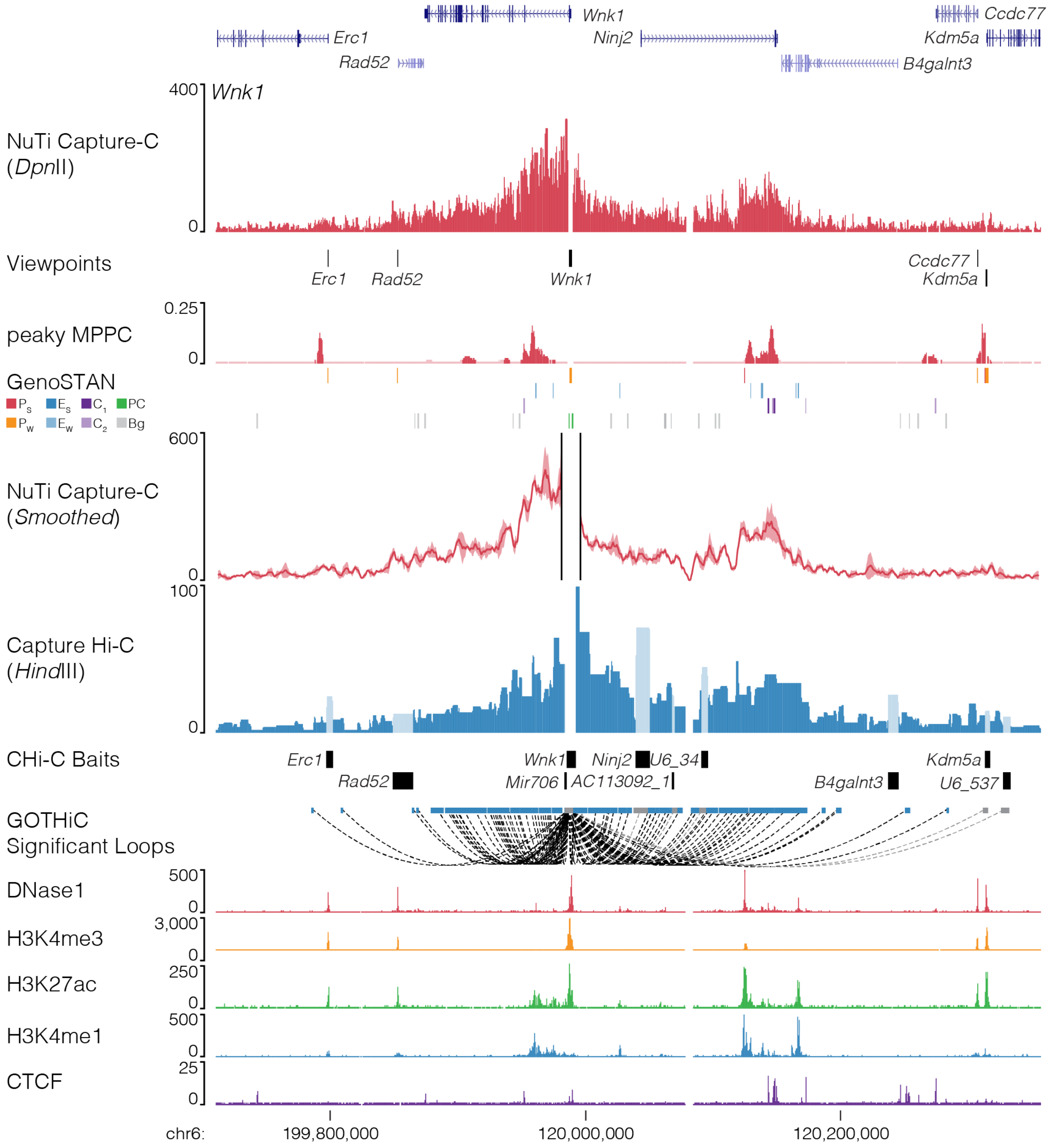


**Fig. 27 | NuTi Capture-C from the *Wnk1* promoter.** Sequence tracks showing the difference between high-resolution 3C (*Dpn*II, NuTi Capture-C) and low-resolution 3C (*Hind*III, Capture Hi-C) at calling interacting fragments (mm9, chr6:119,710,118-120,356,868) in erythroid cells. Tracks in order: UCSC gene annotation, *cis*-normalized mean interactions per *Dpn*II fragment using NuTi Capture-C (n=3), NuTi Capture-C viewpoints, peaky Marginal Posterior Probability of Contact (MPPC) scores with fragments with MPPC ≥0.01 darker, GenoSTAN open chromatin classification, windowed mean interactions using NuTi Capture-C, total supporting reads per *Hind*III fragment with CHi-C (n=2; co-targeted fragments are lighter in colour), CHi-C bait fragments, loops between reported significantly interacting fragments (co-targeting loops are coloured grey), erythroid tracks for open chromatin (DNaseI), promoters (H3K4me3), active transcription (H3K27ac), enhancers (H3K4me1), and boundaries (CTCF). Note overlapping MPPC signals appear darker in colour.

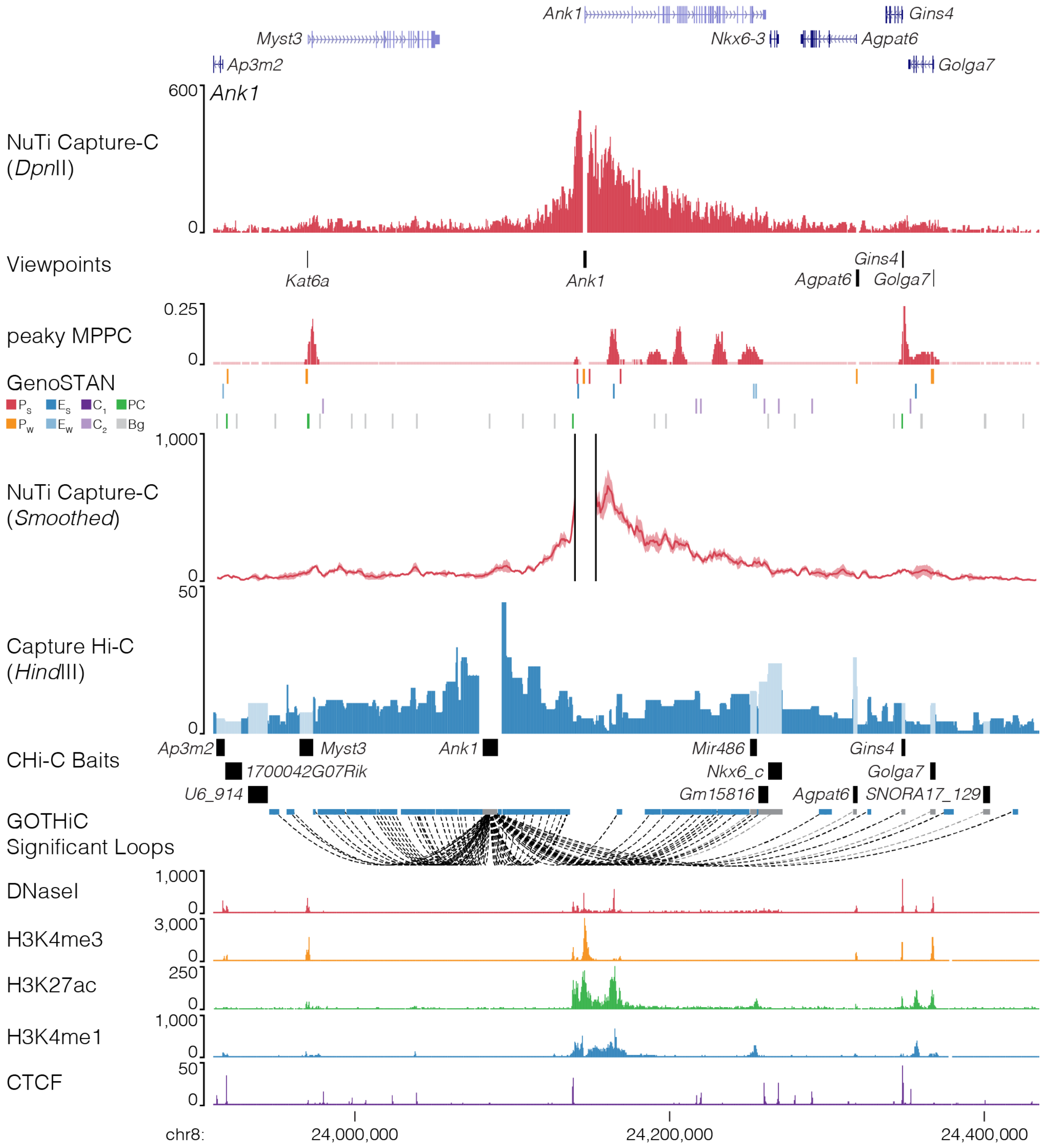


**Fig. 28 | NuTi Capture-C from the *Ank1* promoter.** Sequence tracks showing the importance of tissue specific probe design when performing promoter capture with either high-resolution 3C (*Dpn*II, NuTi Capture-C) or low-resolution 3C (*Hind*III, Capture Hi-C), particularly for genes with multiple promoters (mm9, chr8:23,910,000-24,435,000) in erythroid cells. Tracks in order: UCSC gene annotation, *cis*-normalized mean interactions per *Dpn*II fragment using NuTi Capture-C (n=3), NuTi Capture-C viewpoints, peaky Marginal Posterior Probability of Contact (MPPC) scores with fragments with MPPC ≥0.01 darker, GenoSTAN open chromatin classification, windowed mean interactions using NuTi Capture-C, total supporting reads per *Hind*III fragment with CHi-C (n=2; co-targeted fragments are lighter in colour), CHi-C bait fragments, loops between reported significantly interacting fragments (co-targeting loops are coloured grey), erythroid tracks for open chromatin (DNaseI), promoters (H3K4me3), active transcription (H3K27ac), enhancers (H3K4me1), and boundaries (CTCF). Note overlapping MPPC signals appear darker in colour.

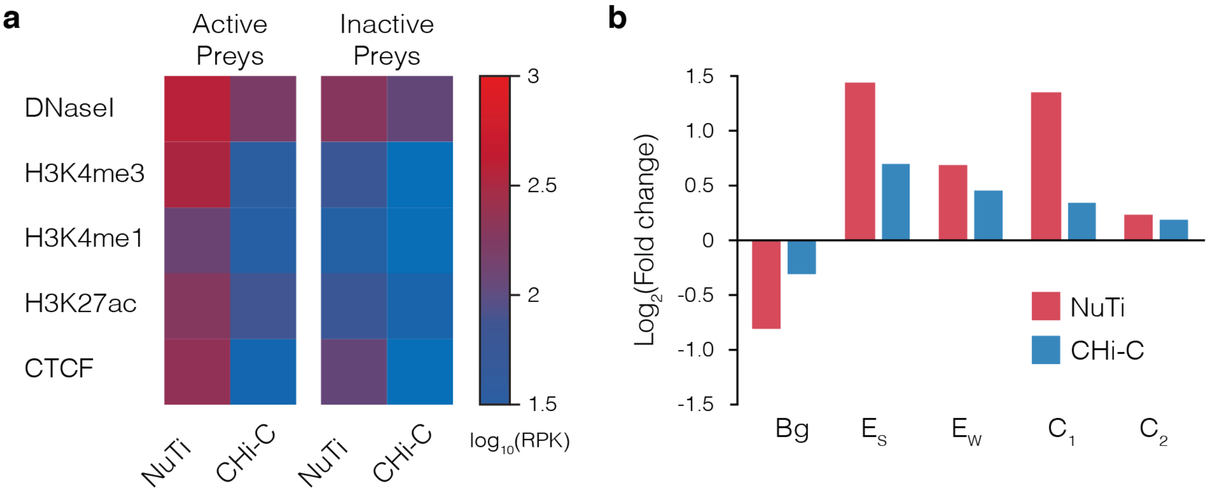


**Supp. Fig. 29 | Comparison of interactions identified by NuTi Capture-C and Capture Hi-C. a,** Average chromatin signature in mouse erythroid cells over fragments identified as being significantly interacting with either active or inactive promoters by NuTi Capture-C (Nu-3C) and Capture Hi-C (CHi-C). **b,** Enrichment of different classes of open chromatin element in fragments identified as being significantly interacting with active promoters. $E_S$: Enhancer (Strong H3K27ac), $E_W$: Enhancer (Weak H3K27ac), $C_1$: CTCF near promoter/enhancer, $C_2$: CTCF, Bg: Background, RPK: Reads per kilobase.
