## Supplementary Note for "Targeted high-resolution chromosome conformation capture at genome-wide scale"

**SUPPLEMENTARY NOTE:** *Mathematical modelling of 3C enrichment bias.*

The number of interactions in 3C experiments are constrained by the fact each fragment can only ligate to two other fragments. Therefore, the total number of interactions is limited by the total number of cells, in effect, it is a closed system. To explore the effects of enrichment for multiple targets on 3C libraries we created a small closed system of fragments where each fragment has 5,000 interactions. Within this system, interactions involving three fragments, “A”, “B”, and “C”, can be sampled (i.e. enriched) with varying levels of efficiency. The remaining fragments can be collectively considered “X”, with “D” being one of these remaining fragments. The absolute number of interactions between each fragment within this system and example interaction profiles are demonstrated below (Supp. Note Table 1, Supp. Note Fig. 1). To demonstrate the effect on interaction calling, within this system “significant interactions” are simply those observed at a frequency of greater than 1 in 50 (>0.02). The significant interactions in this system are A-B, A-D, and B-D.

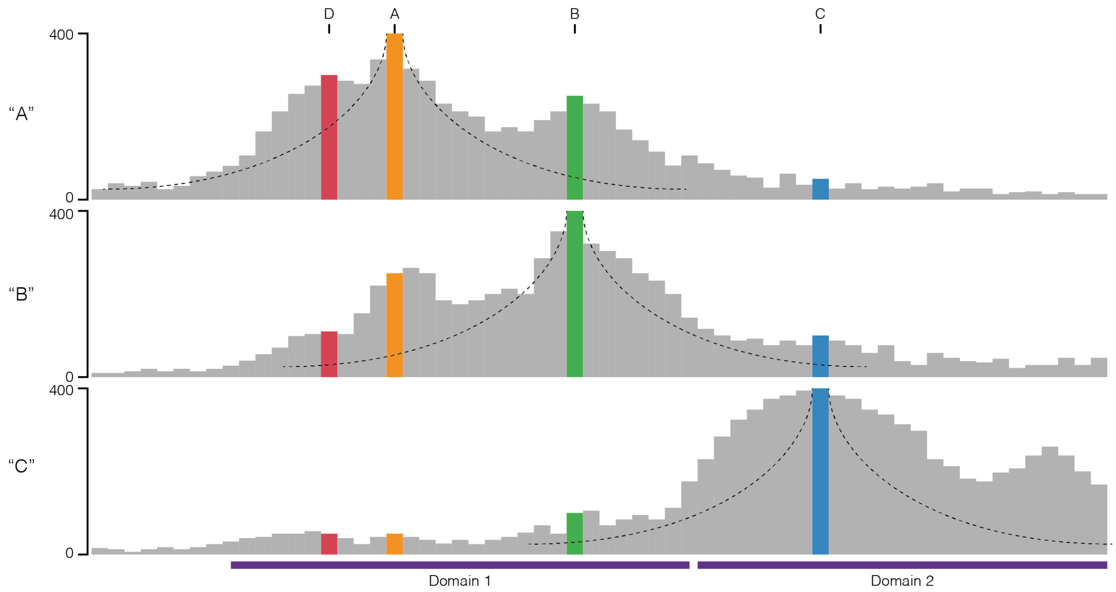


**Supp. Note Fig. 1 I A closed system of interactions.** Profile of absolute interaction counts for three sampleable fragments “A”, “B” and “C” within a closed system. The fragments separate into two interacting domains. Monotonic decay curves associated with the polymer models of interaction are shown.

**Supp. Note Table 1.** A closed system of interactions.

| | | Interacting Fragment / Prey | | | | | | | |
|---|---|---|---|---|---|---|---|---|---|
| | | Real Count | | | | Real Frequency | | | |
| | | “A” | “B” | “C” | “D” | “A” | “B” | “C” | “D” |
| Viewpoint / Bait (Total count) | “A” (5,000) | – | 250 | 50 | 300 | – | **0.0500** | 0.0100 | **0.0600** |
| | “B” (5,000) | 250 | – | 100 | 110 | **0.0500** | – | 0.0200 | **0.0220** |
| | “C” (5,000) | 50 | 100 | – | 50 | 0.0100 | 0.0200 | – | 0.0100 |

*Significant interactions (Freq. > 0.02) are shaded green.

The sampling within this system represents the targeted enrichment of 3C methods (e.g. probe hybridisation or immunoprecipitation). For 3C enrichment, efficiency can be affected by, among other things, the number of individual probes targeting each viewpoint and the melting points (e.g. Capture-C, CHi-C), and the level of target signal (e.g. Hi-ChIP, Hi-ChIRP, ChIA-PET). To represent this diversity in these processes, we assigned sampling of “A” and “C” to be highly efficient at 80% and 90% respectively. Whereas enrichment of “B” is relatively low at 10%. When each of these sampling efficiencies is applied to any one fragment at time, the total number of observed interactions decreases, however the frequency of interaction within the system remains constant (Note Fig. 2, Note Table 2). At this point, it is important to note that while the B-to-C count is lower than the C-to-B count, their proportional frequencies are still equal, and the same set of “significantly interacting” fragments are detected.

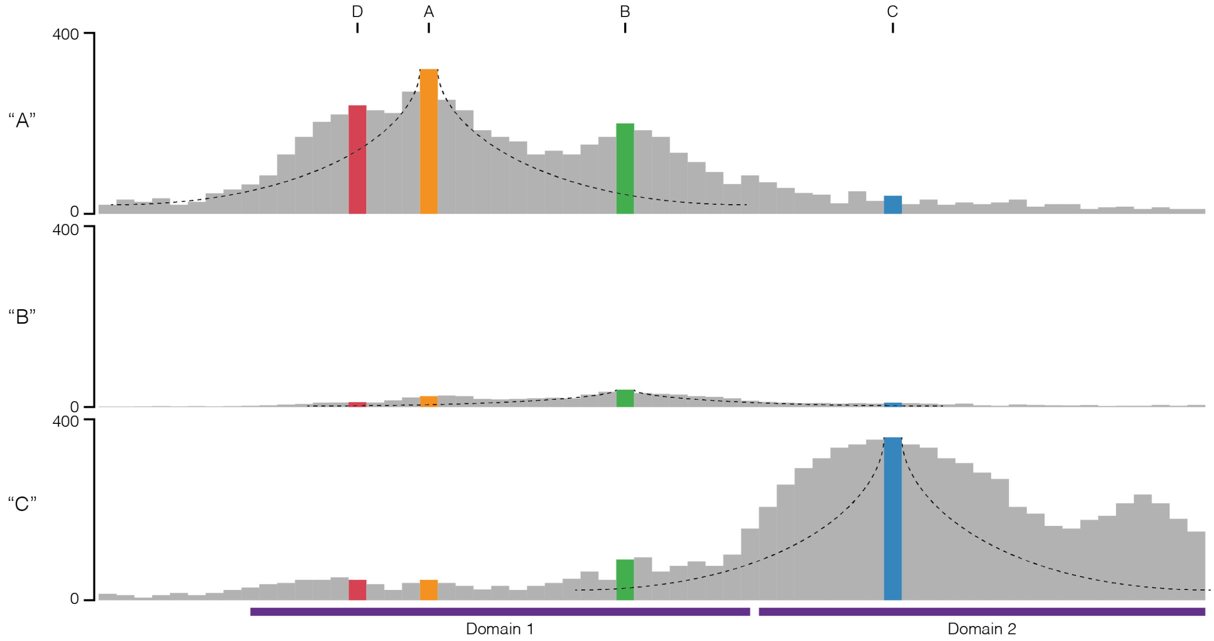


**Supp. Note Fig. 2 I Independent sampling of interactions.** Profile of observed interaction counts for three independently sampled fragments “A”, “B” and “C” within a closed system.

**Supp. Note Table 2.** Interaction counts following independent sampling.

| | | Interacting Fragment (Prey) | | | | | | | | | | | |
|---|---|---|---|---|---|---|---|---|---|---|---|---|---|
| | | Observed Count | | | | Observed Frequency | | | | Observed Freq. / Real Freq. | | | |
| | | “A” | “B” | “C” | “D” | “A” | “B” | “C” | “D” | “A” | “B” | “C” | “D” |
| Viewpoint / Bait (Total count) | “A” (4,000) | – | 200 | 40 | 240 | – | **0.0500** | 0.0100 | **0.0600** | – | 1 | 1 | 1 |
| | “B” (500) | 25 | – | 10 | 11 | **0.0500** | – | 0.0200 | **0.0220** | 1 | – | 1 | 1 |
| | “C” (4,500) | 45 | 90 | – | 45 | 0.0100 | 0.0200 | – | 0.0100 | 1 | 1 | – | 1 |

*Significant interactions (Freq. > 0.02) are shaded green.

When we consider sampling fragment “A”, at 80% efficiency, we fail to see 50 “A-B” interactions, ten “A-C” interactions, and 60 “A-D” interactions. If we were to simultaneously sample (co-sample) the remaining viewpoints, from the missed interactions would can recover five “A-B” interactions (at 10% “B” sampling efficiency), nine “A-C” interactions (at 90% “B” sampling efficiency), and zero “A-D” interactions as it is unsampled. This recovery can be applied to each fragment as the first or second fragment sampled and leads to as much as a 9.1-fold increase in observed interaction counts. When the co-sampled fragments are presented as frequency values significant divergence from the true values within the system is observed (Supp. Note Fig. 3, Supp. Note Table 3). This divergence in frequency varies across each interacting pair and ranges from a 0.65-fold decrease (B-D) to a near 6-fold increase (B-C).

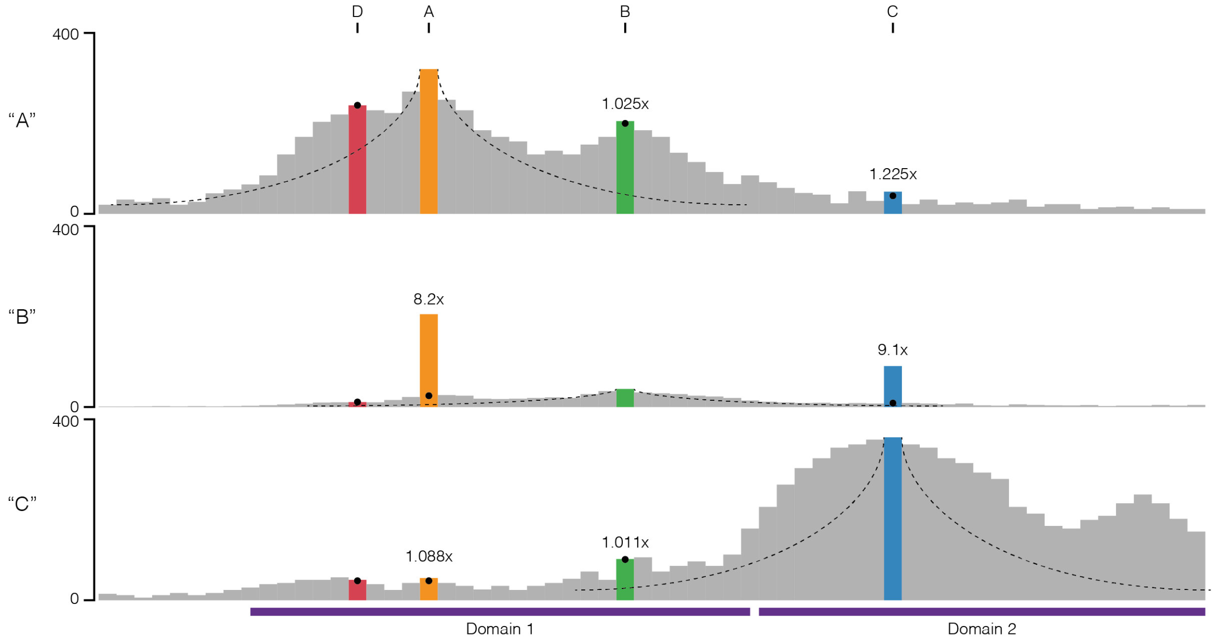


**Supp. Note Fig. 3 l Co-sampling of interactions.** Profile of observed interaction counts for three co-sampled fragments “A”, “B” and “C” within a closed system. Black circles represent counts observed with independent sampling, and the observed fold difference in raw counts is shown.

**Note Table 3.** Interaction counts following co-sampling.

| | | Interacting Fragment (Prey) | | | | | | | | | | | |
|---|---|---|---|---|---|---|---|---|---|---|---|---|---|
| | | Observed Count | | | | Observed Frequency | | | | Observed Freq. / Real Freq. | | | |
| | | “A” | “B” | “C” | “D” | “A” | “B” | “C” | “D” | “A” | “B” | “C” | “D” |
| Viewpoint / Bait (Total count) | “A” (4,014) | – | 205 | 49 | 240 | – | **0.0511** | 0.0122 | **0.0598** | – | 1.021 | 1.221 | 0.997 |
| | “B” (761) | 205 | – | 91 | 11 | **0.2694** | – | **0.1196** | 0.0145 | 5.387 | – | **5.978** | **0.657** |
| | “C” (4,505) | 49 | 91 | – | 45 | 0.0109 | **0.0202** | – | 0.0100 | 1.087 | 1.009 | – | 0.998 |

*Significant interactions (Freq. > 0.02) are shaded green.

Interestingly, where previously the frequency of B-to-C matched C-to-B, it is now the count of interactions which is equal, and the observed frequency is unequal. This change in frequency also results in different significant interactions, B-D is no longer called (false negative), while B-C is now called (false positive). The effect on relative frequency of co-sampling is bi-directional: the observed frequency of interaction with co-sampled fragments increases, and the observed frequency of interaction with un-sampled fragments decreases. Therefore, significant and variable bias is introduced by co-targeting.

To determine the effect of this divergence we formalized this bias into a polynomial equation (Supp. Note Fig. 4) describing the observed interaction frequency of two co-targeted fragments ($O_{AB}$) within all interactions containing "A". For the numerator, the efficiency of targeting "A" ($E_A$) and the real frequency of "A-B" ($f_{AB}$) determines the number of interactions sampled by the A probe and the number of "lost interactions" available for the B probe (1- $E_A$). These lost interactions are then recovered at the efficiency of the B probe ($E_B$). The denominator, which describes the total observations involving A, can be simply denoted as $E_A$ plus the number of recovered events ($E_B$ x $f_{AB}$ x [1- $E_A$]). The level of bias can then be calculated as $O_{AB}$ divided by $f_{AB}$.

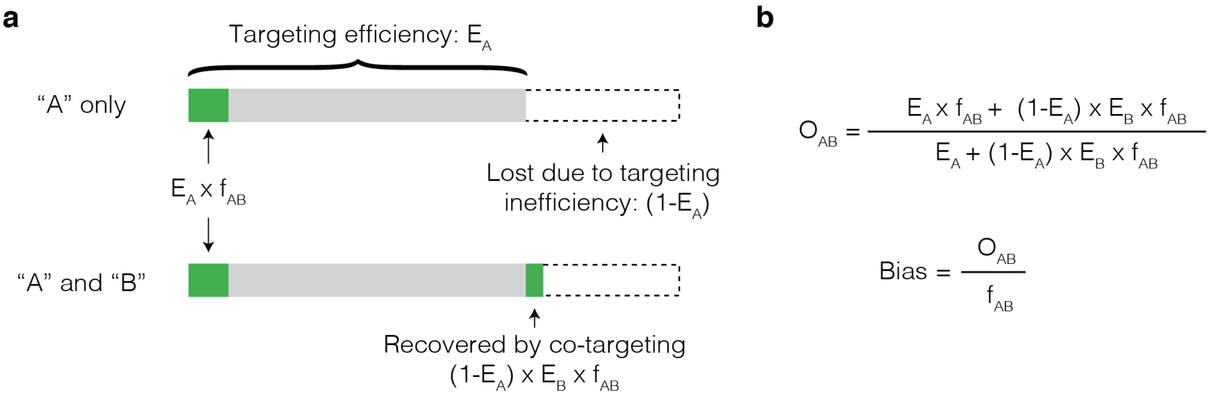


**Supp. Note Fig. 4 I Model for the effect on observed frequency caused by co-targeting. a,** Diagram of the total number of interactions containing A (entire bar), which includes A-B (green) and the effect of incompletely efficient targeting. Un-enriched (or lost) interactions are in dotted lines, but can be recovered by capture with additional probes. **b,** Equations for the observed frequency of A-B interaction ($O_{AB}$) when both A and B are samples and calculation of bias.

We used this equation to model the effects of variable efficiency of enrichment (0.05-1.0), and variable underlying interaction frequencies (1/200,000, 1/25,000 and 1/40; based on the min, median, and max interactions frequencies associated with *Hba-1* capture). Under these tested parameters the highest level of bias was a ~20-fold increase in frequency (Supp. Note Fig. 5), seen when the primary target had a low enrichment efficiency, and the secondary target had a high enrichment efficiency. Notably, for any given level of enrichment efficiency the level of bias was variable across the interaction frequencies, with infrequent interactions more affected then frequent interactions.

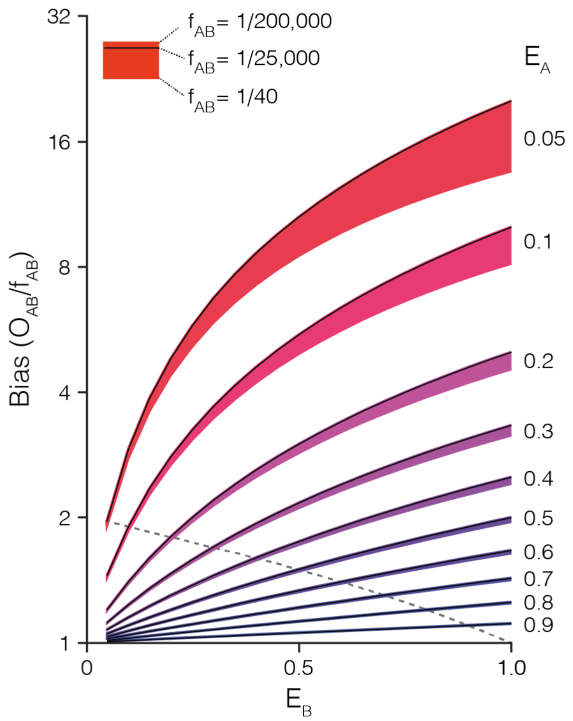


**Supp. Note Fig. 5 I Levels of observed frequency bias caused by co-targeting.** Variable levels of bias are observed in the observed frequency of A-B interaction ($O_{AB}$) when altering the efficiency of targeting fragment A ($E_A$), targeting fragment B ($E_B$), and the real frequency of A-B interaction ($f_{AB}$). The Dashed line shows when efficiency of targeting A and B is equal ($E_A = E_B$). Note that bias is avoided only when $E_A$ is equal to either zero or one.

The implication of these results are quite striking and two-fold. Firstly, when investigating a viewpoint with very low enrichment (say ChIA-PET for a poorly bound PolII site, or Hi-ChIP at a weak H3K27ac peak) then significant enrichment bias is likely to be seen at strong PolII or H3K27ac sites, regardless of whether or not they are actually interacting. In fact, the rarer an interaction is, the stronger the bias effect. Secondly, because all three parameters (enrichment at targets, and the underlying interaction frequency) contribute significantly to the observed bias, proper data correction depends upon having accurate values for all three parameters.
